# Helotiales fungi as potential nutritional partners for non-mycorrhizal plants: a machine learning and experimental approach

**DOI:** 10.64898/2026.03.23.710836

**Authors:** Pauline Bruyant, Lauren M. Gillespie, Jeanne Doré, Pierre Emmanuel Courty, Yvan Moënne-Loccoz, Juliana Almario

## Abstract

**Background:** Most land plants depend on the ancestral arbuscular mycorrhizal (AM) symbiosis for phosphorus (P) acquisition. However, several plant lineages have independently lost this symbiosis, raising fundamental questions about how these ‘non-mycorrhizal’ plants meet their nutritional requirements without this crucial partnership.

**Results:** Comparative genomic analyses confirmed that Cyperaceae, Caryophyllaceae, and Brassicaceae lack genes essential for AM symbiosis, indicating that these lineages independently abandoned this association 90–122 million years ago. Field surveys of 42 wild populations across seven sites revealed that while non-mycorrhizal plants generally maintain shoot P levels comparable to those in AM neighbors, lower shoot P levels can be observed in low P soils.

To identify fungal taxa potentially associated with P nutrition in non-mycorrhizal plants, we applied a machine-learning approach to predict plant P-accumulation from root microbiome composition. The model explained substantial variance in plant P-accumulation (57-69%), and identified 85 fungal taxa as key predictors of shoot P-accumulation, predominantly belonging to the Helotiales (28%) and Pleosporales (23%) orders.

Experimental validation of two phylogenetically distant Helotiales lineages (*Tetracladium maxilliforme* OTU29 and Helotiales sp. OTU7), using isotopic tracing, demonstrated their capacity to enhance plant growth and transfer P (and N) to their native non-mycorrhizal hosts under P-limiting conditions.

**Conclusions:** Our findings suggest that non-mycorrhizal plants engage in nutritional partnerships with diverse Helotiales lineages that could collectively contribute to their mineral nutrition. However, given the widespread distribution of these Helotiales fungi, including in roots of AM plants, they may play a broader role in plant nutrition, i.e. also in mycorrhizal hosts. This study provides proof of concept for a novel framework integrating machine-learning predictions with experimental validation to identify functionally important microbial partnerships in natural plant communities.

## Background

Phosphorus (P) is essential for plant growth, and its low availability is rapidly becoming the main nutritional constrain limiting plant growth and productivity across terrestrial ecosystems [1,2]. Plants can only absorb inorganic P (Pi) as soluble phosphate, a minor soil P-form with low diffusion capacity [3]. However, root-associated microorganisms can support plant Pi acquisition through processes like Pi solubilization and organic-P mineralization [3,4]. Arbuscular mycorrhizal (AM) Glomeromycota fungi can provide up to 100% of a plant’s P needs [3,5,6] by directly transferring soil-derived P that is taken up by the fungus, transported over long hyphal distances, and delivered to plant cells via specialized structures called arbuscules [7,8].

The AM symbiosis is thought to have facilitated plant adaptation to terrestrial environments 450 MYA [9] and is conserved in the large majority of extant plant species, encompassing 71% of vascular plants [10]. Surprisingly, ancestral state reconstruction analyses hint to at least 25 events of AM symbiosis loss throughout plant evolution [11]. While these losses may correspond to non-microbial nutritional adaptations, such as plant parasitism or carnivory, many correspond to symbiont-switching events where non-AM plant lineages establish evolutionarily younger mycorrhizal symbioses with non-Glomeromycota fungi [11]. This is the case of the ectomycorrhizal symbiosis with Basidiomycota Agaricomycetes in non-AM Pinaceae or the ericoid mycorrhizal symbiosis with Ascomycota Helotiales in non-AM Ericaceae [11,12]. Similar symbiont-switching has been observed across a wide range of biological systems, suggesting it is a common adaptive response to environmental change [13].

There are however three major non-mycorrhizal plant families thought to have lost the AM symbiosis with no alternative nutritional strategy described to date (Brassicaceae, Caryophyllaceae, and Cyperaceae) [11], bringing to question whether and by which mechanisms these plants meet their P needs, especially in P-limited soils. Although the non-AM status of Brassicaceae is confirmed by the loss of most AM-essential genes [14], we still lack genomic evidence to establish the non-AM status of Cyperaceae and Caryophyllaceae.

The microbiota can play crucial roles by mediating host adaptation [15], including in plants [16,17]. While little is known about the root microbiota of Cyperaceae and Caryophyllaceae, previous work on Brassicaceae uncovered ‘mycorrhizal-like’ fungal isolates from the Glomerellales and Helotiales orders that transfer P to their hosts without forming specific symbiotic cellular structures inside plant roots [18–20]. However, the broader biological and ecological significance of these interactions remains uncertain, as research on the root microbiota of cultivated and model non-AM plants has failed to provide evidence of a major role for these fungi [16]. It is thus unclear how widespread these associations are in nature – among diverse non-AM hosts and among diverse root-colonizing fungi – and how important they actually are for plant nutrition. Here, we hypothesize that non-mycorrhizal plants rely on novel nutritional associations with diverse fungal partners from their root microbiota, contributing to their P nutrition notably via mechanisms of direct nutrient transfer.

Plants tissues harbor diverse microbial communities comprising thousands of fungal and bacterial species. The inherent complexity and variability of these communities poses a significant challenge for identifying microbial groups with preponderant impacts on plant growth. Large-scale machine-learning approaches linking microbiome diversity to plant traits in agricultural settings have shown the predictive value of microbiome data for forecasting plant disease outcome [21], plant yield [22], or plant vigor [23]. While these studies have pinpointed microbial groups with the potential to modulate plant traits of interest, few have experimentally validated their predictions [24]. Moreover, such approaches remain underutilized in the context of natural plant communities and microbiome-mediated plant nutrient acquisition.

The aim of this study was to dissect the fungal root microbiota of wild-growing non-mycorrhizal plants to uncover microbiota-based mechanism underpinning plant nutrition. Since AM and non-AM plant lineages do not coexist within the same plant family, our approach was to compare closely related families that differ in their AM status but share the same environments.

After confirming the non-mycorrhizal status of our focal plant families using genomics, we leveraged natural variation in microbiome and P-accumulation across plant individuals, species, and families, to identify fungal taxa associated with higher plant P-uptake using a machine-learning approach. Two Helotiales lineages were identified as key taxa and experimentally validated through plant inoculation and nutrient transfer assays. Our results support the hypothesis that non-mycorrhizal plant lineages rely on diverse Helotiales associations that collectively contribute to their mineral nutrition. However, the observed association of these Helotiales with AM plants suggests that they could play a broader role in plant nutrition beyond the case of non-AM hosts.

## Results

### Convergent gene losses in non-mycorrhizal plant families

We examined the presence of five genes essential for the establishment of the AM symbiosis (SYMRK/DMI2, leucine-rich repeat receptor kinase; CCAMK/DMI3, calcium and calmodulin-dependent protein kinase; CYCLOPS/IPD3, transcription factor) or the symbiotic transfer of lipids from the plant to the AM fungus (STR1 and STR2, half-size ABCG transporters) in the genomes of known AM plant families as well as confirmed (Brassicaceae) and putative non-AM families (Caryophyllaceae and Cyperaceae) [11]. Putative orthologues of STR1, STR2, SYMRK, CCAMK, and CYCLOPS were detected in all 11 AM families studied (Figs. S1-4; Table S1) but none in the three known or suspected non-AM families screened. These orthologues were also absent from sister lineages of Caryophyllaceae (i.e. Amaranthaceae, Polygonaceae, Droseraceae, and Cactaceae) and Cyperaceae (i.e. Juncaceae) (Fig. 1). These results point to the independent emergence of three major non-AM lineages through convergent losses of AM-symbiotic genes in the common ancestors of (i) the Brassicaceae family, (ii) the Caryophyllales clade, and (iii) the Juncaceae-Cyperaceae families. Because AM and non-AM lineages do not coexist within the same plant family, we focused our study on three distinct non-AM families (Brassicaceae, Caryophyllaceae, and Cyperaceae), comparing them with co-occurring AM plants from closely related families: Ranunculaceae and Geraniaceae, Asteraceae, and Poaceae, respectively (Fig. 1-2).

**Figure 1.**
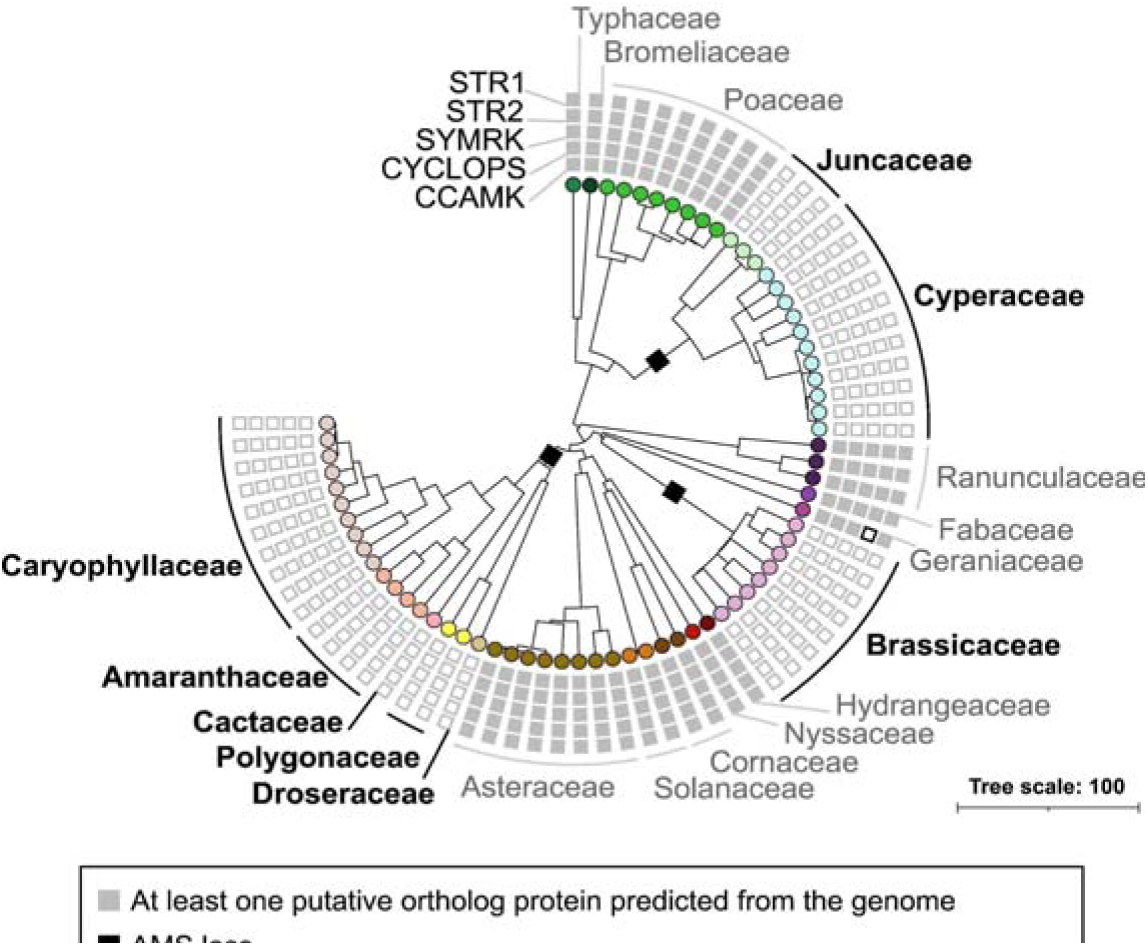
Distribution of genes encoding AM-symbiotic proteins in AM and non-AM plant families Cyperaceae, Brassicaceae, and Caryophyllaceae. Plant genomes of interest available in NCBI (67) were screened for the five major AM symbiotic genes SYMRK, CCAMK, CYCLOPS, STR1, and STR2 using reference sequences from AM model species *Medicago truncatula* and *Oryza sativa* [25]. The deduced protein sequences were used to build phylogenetic trees and identify putative orthologs. In *Geranium maculatum* (Geraniaceae), no STR2 ortholog was found, but a putative ortholog with a Val start codon was predicted (indicated by a black outlined square). AM symbiosis loss events are indicated by black squares on tree branches. The plant phylogeny tree was produced using PhyloMaker [26] and the plant megatree. Specific colors are used to distinguish plant families. Mycorrhizal lineages are indicated in grey and non-AM lineages in black. The scale represents the number of differences between sequences.

**Figure 2.**
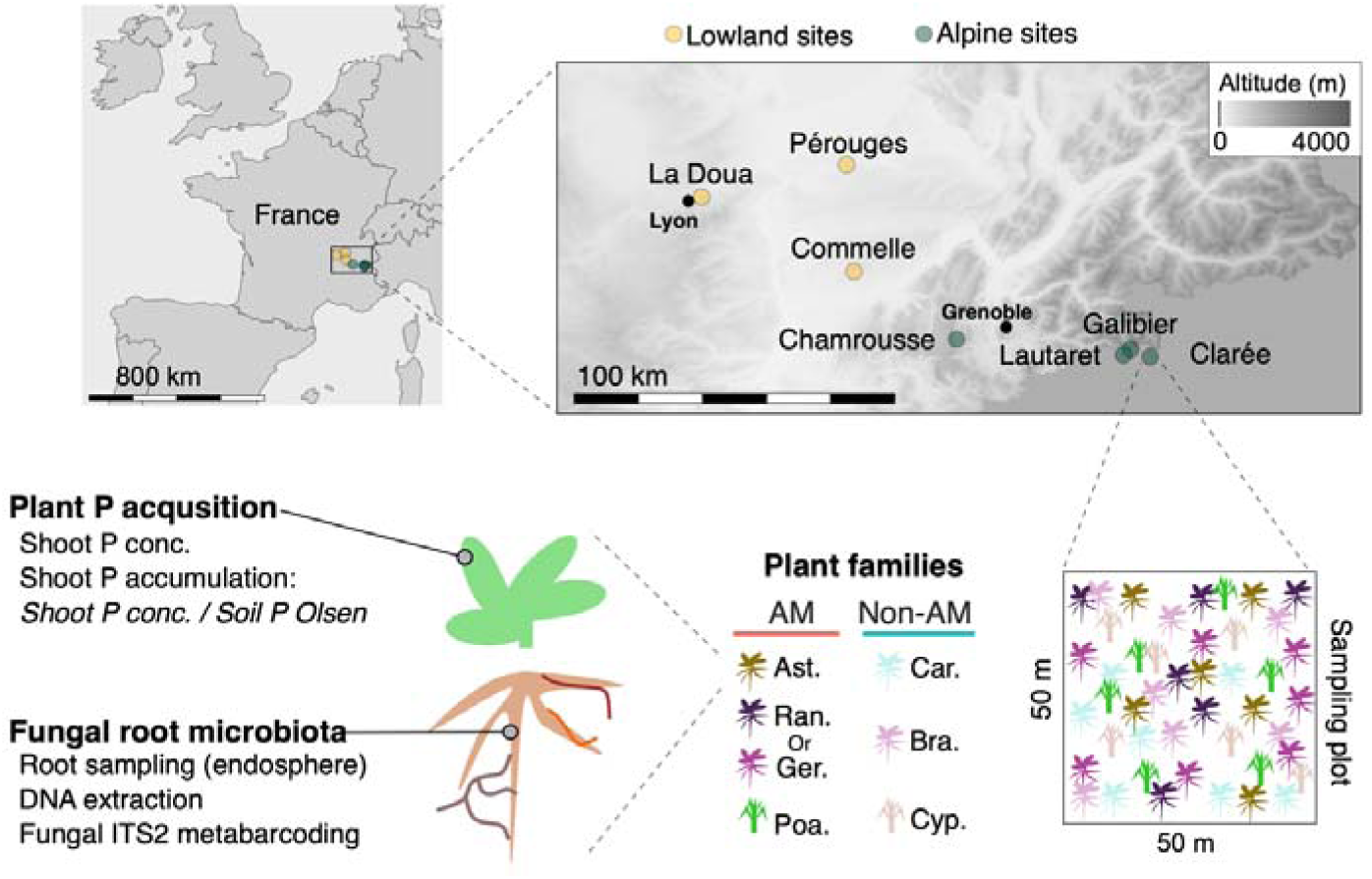
Experimental setup showing the geographical locations of the sampled natural grassland sites (Tab. S2-3), the plant families sampled (Tab. S4), and the analyses conducted on each plant individual. Abbreviations: Ast., Asteraceae; Ger., Geraniaceae; Poa., Poaceae; Car., Caryophyllaceae; Bra., Brassicaceae; Cyp., Cyperaceae; Ran., Ranunculaceae. Six individuals were sampled per family (6) and site (7) for a total of 252 plants.

### In low P soils, non-AM plants tend to have lower shoot P levels than AM plants

Because non-AM plants cannot rely on the AM symbiosis to improve their P uptake, we examined how this affects plant P levels in natural settings, using shoot P as a proxy. We compared neighboring AM and non-AM species across seven grassland sites with contrasting levels of plant-available P (Fig. 2, Tab. S2-4), including low-P alpine grasslands (P Olsen < 17 mg kg^−1^) and high-P lowland grasslands (P Olsen > 25 mg kg^−1^). ANOVA on shoot P levels (model: shoot P ∼ Site/AM-status/Plant Family) showed major effects of plant family (*R^2^* = 0.49; *P* < 0.001) and sampling site (*R^2^* = 0.30; *P* < 0.001), with a marginal but significant effect of the plant mycorrhizal status (*R^2^* = 0.07; *P* < 0.001), indicating lower shoot P levels in non-AM plants (−21%). However, within-site analyses showed that this trend was significant only in the two alpine sites with the lowest P levels (−32% at Galibier and −37% at Lautaret; ANOVA, *P* < 0.05).

**Figure 3.**
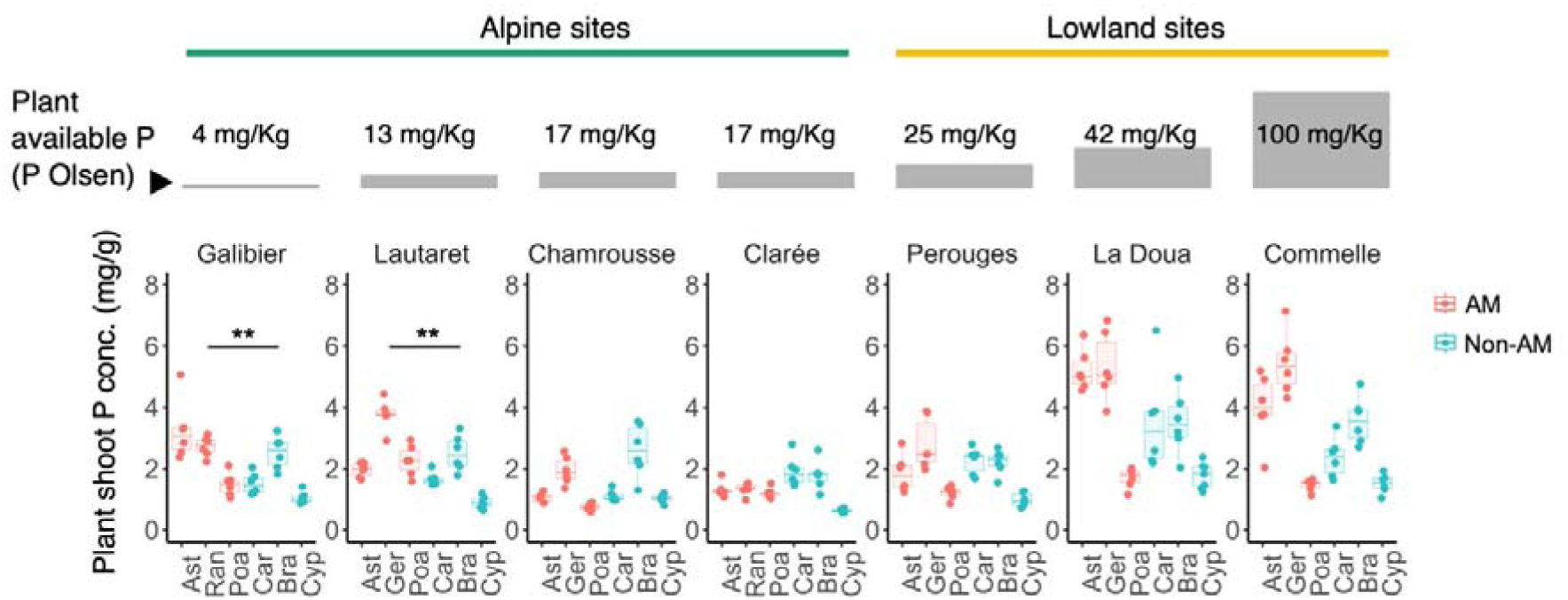
Plant shoot P levels across sampling sites, plant families, and plant mycorrhizal status. (A) Shoot P concentrations (mg/g) measured by ICP-optics for each AM (pink) and non-AM (blue) plant family across sampling sites. Sites are ordered according to levels of plant-available P in the soil (P_Oslen_). Abbreviations: Ast., Asteraceae; Ger., Geraniaceae; Poa., Poaceae; Car., Caryophyllaceae; Bra., Brassicaceae; Cyp., Cyperaceae; Ran., Ranunculaceae. Asterisks indicate significant differences between AM and non-AM plants within sites (ANOVA, *P* < 0.01) (n = 6 plants per species per site).

### Minimal fungal root microbiota differences between AM and non-AM plants

To test the hypothesis that non-AM plants might rely on alternative fungal associations that support their P nutrition, we surveyed the fungal root microbiota of the sampled non-AM species using fungal ITS2 metabarcoding, and compared it to that of neighboring AM plants (Fig. 2). Fungal communities varied widely across biomes, sites, and plant families (Fig. S5A), with no significant differences in alpha diversity between AM and non-AM plants (Shannon’s *H*-index and Chao1 index; ANOVA, *P* > 0.05) (Fig. S5B). Similarly, the structure of root-associated fungal communities was influenced by the sampling site (PerMANOVA on BC dissimilarities, model: Fungal structure ∼ Site/AM-status/Plant Family, *R^2^* = 0.16, *P* < 0.001; Fig. 4A) and the plant family (*R^2^* = 0.21, *P* < 0.001; Fig. 4B). Still, a marginal but significant effect of the plant AM-status was detected (*R^2^*= 0.042, *P* < 0.001; Fig. 4C), even after selectively removing all OTUs from the Glomeromycota phylum before analysis (*R^2^* = 0.041, *P* < 0.001; Fig. S6).

**Figure 4.**
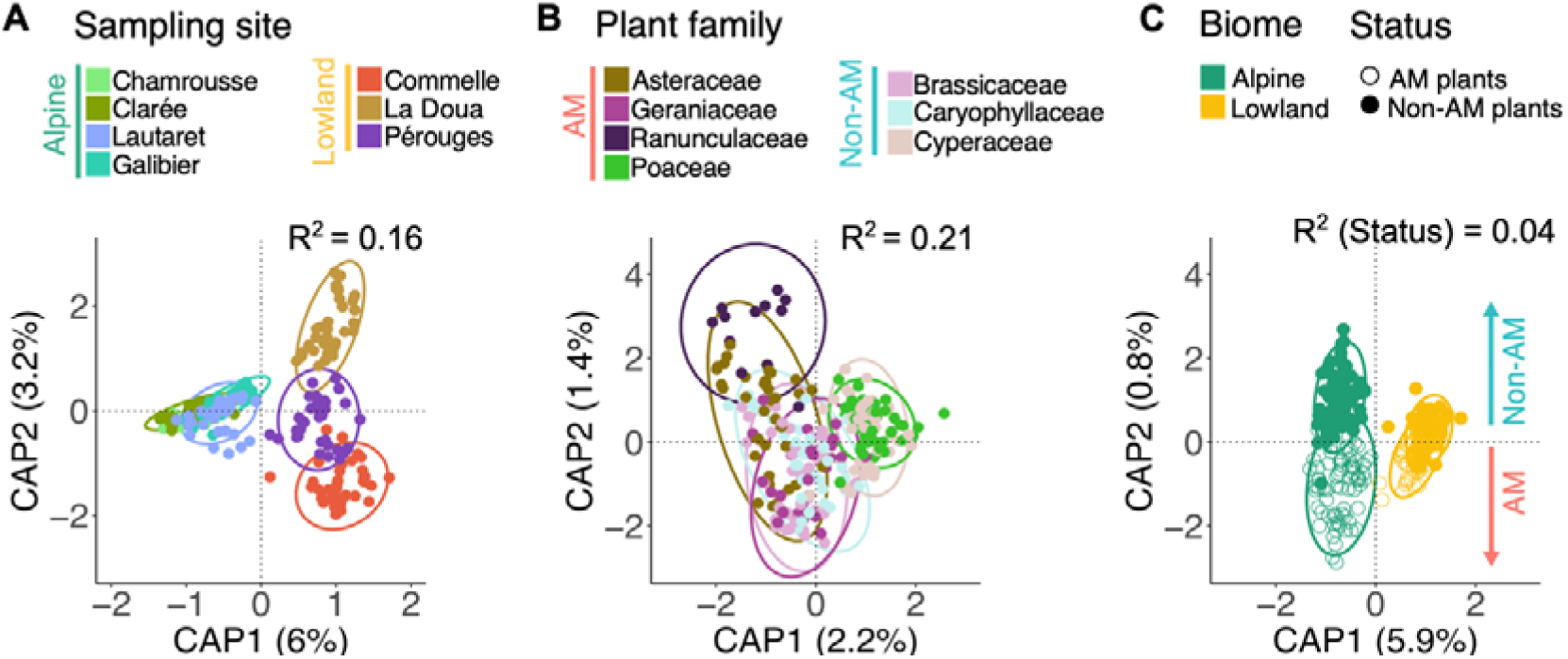
Root fungal microbiota differences between AM and non-AM plants. Fungal community differences across (A) sampling sites, (B) plant families, and (C) mycorrhizal groups within biomes are depicted using distance-based redundancy analysis (db-RDA) on Bray-Curtis (BC) dissimilarities. Ellipses show 95% dispersion surface of data. The variance explained by each factor used to constrain the analysis is indicated at the top (*R^2^*) (PerMANOVA, *P* < 0.05). Similar results were observed when excluding all Glomeromycota OTUs (Fig. S6).

With the aim to identify specific fungal signatures in non-AM plants, we used a mixed approach combining Random Forest modelling with db-RDA graph analysis, which detected 104 fungal OTUs differentially enriched between AM and non-AM plant roots, representing 1.3% of all OTUS detected (8000) (Tab. S5). Several of these differences concerned the enrichment of Glomeromycota lineages in AM plant roots (19/104 OTUs), consistent with measured root mycorrhization levels and Glomeromycota detection at most sites (Fig. S7). OTUs enriched in non-AM plant roots (55/104) showed only quantitative differences with no fungal taxon found exclusively in non-AM plants (Fig. S8, Tab. S5).

### Diverse fungal taxa associated with higher P-accumulation in non-AM plants

We further employed a machine-learning approach to specifically identify and compare fungal taxa associated with better plant P-acquisition in AM and non-AM plants. P-accumulation in plant shoots, calculated as [Plant shoot P concentration] / [Plant-available P (Olsen)], was used as a proxy for plant P-uptake, as done in agronomical studies (e.g. [27]) (Fig. 2). The trained Random Forest regression model showed excellent performance in predicting plant P-accumulation from root microbiome data in low-P alpine sites, reaching 79% of explained variance for AM plants and 69% for non-AM plants (*R^2^* scores of predicted *vs.* observed curves). Although lower performance was observed in high-P lowland sites, where plant P-accumulation values were lower (Fig S9), *R^2^* scores remained good, reaching 58% for AM plants and 57% for non-AM plants.

A total of 165 fungal OTUs that were identified as best predictors of plant P-accumulation based on their importance in the model (mean MSE) were further verified by graph analysis (db-RDA) and Spearman rank correlation (Fig. 5, Tab. S6). Fungal OTUs associated with higher plant P-acquisition were different in alpine (100 OTUs) and lowland sites (60 OTUs), with only 5 OTUs common to both biomes. While 80 of these 165 OTUs were best predictors of P-accumulation in AM plants (48 in alpine sites, Cluster C1; 32 in lowland sites, Cluster C4; Fig. 5), and 55 in non-AM plants (33 in alpine sites, C3; 20 in lowland sites, C6; 2 in both, C2), 30 were common to both plant groups (19 in alpine sites, C2; 8 in lowland sites, C5; 3 in both, C2) (Fig. 6).

**Figure 5.**
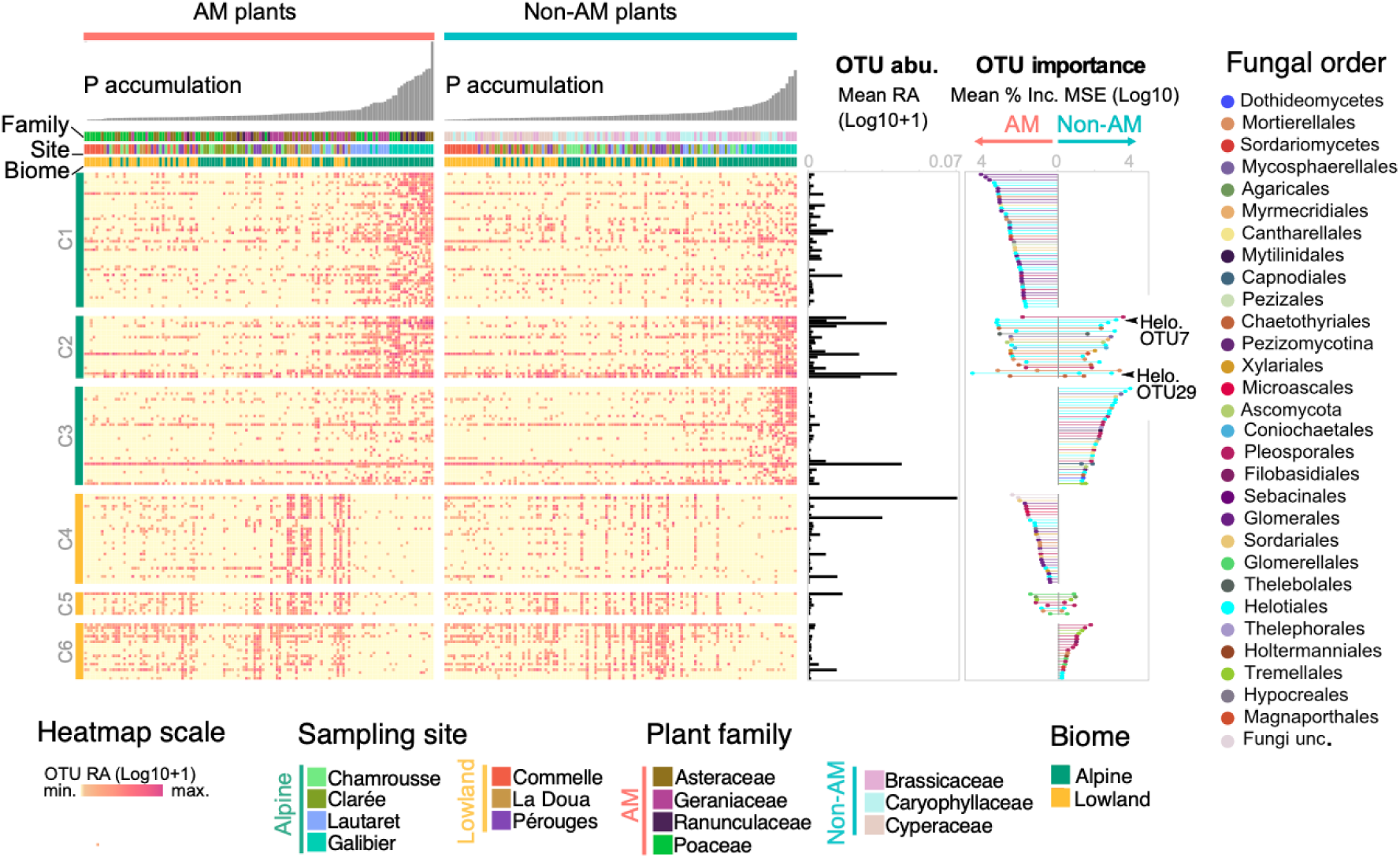
Machine learning-based identification of fungal taxa associated with higher plant P-accumulation. Heatmap showing per-sample relative abundances of OTUs predicted to be associated with higher plant P-accumulation in AM and non-AM plants growing in alpine or lowland biomes (165 OTUs). Samples are ordered by plant P-accumulation values. Taxa identified by Random Forest were verified by db-RDA and Spearman correlation. Plots on the right side show OTU mean relative abundance (mean RA) and importance in the Random Forest model (mean percentage increase in the model’s MSE) in AM or non-AM plants. For OTUs identified in both environments, two values (dots) are shown (e.g. for OTU29). Importance values are colored by OTU taxonomy (order). OTU7 and OTU29 are highlighted. Abbrevations: Unc., Unclassified.

**Figure 6.**
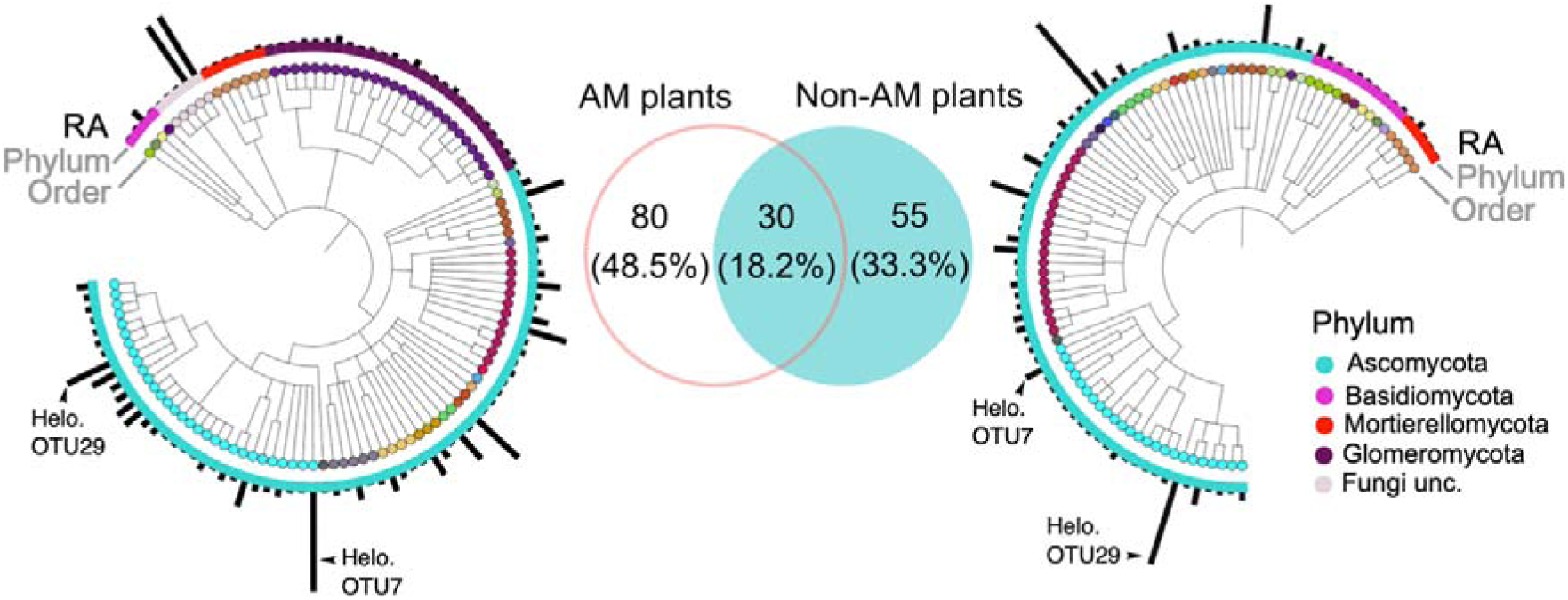
Taxonomical distribution of fungal taxa associated with higher plant P-accumulation in AM *vs.* non-AM plants. Venn diagram and dendrogram representing the number and taxonomy of OTUs associated with higher plant P-accumulation in AM (left; 110 OTUs) and non-AM plants (right; 85 OTUs). In the dendrograms, OTUs are colored by taxonomy (order and phylum), and histograms on the outer circle show OTU mean relative abundance in AM or non-AM roots. Abbrevations: Unc., Unclassified.

In non-AM plants, Helotiales was the most overrepresented order (24/85 OTUs, 28%). Six OTUs belonging to the *Tetracladium* genus were identified. Among these was a highly abundant and prevalent *T. maxilliforme* lineage (OTU29; 0.9% average relative abundance; Fig. S10), which showed the strongest association with higher shoot P-accumulation in non-AM plants (Fig. 7). Similar results were observed with another Helotiales lineage closely related to the mycorrhizal-like fungus F229 (OTU7; 0.85% average relative abundance; Fig. S10), albeit only in alpine sites (Fig. 7). Other identified Helotiales included taxa with described plant-beneficial activities [28,29] such as *Lachnum* (1 OTU), *Hyalopeziza* (3), and *Laetinaevia* (4) taxa. Pleosporales (20/85 OTUs, 23%) were the second most represented order. Most OTUs belonged to lineages with complex ecologies, described both as plant pathogenic and plant beneficial endophytes in the *Didymellaceae* family (5 OTUs) and *Alternaria* genus (2) [30]. Other noteworthy taxa included a *Colletotrichum* (Glomerellales), closely related to the mycorrhizal-like fungus *C. tofieldiae* [18], eight Mortierellales OTUs (including seven *Mortierella* OTUs), and a highly abundant *Cladosporium* taxon (Fig. 5-6, Tab. S6).

**Figure 7.**
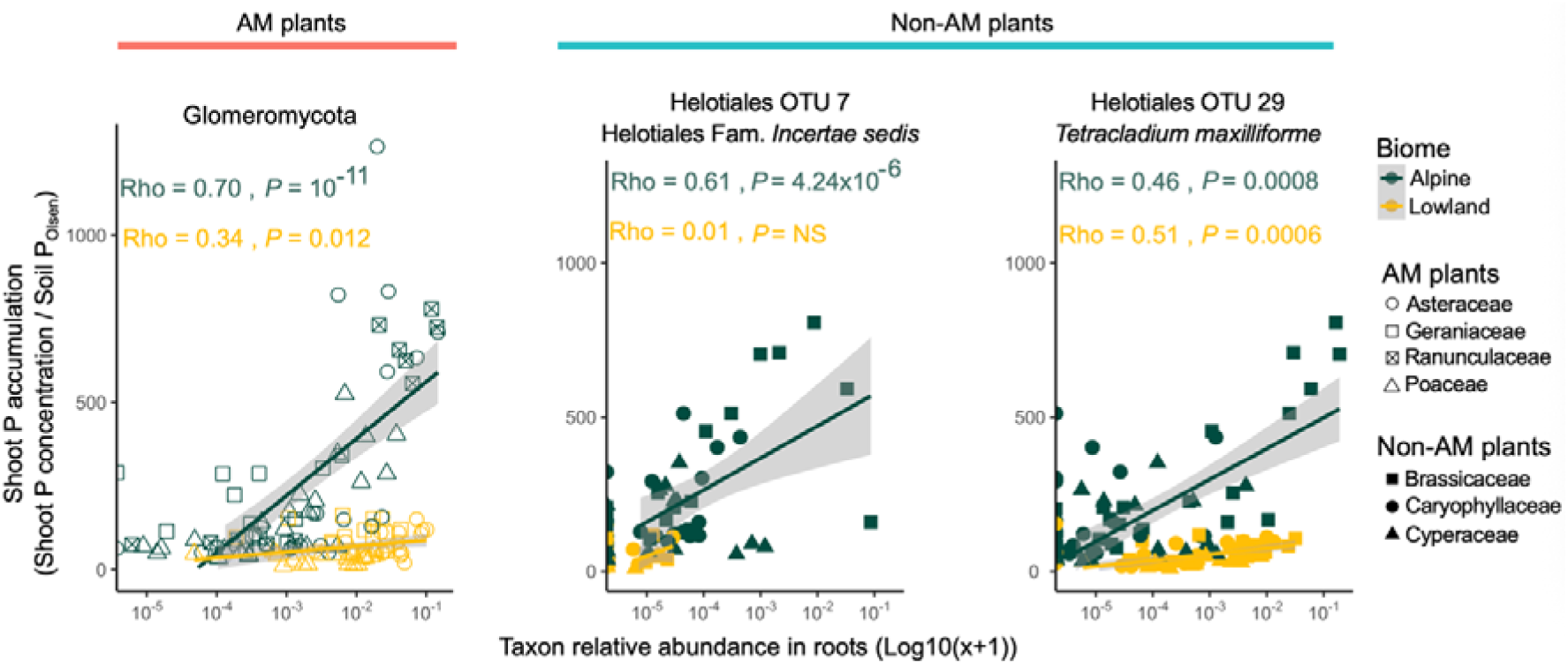
Correlation between shoot P-accumulation in plants and relative abundance of key fungal taxa in roots within each biome. Each point represents an individual plant. Correlations were calculated across AM or non-AM plants within each biome. Shoot P-accumulation was calculated as shown in Fig. 2. Glomeromycota relative abundance in roots was estimated by summing the relative abundances of individual Glomeromycota OTUs (ITS2 metabarcoding). Spearman correlation coefficients (rho) and associated *P*-values are indicated for each condition. For other correlations tested see Fig. S11.

In AM plants, as much as a quarter of the identified OTUs belonged to the AM fungal order of Glomerales (26/110 OTUs) (Fig. 6, Tab. S6). While the relative abundance of Glomeromycota in AM roots was low (4% average; Fig. S7C), it was highly correlated with P-accumulation in shoots. This was true for lowland sites (rho = 0.34, *P* = 0.012) and alpine sites (rho = 0.70, *P* = 10^−11^), with a stronger correlation in the latter (Fig. 7). However, Glomeromycota levels showed weak correlation with measured mycorrhization rates (rho = 0.36, *P* = 0.03). Still, the identification of Glomeromycota as an expected key group demonstrated the efficacy of our proposed method in identifying fungi key for plant P-acquisition. Surprisingly, 30 of the lineages identified as important in AM plants overlapped with those identified in non-AM plants. This concerned 9 Helotiales, with this order being the most overrepresented both in AM and non-AM plants (31 of 110 and 24 of 85 OTUs, respectively) (Fig. 6). Lineages *T. maxilliforme* OTU29 and Helotiales OTU7 were also associated with higher shoot P-accumulation in AM plants (Fig. S11), sometimes exhibiting higher abundances in AM than in non-AM plants, in the case OTU7 in alpine sites (Wilcoxon test, *P* = 0.01, FDR corrected; Tab. S5).

### Helotiales enhance growth and P-acquisition of non-AM hosts upon inoculation

With the hypothesis that associations with Helotiales could be of prime importance for the P nutrition of non-AM plants growing in P-limited alpine soils, two isolates belonging to key lineages Helotiales sp. OTU7 (strain 1_39) and *T. maxilliforme* OTU29 (strain 1_40) were selected for further *in planta* experiments. These fungi were previously isolated from the roots of non-AM *Arabis alpina* growing at Galibier (low-P alpine site) [31] (Fig. 2). Plants were grown in low-P microcosms using seeds collected from wild populations of *A. alpina* and *Carex sempervirens* Vill. (at the Galibier site) or commercial seeds for *Minuartia verna* L. (Fig. 8A). At 21 days after inoculation of *A. alpina*, both fungi had improved plant growth (+ 57-75% shoot biomass, Wilcoxon tests, *P* < 0.024), shoot P concentration (+ 42-88%, *P* < 2.9 × 10^−5^), and total shoot P content (+ 136-154%, *P* < 4.7 × 10^−7^). Similar effects were observed on growth (+ 19-38% shoot biomass, *P* < 0.03), shoot P concentration (+ 68-85%, *P* < 3.9 × 10^−11^), and total shoot P content (+ 114%, *P* < 3.7 × 10^−12^) of *M. verna* (Fig. 8B). Alongside these beneficial effects, the plants remained completely asymptomatic (Fig. 8C) despite extensive fungal root colonization (Fig. 8D, Fig. S11). Although neither Helotiales 1_39 nor *T. maxilliforme* 1_40 significantly improved *C. sempervirens* growth, inoculation of isolate 1_40 resulted in higher shoot P concentration (+ 30%, *P* < 0.032) (Fig. 8BC). While both isolates could colonize plant root surface and cortex, with evidence of intra-cellular colonization in all three plant species (Fig. 8D, Fig. S12A), higher colonization rates were observed with Helotiales 1_39 (Fig. S12B). These results indicate that fungi from these phylogenetically-distant Helotiales lineages have the potential to enhance P-acquisition in diverse non-AM plants, albeit with stronger effects on Brassicaceae and Caryophyllaceae than on Cyperaceae.

**Figure 8.**
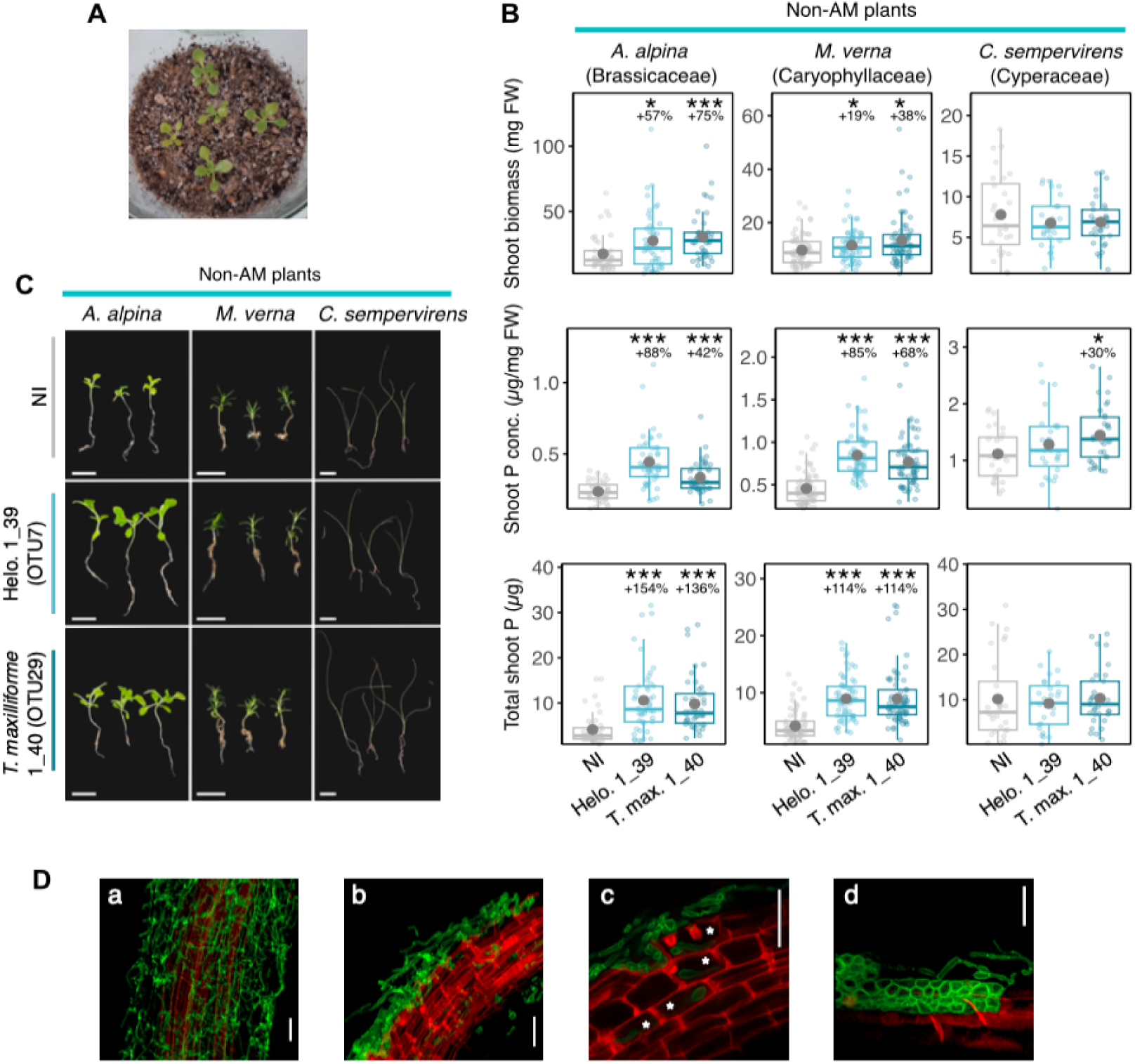
Root endophytic Helotiales 1_39 (OTU7) and *T. maxilliforme* 1_40 (OTU29) enhance growth and P-acquisition of diverse non-AM hosts in low-P soil microcosms. (A) *A. alpina* grown 21 days in sterile sand-perlite-peat microcosms amended with 0.5X low-P Hoagland solution (100 µM P) either non-inoculated (NI) or inoculated with isolate 1_39 or 1_40 (10^4^ propagules per plant). (B-C) Effects of isolates 1_39 and 1_40 on shoot fresh weight (FW), shoot P concentration (conc.), and total shoot P content. The experiment was done twice with 4-8 microcosms per treatment and 4-5 plants per microcosm. Compiled results from both experiments are shown. All measurements were done on individual plants (n = 29-62 plants per treatment). Shoot P was measured by the molybdenum blue method. Asterisks indicate significant differences with the non-inoculated (NI) treatment (Wilcoxon test; * *P* <0.05, *** *P* < 0.001). Scale bars represent 1.5 cm. (D) Examples of fungal root colonization visualized by confocal microscopy after staining the fungal cell wall (WGA-Alexa, green) and the plant cell wall (propidium iodide, red). Isolate 1_40 in *A. alpina* (a) or *C. sempervirens* (d) and isolate 1_39 in *A. alpina* (b-c). Asterisks indicate intracellular colonization by the fungus (scale bars indicate 20 µm) (see Fig. S12).

### Direct transfer of P and N from Helotiales to non-AM hosts

While little is known about the ecology of root endophytic *T. maxilliforme* fungi, previous work on isolate F229, phylogenetically close to Helotiales 1_39, demonstrated its capacity to transfer P to *A. alpina* [19]. Using compartmentalized soil microcosms classically employed in AM research [6], we tested the capacity of Helotiales 1_39 and *T. maxilliforme* 1_40 to improve P (and N) acquisition in Brassicaceae and Caryophyllaceae non-AM hosts via direct nutrient transfer. The AM fungus *Rhizophagus irregularis* and its host *Allium ampeloprasum* (leek) were included as mycorrhizal references (Fig. 9A). Helotiales inoculation significantly increased total shoot P content in both *A. alpina* and *M. verna*, and shoot N content in *M. verna* but not *A. alpina* (pairwise Wilcoxon test, *P* < 0.05, FDR corrected) (Fig. S13). Two weeks after adding the ^33^P and ^15^N sources in the fungal compartment, both isotopes were detected in the shoots of inoculated plants at levels significantly higher than in non-inoculated controls (Fig. 9B), indicating that both fungi were able to transfer P and N to their non-AM hosts. Conversely, inoculation of these Helotiales in the AM model host *A. ampeloprasum* did not improve plant P or N acquisition (Fig. S13). While ^33^P and ^15^N were detected in shoots, the levels were much lower than observed with *R. irregularis* (21 fold difference for ^33^P, 5 fold difference for ^15^N) (Fig. 9B), suggesting limited nutrient transfer by the Helotiales to the AM host (Fig. 9B). In all treatments, fungal colonization was consistently restricted to roots, with no fungal structures observed in stems or leaves (Fig. S14).

**Figure 9.**
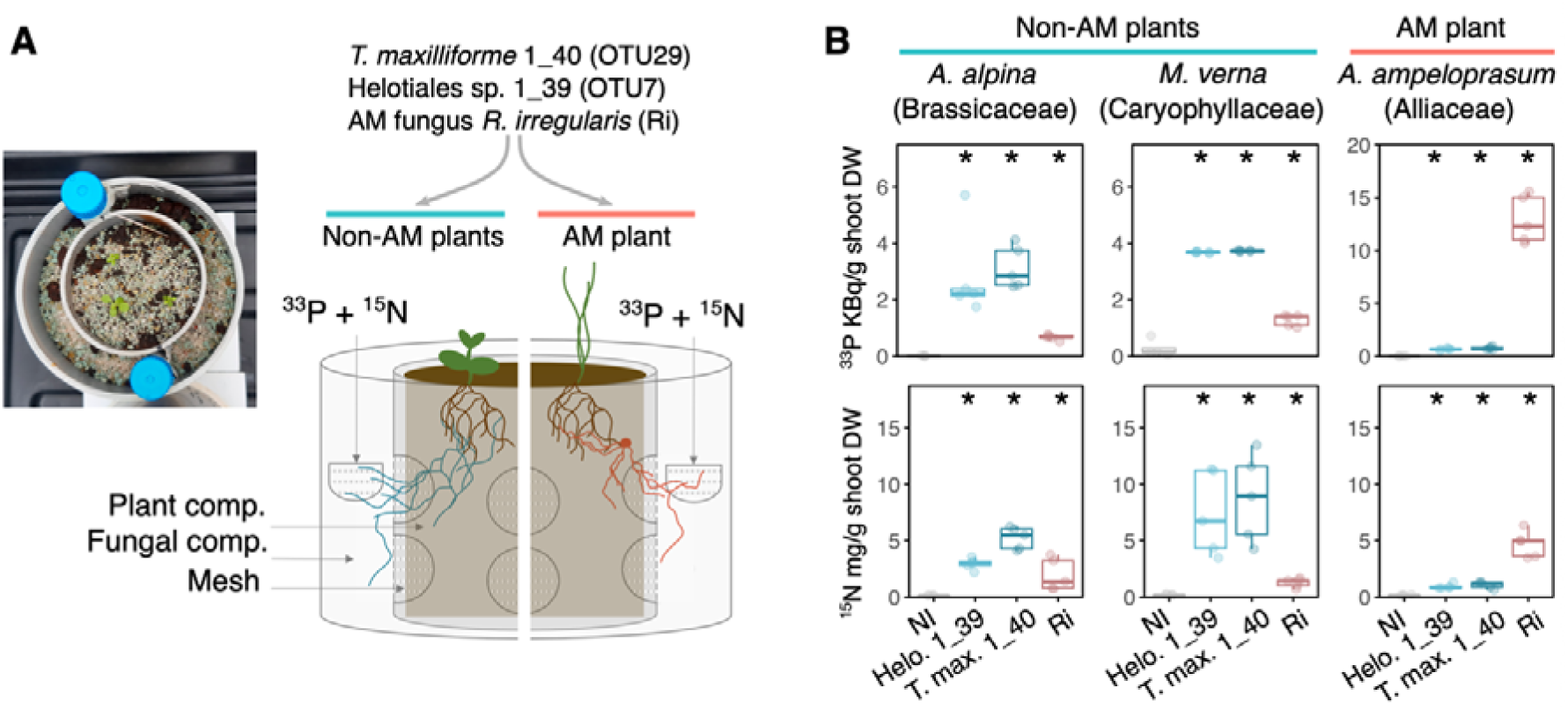
Helotiales 1_39 and *T. maxilliforme* 1_40 transfer P and N to their non-AM host plants. (A) Non-AM *A. alpina* and *M. verna* as well as the model AM plant *A. ampeloprasum* were inoculated with Helotiales isolate 1_39 or 1_40, or the AM fungus *R. irregularis* (Ri) in low-P microcosms used to detect P and N transfers from fungi to plants. A nursing plant was used when inoculating the AM fungus on non-AM plants. ^33^P-labelled H_3_PO_4_ (50 KBq) and ^15^N-labelled (NH_4_)_2_SO_4_ (10 mg) were added to the fungal compartment (comp.) in a nylon mesh pocket (21 µm grid). The fungal compartment was separated from the plant compartment by a plastic cylinder pierced with holes covered by a second nylon mesh (20 µm grid), thus ensuring that plants could only access ^33^P and ^15^N through hyphal transfer. Fungi were inoculated at the start of the experiment (10^5^ mycelial propagules per microcosm for 1_39 and 1_40, or 200 spores for *R. irregularis*) and plants were added 3 weeks later. After 3 weeks of fungal colonization of the roots, ^33^P and ^15^N were added to the fungal compartment and measured in plant shoots 2 weeks later. (B) ^33^P and ^15^N levels in plant shoots in the different treatments, with 4-5 microcosms per treatment (the 3-4 plants per microcosm were pooled at sampling). Water was added to the fungal compartment in the non-inoculated (NI) treatments (n = 4-5).

Surprisingly, inoculation of the AM fungus *R. irregularis* also led to the detection of ^33^P and ^15^N in shoots of non-AM plants, though at levels much lower than observed with Helotiales (3-5 fold difference for ^33^P, 1.5-7 fold difference for ^15^N), or with *R. irregularis* in the AM host *A. ampeloprasum* (16 fold difference for ^33^P, 5 fold difference for ^15^N) (Fig. 9B). However, these results should be interpreted with care as the treatments are not directly comparable. Indeed, because AM fungi cannot grow in the absence of a compatible host [32], establishment of the fungus in the non-AM plant microcosms had to be aided using *A. ampeloprasum* as a nursing plant [33] (Fig. 9A), introducing a bias. Taken together, our results show that these Helotiales have the capacity to transfer significant amounts of N and P to non-AM plants.

## Discussion

Numerous plant lineages have independently lost the capacity to establish the ancestral AM symbiosis [11,12]. Here we tested the hypothesis that prominent non-AM lineages might rely on overlooked nutritional associations with new types of nutrient-delivering fungi. In Brassicaceae, symbiosis abandonment has been dated to 88-97 MYA [34], but our screening of genomes of suspected non-AM lineages identified two additional major AM loss events. One in the last common ancestors of the Caryophyllales order, dated around 111-122 MYA [35], and one in the last common ancestor of the Juncaceae/Cyperaceae families, estimated around 90 MYA [36] (crown ages) (Fig. 1). These results, along with those from nutrient transfer experiments (Fig. 9), confirm that Caryophyllaceae, Cyperaceae, and Brassicaceae do not rely on AM symbiosis to meet their mineral nutrition requirements, prompting us to assess the ecological consequences of AM abandonment in these lineages in terms of plant P-acquisition and fungal recruitment to their roots.

First, our survey of plant shoot P levels in seven natural grassland communities showed non-AM plants reach P levels comparable to those of their AM neighbors in most soils. However, in soils with very low P availability, non-AM plants showed reduced shoot P concentrations (Fig. 3). This warrants careful interpretation as plant species can differ in phosphorus use efficiency, meaning that lower shoot P concentrations do not necessarily indicate P deficiency [37]. Non-mycorrhizal lineages may have evolved more efficient P utilization strategies or have different P allocation patterns, prioritizing, for example, P investment in roots. Further investigation of P use efficiency and P allocation in a large set of non-AM species is needed to test this hypothesis.

Second, our search for unique fungal microbiota signatures in these plants showed that root microbiota of AM and non-AM plants were only marginally different (Fig. S8). This is in line with previous observations made on cultivated and model non-AM plants [38] and does not support the hypothesis of a fungal symbiont switching (i.e. replacing ancient symbionts with novel ones) in these non-AM lineages.

This result prompted us to develop a more analytical approach, leveraging the variability in microbiome composition and shoot P-concentrations observed across sampled plant individuals, species, and families to identify fungal taxa associated with enhanced plant P-acquisition using machine learning. The Random Forest regression model demonstrated strong predictive performance in low-P alpine soils (*R²* = 0.79 for AM plants, 0.69 for non-AM plants), suggesting that fungal community composition is a substantial predictor of plant P-accumulation under P-limited conditions. This performance is particularly notable given the complexity of field-collected microbiome data and the multiple environmental and genetic factors influencing plant nutrition, and it compares favorably with similar studies in agricultural settings [22,39]. Furthermore, this approach successfully identified Glomeromycota as key taxa associated with higher P-accumulation exclusively in AM plants (Fig. 5-7), consistent with their established role in plant P nutrition. However, Glomeromycota abundance in roots did not strongly correlate with microscopy-based root colonization rates. Such discrepancies between molecular and microscopic assessments have been reported previously (e.g. [40]) and can depend on the host plant species [41]. It is likely that here these discrepancies result from methodological limitations as mycorrhization levels were estimated from root system subsamples, and can hence fail to capture spatial heterogeneity [42]. Alternatively, they could result from fungal primer biases against certain Glomeromycota species [43] or from biased estimation of fungal biomass by ITS-based methods [44]. Despite these complexities, the consistent identification of Glomeromycota as predictors of P-accumulation in AM plants supports the utility of this framework for linking root fungal communities to host nutritional status.

When applied to non-AM plants, the method identified 85 fungal taxa best predicting plant P-accumulation, with notably, 24 belonged to Helotiales lineages, including important predictor taxa Helotiales sp. OTU7 and *Tetracladium maxilliforme* OTU29 (Fig. 5-7). Helotiales is the largest Ascomycota order, with over 5000 species, 70% of which have an unknown or cryptic ecology [45]. These fungi are particularly enriched in alpine and arctic environments with low nutrient availability, where they often dominate root-associated fungal communities [19,45]. While isolate F229, phylogenetically close to Helotiales sp. OTU7, has been shown to transfer P to its non-AM host *A. alpina* [19], the ecological relevance of similar trophic associations for the nutrition of other non-AM plants remained uncertain. Here we show that two phylogenetically-distant Helotiales lineages – OTU7 (isolate 1_39) and *Tetracladium maxilliforme* OTU29 (isolate 1_40) – can dramatically improve nutrient acquisition of diverse non-AM plants, doubling shoot P content in inoculated plants (Fig. 8) and transferring both P and N to their hosts (Fig. 9). These results broaden our understanding of Helotiales’ role in the nutrition of non-AM plants showing that the capacity for nutrient transfer – specifically N and P – may be widely distributed throughout the fungal order and that these nutritional effects are conserved across phylogenetically distant non-AM host families.

Overall, our unexpected findings that key Helotiales lineages also associate with AM plants and correlate with their P-accumulation raise questions about the specificity of these nutritional partnerships (Fig. 6, Fig. S11). While the reduced nutrient transfer observed when these fungi interact with a non-native AM host suggests some degree of host specificity (Fig. 9), the widespread distribution of these fungi across both AM and non-AM plant roots indicates they are not restricted to non-mycorrhizal lineages. However, the ecological significance of these associations may differ substantially between plant groups. For non-AM plants that lack access to the highly efficient AM symbiosis even modest nutrient contributions from multiple Helotiales lineages may collectively represent a significant fraction of their mineral nutrition, particularly in P-limited environments. In contrast, for AM plants benefiting from Glomeromycota-mediated nutrient acquisition, the same Helotiales associations may instead provide supplementary functions. Further research quantifying the relative nutritional contributions of Helotiales and AM fungi across diverse AM and non-AM host species will be needed to test this hypothesis and determine whether the loss of AM symbiosis has indeed increased the nutritional dependence of non-AM plants on alternative fungal partnerships. Our results are consistent with accumulating evidence that fungi not traditionally recognized as mycorrhizal can transfer nutrients to plants, suggesting this trait is more widespread than previously thought [20,46–48].

### Conclusions

Given the central importance of AM symbiosis for plant nutrition, how non-mycorrhizal plants adapted to the loss of this ancestral symbiosis has remained an open question. This study establishes the biological and ecological importance of associations with diverse mycorrhizal-like Helotiales lineages as alternative nutritional strategies in non-AM plants. To our knowledge, this work represents the first use of predictive modelling to link root-associated fungal communities with host nutritional status in natural plant populations, providing experimentally validated predictions of key microbiota-based mechanisms supporting the nutrition of non-AM plants.

## Methods

### Presence of AM symbiotic genes in AM and non-AM plants

Available genomes from families of interest including known (Brassicaceae) and suspected (Cyperaceae, Caryophyllaceae) non-AM families, as well as closely-related known AM mycorrhizal plant families, were downloaded from the NCBI database (67 genomes; November 2023). The GeMoMa program, developed to predict gene models in target species based on gene models from related reference species using amino acid sequence conservation and intron position conservation [25], was used to screen these genomes for the five major symbiotic genes SYMRK (DMI2; leucine-rich repeat receptor kinase), CCAMK (DMI3; calcium and calmodulin-dependent protein kinase), CYCLOPS (IPD3; transcription factor), STR1, and STR2 (Half-size ABCG transporters). These genes are known to be highly conserved among AM symbiotic plants and lost in non-AM lineages [12]. Each genome was screened twice using gene models from reference model AM species *Oryza sativa* (Poaceae) (IRGSP-1.0 genome, Rice Annotation Project version 2023-09-07) and *Medicago truncaluta* (Fabaceae) (MtrunA17r5.0-ANR, NCBI annotation release 102), taking into account all transcript variants (Supplementary Table 1). Predicted gene models with a “Met” start codon and a “score/aa” value higher than 0.75 were kept, and the predicted proteins were used for phylogenetic analyses to identify putative orthologs. For each gene, all predicted proteins were merged with selected reference sequences before aligning using Muscle (version 5.1) [49]. The obtained alignment was fed into IQ-TREE (version 2.2.5) [50,51] for best substitution model determination (best-fit model), Maximum-Likelihood tree inference, and bootstrapping (SH-LRT method). The obtained gene phylogenetic trees were visually analyzed to identify putative orthologs clustering with reference sequences of interest (Fig. S1-4). Each analyzed plant genome was then scored for the presence of at least one putative ortholog per gene (Fig. 1). The species tree for the analyzed genomes was produced in R using PhyloMaker [26] and the plant megatree.

### Plant sampling in natural conditions

To characterize the fungal microbiota of non-AM plants, three non-AM and three AM mycorrhizal species were sampled from seven 250-m^2^ natural grassland sites, which have different soil P availability, in the Auvergne Rhône-Alpes region, France (Fig. 1, Tab. S2-4) in June-July 2020. Four of these sites were alpine grasslands (Chamrousse, Clarée, Galibier, and Lautaret) and three were lowland grasslands (Commelle, La Doua, and Pérouges). Six individuals were sampled in each plant population. For each individual, shoots were cut off and used for shoot P measurements. Roots were thoroughly washed with ultrapure water and stored at −20°C prior to DNA extraction and fungal metabarcoding (see below). For each plant population, a composite sample of fine roots was collected and stored in 70% ethanol to assess AM mycorrhization rates. A sample of bulk soil was taken at each site for soil chemistry analysis and fungal metabarcoding.

### Soil analysis and shoot P measurements

Bulk soil samples were analyzed to determine soil characteristics including plant-available P levels (Olsen method) (Tab. S2). All analyses were conducted by the Centre Scientifique Agricole Régional (CESAR laboratory, Ceyzeriat, France), except total P levels in soil, which were done using the molybdenum blue method after mineralization [52]. Shoot P levels in wild plants were quantified by ICP-OPTICS (CESAR laboratory).

In controlled plant inoculation experiments, shoot P levels were measured using the molybdenum blue method after mineralization [52]. Briefly, individual plant shoots were dried (65°C, 18h) before mineralization in 500 µl of nitric acid 65% and 125 µl of hydrogen peroxide (100°C, 2h). Samples were then diluted five times in ultrapure water. Total P was measured using the molybdenum blue colorimetric method [52] adapted for 96-well microplates (Greiner, Les Ulis, France) and using 100 µl of the sample, 100 µl of sodium hydroxide (2M), 50 µl of molybdenum blue solution (containing 7 g.l^−1^ of sodium molybdate, 0.3 g.l^−1^ of potassium antimonyl tartrate and 18% of nitric acid 65%), and 50 µl of ascorbic acid (10 g.l^−1^). The microplate was incubated for 45 min at 40°C and the optic density was read at 880 nm using an infinite 200 PRO (TECAN, Männedorf, Switzerland).

### DNA extraction, ITS2 amplification, and sequencing

Total DNA was extracted from roots and soils (about 500 mg) using the FastDNA SPIN Kit for Soil (MP Biomedicals, Illkirch, France). Before extraction, root samples were mixed with silica beads, dipped in liquid nitrogen, and ground using a FastPrep-24 5G (MP Biomedicals) for 2 × 30 s at 5 m.s^−1^. Extracted DNA was quantified using a spectrophotometer (NanoPhotometer NP80; Implen, Munich, Germany). DNA was diluted 1:2 in Tris-HCl buffer (10 mM, pH 8), and 5 µl were used for PCR amplification. The fungal taxonomic marker ITS2 was amplified using primers fITS7 [43] at 500 nM and ITS4 [53] at 300 nM with universal Illumina adaptors. PCR mix included Q5 High-Fidelity polymerase (New England Biolabs, Ipswich, MA) at 0.02 U.µl^−1^, Q5 buffer at 1X, Q5 High-GC enhancer at 1X, and 200 µM of dNTPs. When PCR amplification was unsuccessful, genomic DNA was further diluted at 1:100, and spermine (0.06 mM) was added to the PCR (4 samples). PCR cycling conditions were 98°C for 2 min followed by 30 amplification cycles (98°C × 10 s, 57°C × 30 s, 72°C × 15 s), followed by a final elongation at 72°C during 2 min. Two PCRs were performed per sample before pooling. PCR products were purified using magnetic beads [54], and ITS2 amplification was verified by gel electrophoresis. The purified amplicons were sequenced by Microsynth AG (Balgach, Switzerland). Library preparation and indexing were done using the MiSeq Reagent V3 kit and sequenced in an Illumina MiSeq instrument (2 × 300 cycles), including 5% PhiX control. Sequence processing was done using mothur v.1.44.3 [55], cutadapt [56], and ITSx [57]. Contigs were assembled using mothur v.1.44.3 [55], and cutadapt [56] was used to delete contigs with more than 2-base differences with primer sequences. The sequences were sorted to only keep those between 100 and 600 bp, with an overlap of at least 5 bases (no mismatch). Sequences with ambiguous bases or with long homopolymers (>10) were discarded. Chimeras were detected using Uchime, ITS2 sequences were extracted using ITSx [57] and were then classified using the RDP classifier and the eukaryote ITS database UNITE (full UNITE+INSD from 29/11/2022 [58,59]) in mothur. Sequences were clustered at 99% similarity using distance greedy clustering (dgc) and only OTUs with more than 50 reads were kept and classified. The most abundant sequence in each OTU was chosen as the representative. Only fungal OTUs were kept. Raw sequence data have been deposited (NCBI SRA Bioproject PRJNA1141433). In our dataset, OTUs belonging to the Helotiales species *Titaea maxilliformis* (current name: *Tetracladium maxilliforme)* were wrongly classified in the Tubeufiales order. Because this is taxonomically incorrect given the taxonomical classification of *T. maxilliforme* (IndexFungorum, https://www.speciesfungorum.org/Names/SynSpecies.asp?RecordID=291360 [60], we have manually reclassified these taxa under the order Helotiales (OTUs 29, 2056, 3592, 4967, 5251, 5784, 6163, 7806, 8160, 8906). OTU relative abundances (RA) were calculated, and values were transformed (log10 (RA+1)) before statistical analyses and visualization.

### Plant inoculation assays in sterile soil microcosms

Inoculation assays were conducted with two Helotiales isolates 1_39 and 1_40, previously obtained from *A. alpina* roots [31], that show a 100% similarity with the representative ITS2 sequences of OTU7 and OTU29, respectively. Inoculation of these fungi on non-AM plants were conducted in artificial soil microcosms composed of a mix of perlite clay (3.1% volume), sand (67.3%), and peat (2.7%). Plant nutrients were supplied using half-strength low P Hoagland’s solution containing (100 µM P; 0.4 ml per g of substrate). Microcosms were sterilized by autoclaving twice with a 48h interval (121°C for 20 min). Commercial seeds of *M. verna* (Jelitto, Shwarmstedt, Germany) and wild seeds from *A. alpin*a (Galibier population) were surface-sterilized (ethanol 70% with triton 0.05X; then ethanol 100%) and stratified in gibberellic acid (150 mg.l^−1^) at 4°C in the dark for 5 days. Wild seeds from *C. sempervirens* (Galibier population) were scarified with 96% sulfuric acid for 5 min, rinsed thoroughly in ultrapure water, then rinsed with gibberellic acid (150 mg.l^−1^). All seeds were left to germinate for 8 to 9 days on Murashige and Skoog (MS) agar medium with 100 µM P [61]. Seedlings with developed roots and cotyledons were then transferred to soil microcosms (4 to 5 per microcosm) and placed in a climatic chamber (CP9000, Aralab, Rio de Mouro, Portugal; 16/8 h 22/15°C day/night cycles, 50% relative humidity, 12000 lux). One week later, a suspension of mycelial propagules (20000 propagules.ml^−1^) obtained from four-week-old PDA plates was added at the foot of each seedling, as described previously [19]. Three weeks later, fresh shoot and root biomass were measured. Roots were conserved in 70% ethanol at 4°C for fungal colonization analyses, and shoots were used for biomass and P measurements, as described above.

### ^33^P and ^15^N transfer experiments

To investigate the transfer of P from fungi to plants, we conducted a greenhouse experiment to analyze nutrient exchange in compartmentalized systems. We tested isolates 1-39 and 1-40, along with the AM fungus *R. irregularis*, on non-AM plants *A. alpina* and *M. verna* and the AM plant *A. ampeloprasum* var. porrum. *R. irregularis*-inoculated *A. ampeloprasum* plants were used as a positive reference for detecting P and N transfers during AM symbiosis. Additionally, a non-inoculated condition for each plant species was included as a control. Compartmentalized systems were filled with a mixture of sterile silica sand (48.1% volume; SRM, Biot, France), zeolite (48.1%; Symbiom, Lanškroun, Czech Republic), and peat (3.85%; Floraguard, Oldenburg, Germany) amended with 0.5x Hoagland’s solution with low P (100 µM). Each microcosm was composed of two nested cylinders. The inner cylinder was approximately 0.75 l (diameter 8 cm × height 15 cm) and contained the plant, and the outer one was approximately 1.7 l (d 12 cm × h 15 cm) and contained the fungus. Three circular openings (3 cm in diameter) were created in the inner cylinder and were covered with a fine nylon mesh (21 µm grid) to confine the roots in the plant compartment while allowing the fungal mycelium to pass the mesh and colonize the plant compartment. The bottom of the cylinder was perforated to allow water to pass through. In the fungal compartment, two 15-ml nylon bags (21-µm grid) were inserted and kept empty until the introduction of 50 kBq of ^33^P carrier-free phosphoric acid (H_3_PO_4_, Hartmann Analytic, Braunschweig, Germany) and 13 g of sand containing 10 mg of ^15^N-labelled (NH_4_)_2_SO_4_ (Fig. 9A). Microcosms were first inoculated with either isolate (10^5^ mycelial propagules per system) and incubated for 3 weeks, allowing for fungal colonization. Ultra-pure water was used for non-inoculated (NI) controls. After this, three to four 10-day-old seedlings of non-AM *A. alpina*, non-AM *M. verna*, or of AM *A. ampeloprasum* were transplanted into the inner cylinders (Fig. 9A). Seedlings were obtained as described above. Because *R. irregularis* cannot grow without a receptive host plant, *A. ampeloprasum* was used to allow the establishment of the AM fungus *R. irregularis* DAOM197198 (200 spores; Premier Tech, Rivière-du-Loup, Canada) when inoculated on non-AM hosts. These nurse plants were removed from the systems a week before adding the isotopic tracers. In all conditions, ^33^P and ^15^N sources were added to the fungal compartment after three weeks of plant-fungus contact. Plant shoots were harvested two weeks later, and all plants from each compartmentalized system were pooled, resulting in 4 to 5 replicates per condition. The shoot material was dried (oven-dried for 24h at 55°C) and used for ^33^P and ^15^N detection and measurement of total of P and N content. For ^33^P, plant material (approx. 500 mg) was incinerated for 8 hours at 550°C and digested in 2 ml of 2% hydrochloric acid. A volume of 1.4 mL of the extract was heated at 95°C under agitation for 10 min, centrifuged (12,000 rpm, 10 min), and 675 µl of the supernatant was mixed with the scintillation cocktail (ratio of 1:5 in 5-ml scintillation vials) for subsequent analysis in in a liquid scintillation counter (Tri-Carb 2100 TR; Packard, Meriden, CT). Total shoot P was measured on the acid-digested samples using the molybdenum blue method [52] with a Shimadzu UV-160 spectrophotometer. Total nitrogen content and ^15^N/^14^N ratio in shoots were measured using an elemental analyzer coupled to a mass spectrometer (Europe Scientific) on 2 mg of ground dried samples. All plants were grown side by side in a greenhouse with 16 h of light (10,000 lux; 220 µmol m^−^² s^−1^) at 24°C and 8 h of night at 18°C, with constant relative humidity (65%). Plants were watered twice a week throughout the experiment. The experiment was performed once. One plant per treatment was used to control for the absence of fungal colonization (Fig. S14).

### Microscopy analysis of fungi in root tissues

AM mycorrhization rates were assessed in wild-growing plants. From each condition approximately 10 fine root pieces from each plant individual were taken and pooled into a composite sample to asses average mycorrhization within the plant population with 20-30 root pieces analyzed for each condition. Roots were cleared in 20% KOH 20% for 30-240 min at 100°C, rinsed three times in sterile ultrapure water, acidified with HCl 0.1 M for 45 min, and stained with a lactoglycerol solution containing 0.005% of Trypan Blue for 15 min at 100°C. Roots were left to discolor in 50% glycerol before being mounted on a glass slide. They were observed at X20-40 magnification using an optical transmission microscope (Axioskop 50; Zeiss, Oberkochen; Germany). For each root piece, the mycorrhization level was scored following the method of Trouvelot *et al.* [62] (Fig. S7).

For fungal colonization analysis via confocal microscopy, roots were treated as described above, except that staining was done with propidium iodide (13 mg.l^−1^; Molecular Probes, Invitrogen, Eugene, OR) and *N*LJacetylglucosamine-targeted wheat germ agglutinin WGA-Alexa fluor^TM^488 conjugate (10 µg.ml^−1^; Invitrogen). Stained roots were observed with a LSM 800 confocal microscope (Zeiss, Jena, Germany) at the Centre Technologique des Microstructures (CTµ, Villeurbanne, France) with excitation and detection parameters set to 561 nm and 570-700 nm for propidium iodide and 488 nm and 500-600 nm for WGA-Alexa Fluor 488 (Fig. 8, Fig. S12). In the ^33^P/^15^N transfer experiment, plant shoots were stained as described above and observed under a stereomicroscope with fluorescence (Leica, Wetzlar, Germany) (Fig. S14).

### Statistical analyses

All statistical analyses were done using R 4.3.2. [63]. Unless stated otherwise, means were compared using ANOVA (*P* < 0.05) followed by Tukey’s HSD tests, with data transformed using the bestNormalize package [64] when normality or homoscedasticity were not respected. The function bestNormalize in the bestNormalize package performs various normalization transformations and then selects one optimal transformation based on minimizing the Pearson P statistic, a test for normality. If conditions were still not met, Kruskal-Wallis test and pairwise-Wilcoxon tests (*P* < 0.05) were conducted. *P*-values were corrected for multiple comparisons when necessary (FDR correction). Plant P-accumulation in shoots was calculated as the ratio of P concentration in shoots divided by plant-available P concentration in soil. These values were used as proxies for plant P-acquisition as traditionally done in agronomical studies [27].

Fungal alpha-diversity was analyzed using the Shannon and Chao1 indices with the phyloseq package [65]. Bray-Curtis (BC) dissimilarities were calculated using the vegan package [66]. To study the effect of different factors on the structure of the fungal community, permutational analysis of variance (PerMANOVA) was done using the adonis2 package (*P* < 0.05, 99999 permutations), considering a fully nested model: BC dissimilarities ∼ Site/Mycorrhizal Status/Family.

Two distinct questions were addressed using machine learning approaches. The first aimed to identify differences between non-AM and AM plant microbiota. To reduce noise and ensure meaningful associations, we selected only OTUs present in at least six samples (2% occurrence). Given high fungal community variability across biomes (Fig. 4), analyses were performed on biome-based subsets: one for alpine sites with low available P (i.e. Galibier, Lautaret, Chamrousse, Clarée) and one for lowland sites with high available P (i.e. Commelle, Pérouges, La Doua). To identify specificities of non-AM plant microbiota, we employed two methods on each dataset. First, we used the capscale function from the vegan package [66] to conduct a distance-based redundancy analysis. This involved constructing a Bray-Curtis (BC) dissimilarity matrix and applying a constrained ordination model, BC ∼ Status. OTU CAP1 scores were extracted, and the most contributing OTUs showing a significant differential enrichment (Wilcoxon test, *P* < 0.05) were retained. Second, a Random Forest (RF) classification was implemented to identify OTUs (features) best predicting the plant mycorrhizal status using the randomForest package [67]. Low abundance OTUs (RA < 0.0001) were filtered out to limit noise in the data. RF parameters mtry and ntree were optimized manually. OTU importance was evaluated by calculating average feature importance (Mean Decrease Accuracy values, MDA) across 10 independent rounds of 5-fold cross validation with 80% train/20% test division of the data. The mean out-of-bag error for classifying AM and non-AM plants are reported for each dataset in (Tab. S7). The top 100 most important features (OTUs) based on the average computed MDA were considered the most important features. Results from both methods were merged, retaining only OTUs common to both analyses (Tab. S5).

The second question aimed to identify OTUs associated with higher plant P-accumulation. As in the first analysis only OTUs present in at least six samples (2% occurrence) were selected and low abundance OTUs were filtered out (RA < 0.0001). Given high fungal community variability across biomes (Fig. 4) the data was divided in four subsets analyzed independently: alpine AM plants, alpine non-AM plants, lowland AM plants, and lowland non-AM plants. Each dataset was analyzed with two methods. First, the capscale function [66] was used to perform a distance-based redundancy analysis, constructing a BC dissimilarity matrix, and applying the model BC ∼ plant P-accumulation. CAP1 scores projecting onto the plant P-accumulation vector were extracted, and the most contributing OTUs showing a significant correlation with plant P-accumulation (Spearman correlation, *P* < 0.05) were retained. Second, RF regression was used. Again, RF parameters were optimized, and OTU importance was evaluated by calculating average feature importance (percentage increase of Mean Squared Errors, “% Inc. MSE”) across 10 independent rounds of 5-fold cross validation, with 80% train/20% test division of the data. Mean of squared residuals for each iteration of the regression model are reported (Tab. S8). Model accuracy was measures by calculating *R^2^* scores (average explained variance) in observed *vs.* predicted curves. Results from both methods were merged, retaining only OTUs common to both analyses (Tab. S6). Results were visualized as heatmaps created using Morpheus software, and taxa dendrograms were computed using the “as.phylo” function from the ape package in R and annotated with iTOL software.

## Declarations

### Ethics approval and consent to participate

’Not applicable’

### Consent for publication

’Not applicable’

### Availability of data and materials

The datasets generated and analysed during the current study are available in the NCBI repository https://www.ncbi.nlm.nih.gov under accession numbers PQ286942-PQ286943 for complete ITS Sanger sequences, PQ323610 - PQ323611 for the ITS2 Sanger sequences, and PRJNA1141433 for the fastq-files generated by Illumina ITS2 sequencing. The code and tables used for the analyses and figures are publicly available via github https://github.com/julianaalma/Root-fungal-microbiota-of-non-mycorrhizal-plants-and-plant-P-nutrition-

### Competing interests

The authors declare that they have no competing interests

### Funding

This work was supported by a PhD scholarship from the French Ministry of Higher Education and Research to P.B., a MOPGA 2023 young researcher fellowship from the French Ministry for Europe and Foreign Affairs for L.G, the French National program EC2CO (Ecosphère Continentale et Côtière), the French National Research Agency (ANR) through the “SymbiLoss” project (grant ANR-22-CE02-0018), and the Défi ISOTOP initiative (CNRS, MITI, France).

### Authors’ contributions

J.A., P.B., and Y. M.-L. initiated the research, conceived and designed the experiments. J.A. and J.D. collected samples from wild-growing plants and P.B. conducted microbiota analyses. P.B, L.G., and J.D. conducted and analyzed *in planta* experiments. P.B. and P.-E.C. conducted radioactive P tracing experiments. P.B., J.A., and Y. M.-L. wrote the manuscript with help from all other co-authors. All authors reviewed the manuscript.

## Acknowledgements

This work was supported by the Lautaret Garden – UAR 3370 (Univ. Grenoble Alpes, CNRS, 38000 Grenoble, France), a member of AnaEE France (ANR-11-INBS- 0001 AnaEE-Services, Investissements d’Avenir frame), and of the eLTER European network project (Zone atelier Alpes, CNRS, UGA 38000 Grenoble France). We specially thank Maxime Rome for collecting seeds from wild plants and Alexandre Noiriel and the Radioactivity Platform (ICBMS, Université Lyon 1) for preliminary tests. We also thank the CTµ platform (Université Lyon 1) for the use of the confocal microscope. Additionally, we are grateful to the PRABI (Université Lyon 1) and the iBio (LEM UMR 5557) for access to computer clusters, and the PGE and ‘Serre’ platforms (FR BioEEnViS) for the use of machines and phytochambers. We also thank Noureddine El-Mjiyad from UMR Agroécologie (Dijon, France) for help with the greenhouse experiment. We specially thank J. Keller and P-M. Delaux for discussions.

## Supplementary figures

**Supplementary Figure 1.**
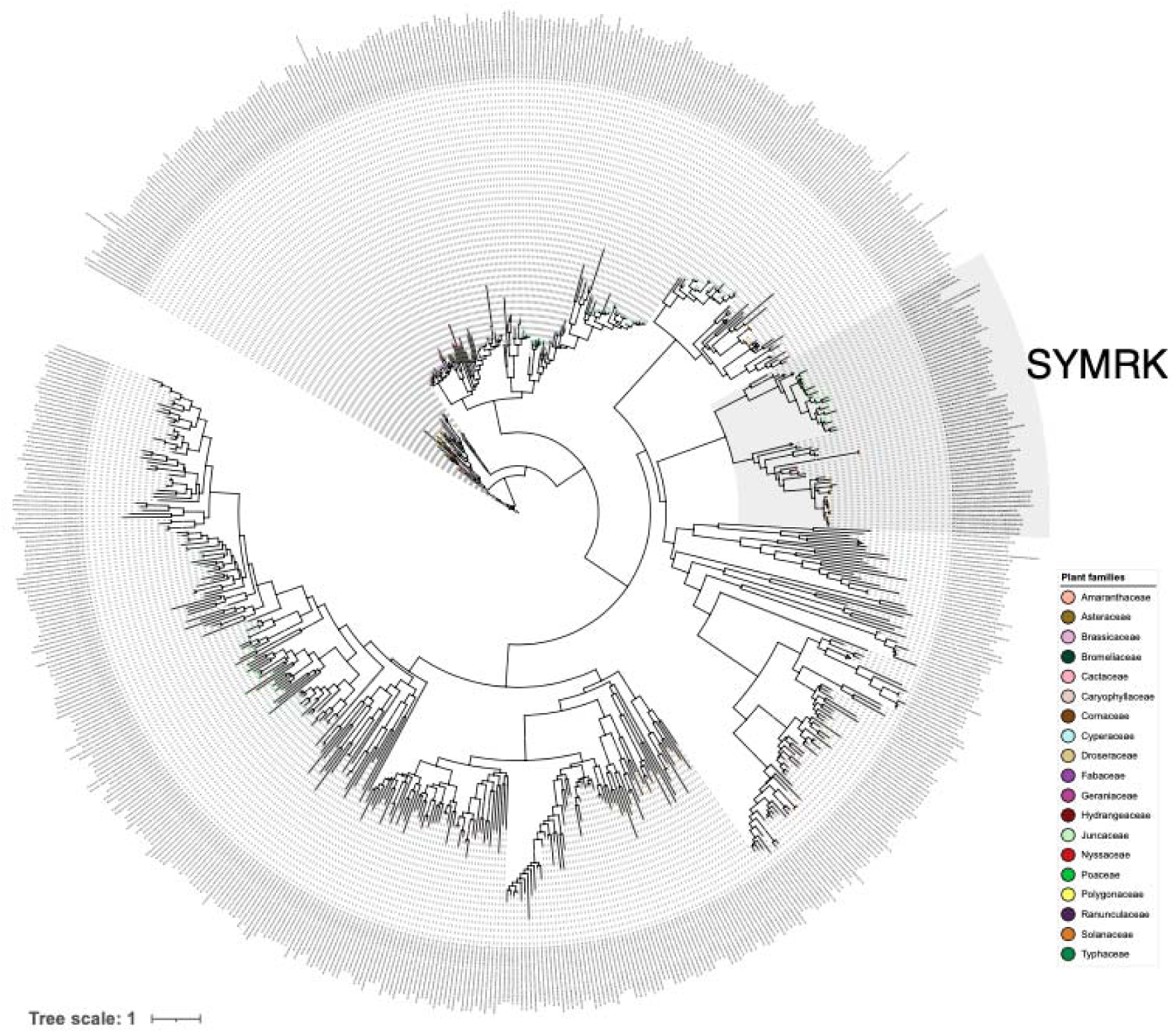
Phylogenetic tree of SYMRK-like proteins predicted from the screened plant genomes. Putative orthologs of *O. sativa* (Os) and *M. truncatula* (Mt) leucine-rich repeat receptor kinase SYMRK, clustering with reference SYMRK sequences (black triangles) are highlighted in grey. Putative orthologs were detected only in AM families (highlighted clades) while proteins clustering with reference sequences representing other types of leucine-rich repeat receptor kinase (white triangles) were detected in AM and non-AM lineages. Maximum Likelihood tree (unrooted), best-fit model: VT+F+R10, blue dots on branches depict 100% bootstrap values (SH-LRT method).

**Supplementary Figure 2.**
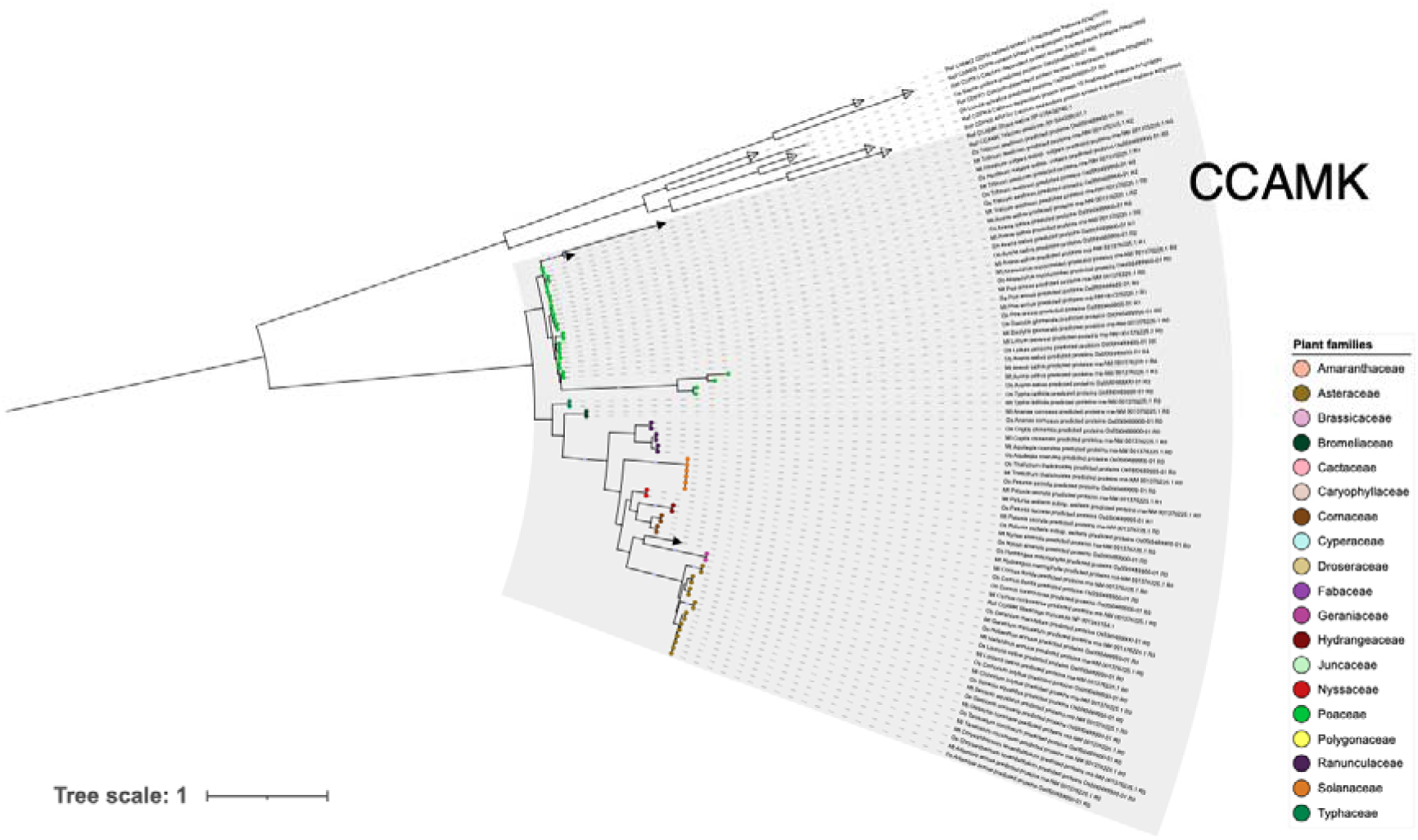
Phylogenetic tree of CCAMK-like proteins predicted from the screened plant genomes. Putative orthologs of *O. sativa* (Os) and *M. truncatula* (Mt) calcium and calmodulin-dependent protein kinase CCAMK, clustering with reference CCAMK sequences (black triangles), are highlighted. Putative orthologs were detected only in AM families (highlighted clade) while proteins clustering with reference sequences representing other types of calcium-dependent protein kinase (white triangles) were detected in two non-AM lineages. Maximum Likelihood tree (unrooted), best-fit model: JTT+R4, blue dots on branches depict 100% bootstrap values (SH-LRT method). Sequences from non-CCAMK calcium dependent protein kinases (white triangles) were used to root the tree.

**Supplementary Figure 3.**
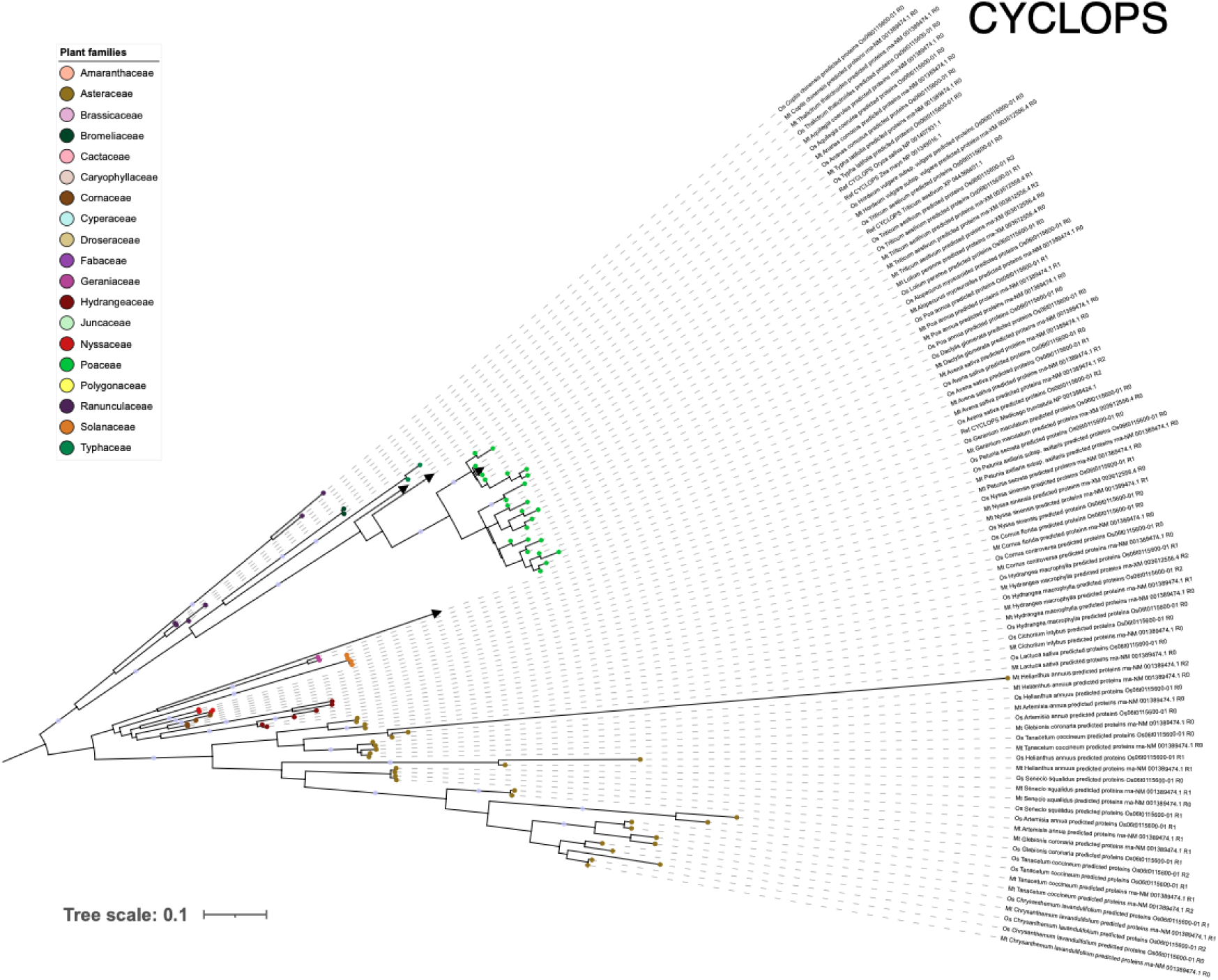
Phylogenetic tree of CYCLOPS-like proteins predicted from the screened plant genomes. Putative orthologs of *O. sativa* (Os) and *M. truncatula* (Mt) transcription factor CYCLOPS, clustering with reference CYCLOPS sequences (black triangles), are shown. These putative orthologs were detected only in AM families. Maximum Likelihood tree (unrooted), best-fit model: Q.mammal+I+G4, blue dots on branches depict 100% bootstrap values (SH-LRT method). Monocot sequences were used to root the tree.

**Supplementary Figure 4.**
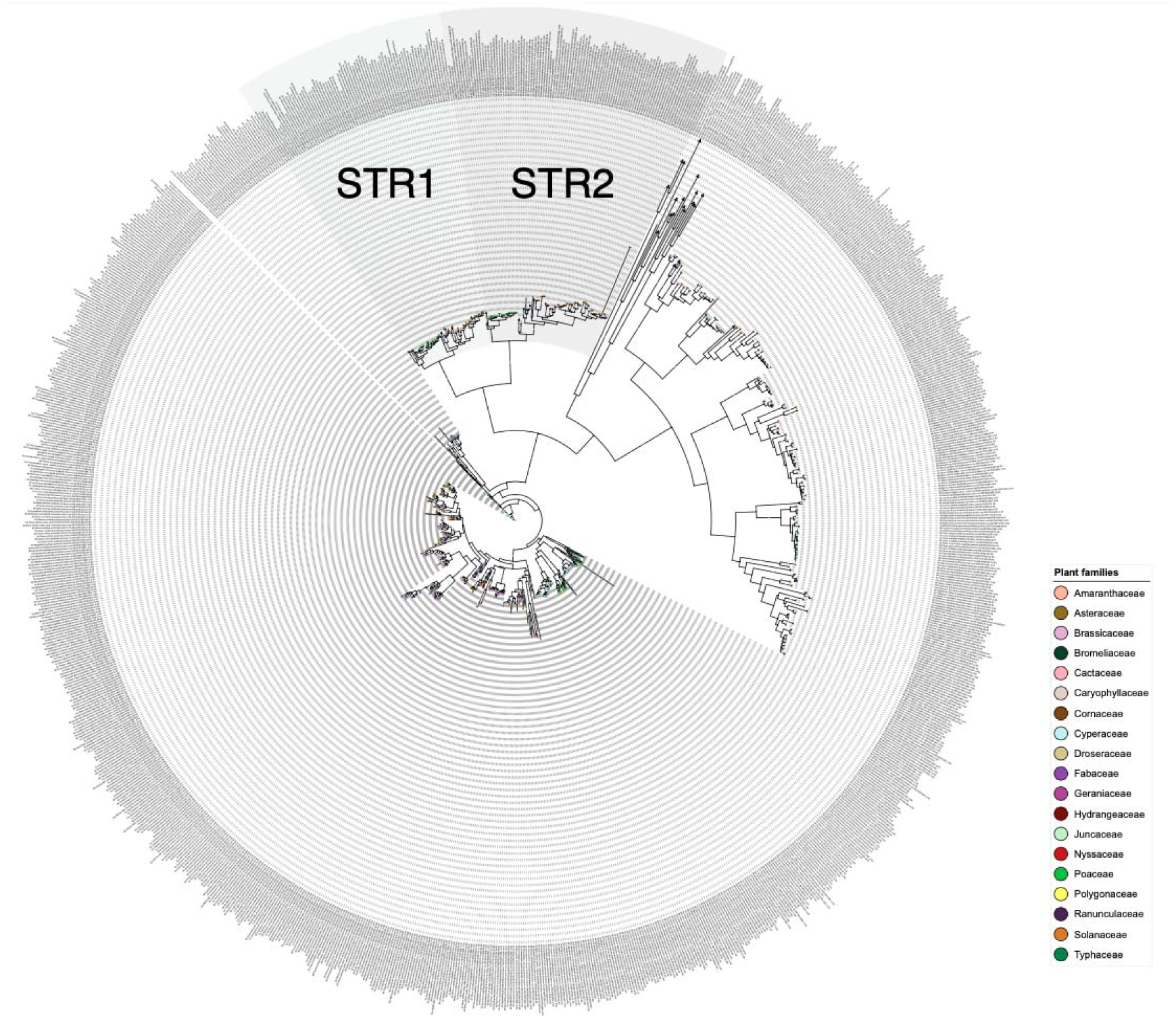
Phylogenetic tree of STR-like proteins predicted from the screened plant genomes. Putative orthologs of *O. sativa* (Os) and *M. truncatula* (Mt) calcium and calmodulin-dependent protein kinase STR1 and STR2, clustering with reference STR1 and STR2 sequences (black triangles), are highlighted. Putative orthologs were detected only in AM families (highlighted clades) while proteins clustering with reference sequences representing other types of ABCG transporters (white triangles) were detected in AM and non-AM lineages. Maximum Likelihood tree (unrooted), best-fit model: JTT+F+R10, blue dots on branches depict 100% bootstrap values (SH-LRT method).

**Supplementary Figure 5.**
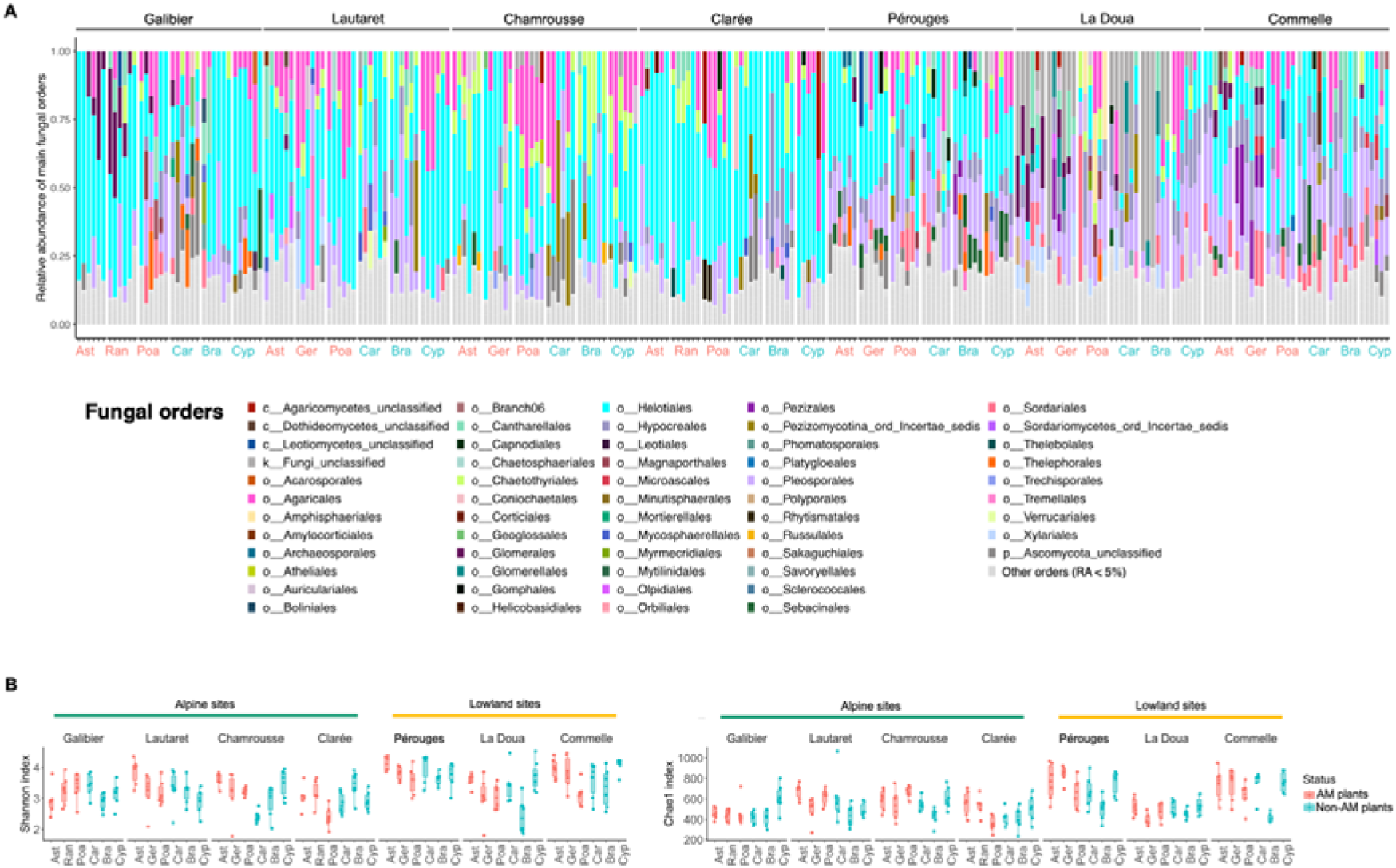
Fungal diversity in roots. (A) Mean relative abundance of the major fungal orders for the different sites and plant family. Abbreviations: Ast, Asteraceae; Ger, Geraniaceae; Poa, Poaceae; Car, Caryophyllaceae; Bra, Brassicaceae; Cyp, Cyperaceae; Ran, Ranunculaceae. AM families are indicated in pink and non-AM families in blue. (B) Alpha diversity of root fungi is similar between AM and non-AM plants. Fungal alpha diversity estimated by Shannon’s H index and Chao1 index. No overall differences were observed between AM and non-AM plants of for each site (ANOVA, *P* > 0.05).

**Supplementary Figure 6.**
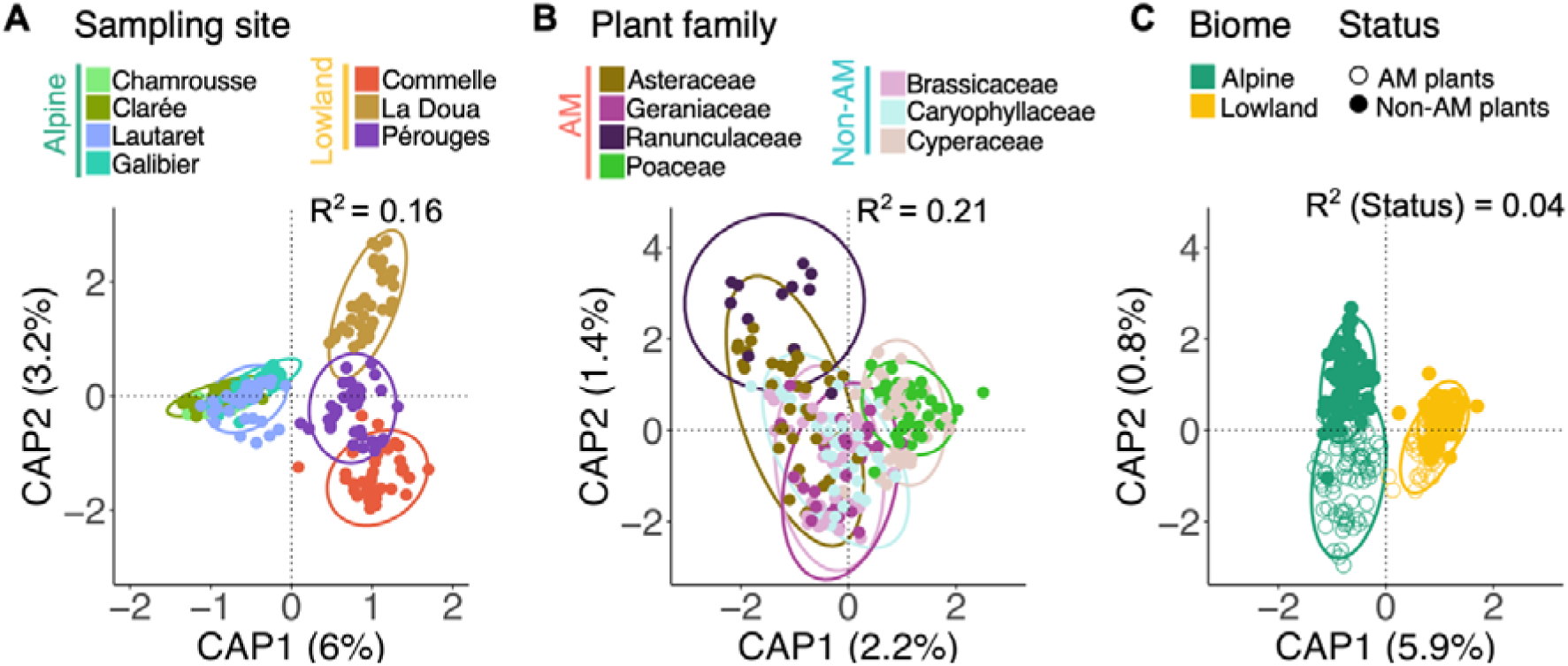
Root fungal microbiota differences between AM and non-AM plants, after excluding all Glomeromycota taxa. Differences in fungal communities across (A) sampling sites, (B) plant families, and (C) mycorrhizal groups within biomes are depicted using distance-based redundancy analysis (db-RDA) on Bray-Curtis (BC) dissimilarities. Ellipses show 95% dispersion surface of data. The variance explained by each factor used to constrain the analysis is indicated at the bottom (*R^2^*) (PerMANOVA, *P* < 0.05).

**Supplementary Figure 7.**
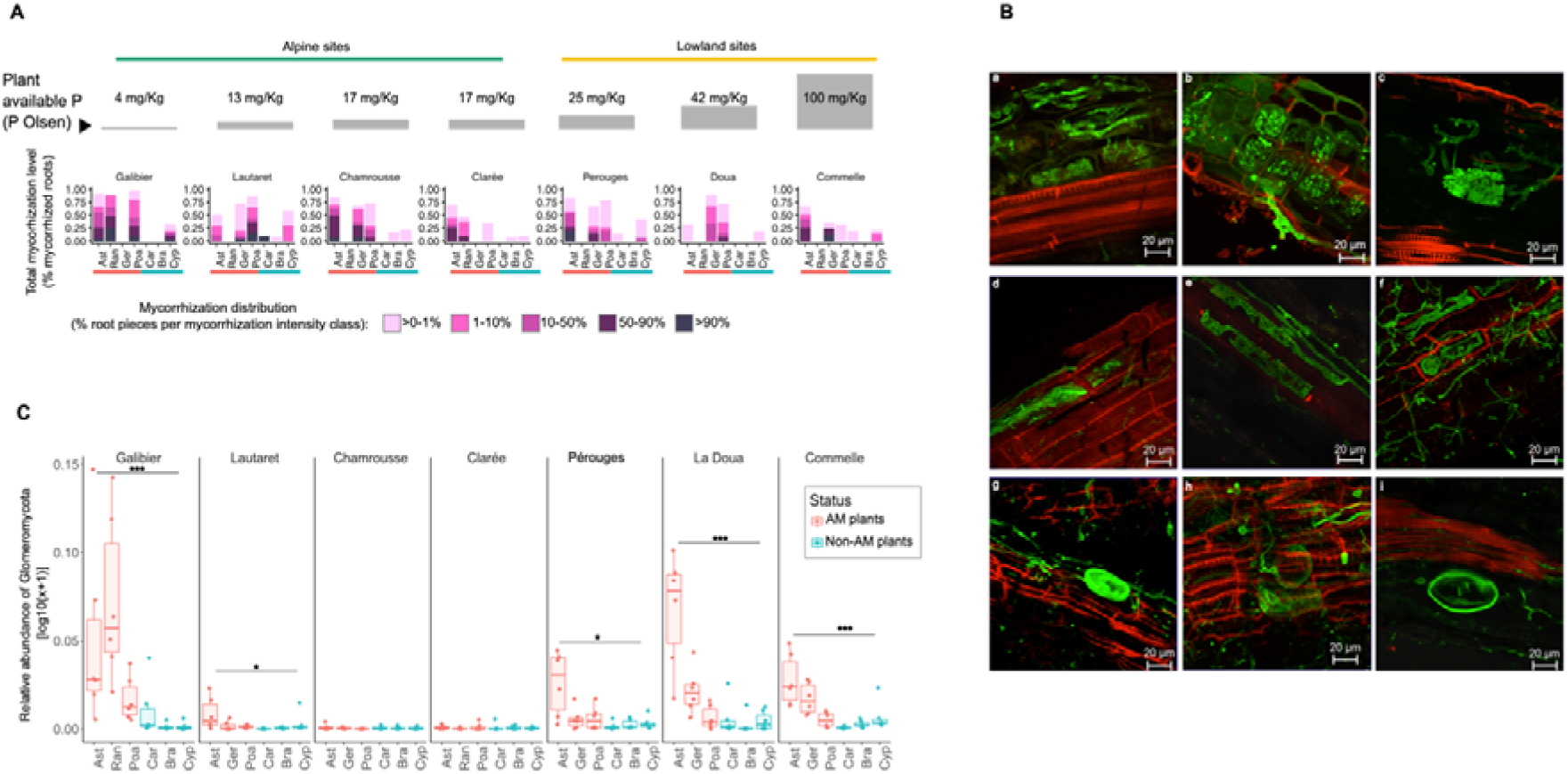
AM fungal colonization in the roots of AM and non-AM plants. (A) Mycorrhization levels measured across plant populations using trypan blue staining. Mycorrhization level (total percentage of mycorrhized roots) and mycorrhization distribution (percentage of roots per intensity class) were estimated using the method of Trouvelot *et al.* [62] scoring the presence of arbuscules and vesicles. From each condition approximately 10 fine root pieces from each individual plant were taken and pooled into a composite sample to asses mycorrhization levels within the plant population, 20-30 root pieces were analyzed for each condition. Abbreviations: Ast, Asteraceae; Ger, Geraniaceae; Poa, Poaceae; Car, Caryophyllaceae; Bra, Brassicaceae; Cyp, Cyperaceae; Ran, Ranunculaceae. (B) Intercellular or intracellular fungal structures observed by confocal microscopy after staining the fungal cell wall with WGA-Alexa (green) and the plant cell wall with propidium iodide (red). a-c, arbuscules observed in roots of AM plants *Taraxacum* sp., *Poa alpina*, and *Anemone baldensis*; d-f, coil-like structures observed in the roots of non-AM Cyperaceae *Carex sempervirens, Carex hirta*, and *Carex curvula*; g-i, vesicules observed in the roots of non-AM Caryophyllaceae, *Minuartia verna, Stellaria graminea*, and *Silene acaulis*. Scale bars are indicated for each photo. No arbuscules were observed in the roots of non-AM plant families. (C) Relative abundance of Glomeromycota in each root sample. Conditions are color coded according to mycorrhizal status (AM in pink et non-AM in blue). Asterisks indicate significant differences (Wilcoxon tests: *, *P* <0.05; ***, *P* < 0.001; NS, non-significant) between AM and non-AM plants within each site (Wilcoxon tests: *, *P* <0.05; ***, *P* < 0.001; NS, non-significant). Conditions are color-coded according to the plant mycorrhizal status (AM in pink et non-AM in blue).

**Supplementary Figure 8.**
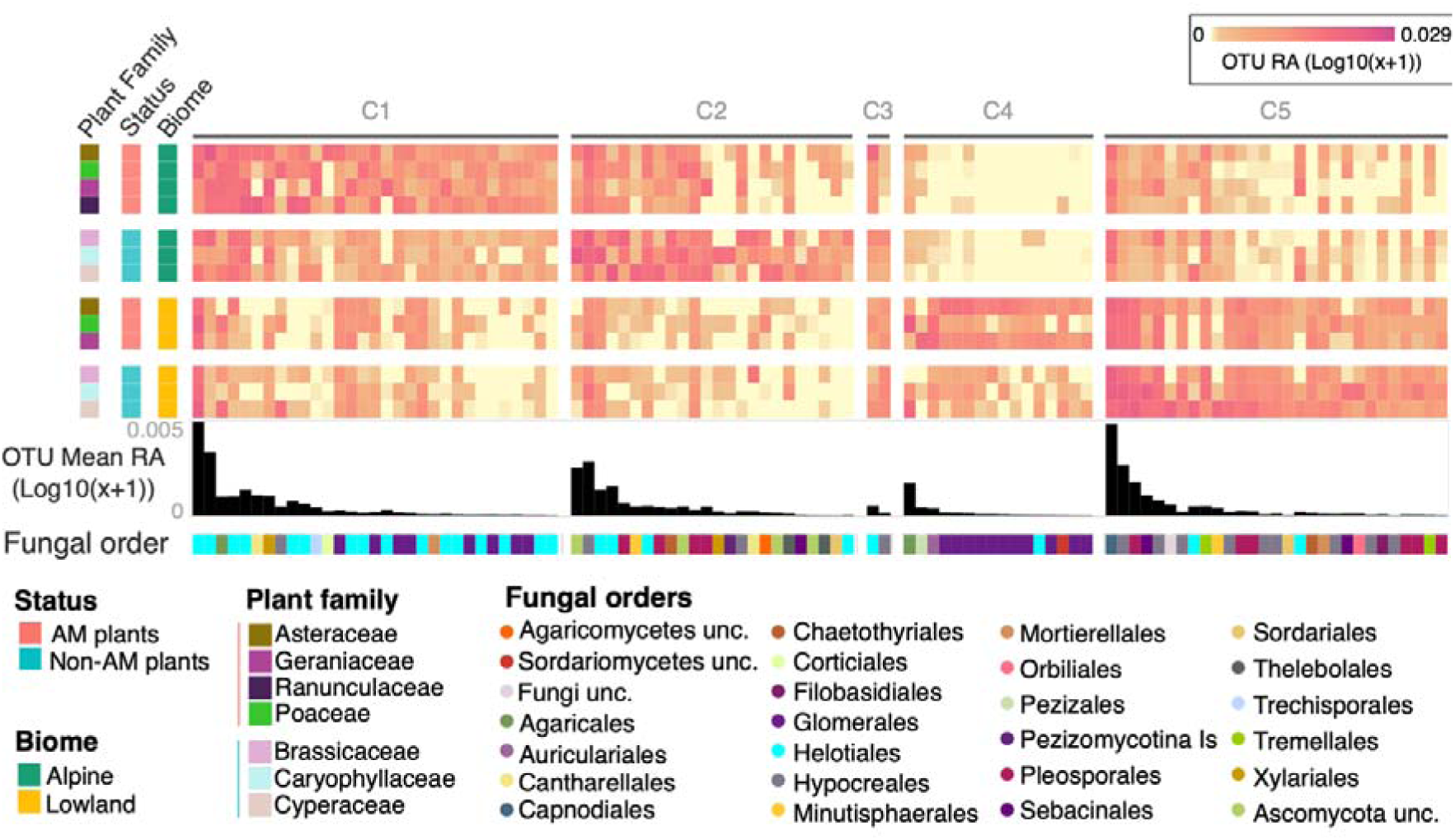
Fungal OTUs (104) contributing to root microbiota differences between AM and non-AM plants in alpine and lowland biomes. OTUs were identified using a mixed approach combining machine learning (Random Forest) and graph analysis (db-RDA) (Tab S5). Clusters of differentially abundant OTUs are indicated (C1-C6). The heatmap depicts OTU mean relative abundances per “family × site” combinations. Black barplots below show overall mean OTU relative abundance. OTU affiliation at the order level is indicated. Abbreviations: Unc., Unclassified.

**Supplementary Figure 9.**
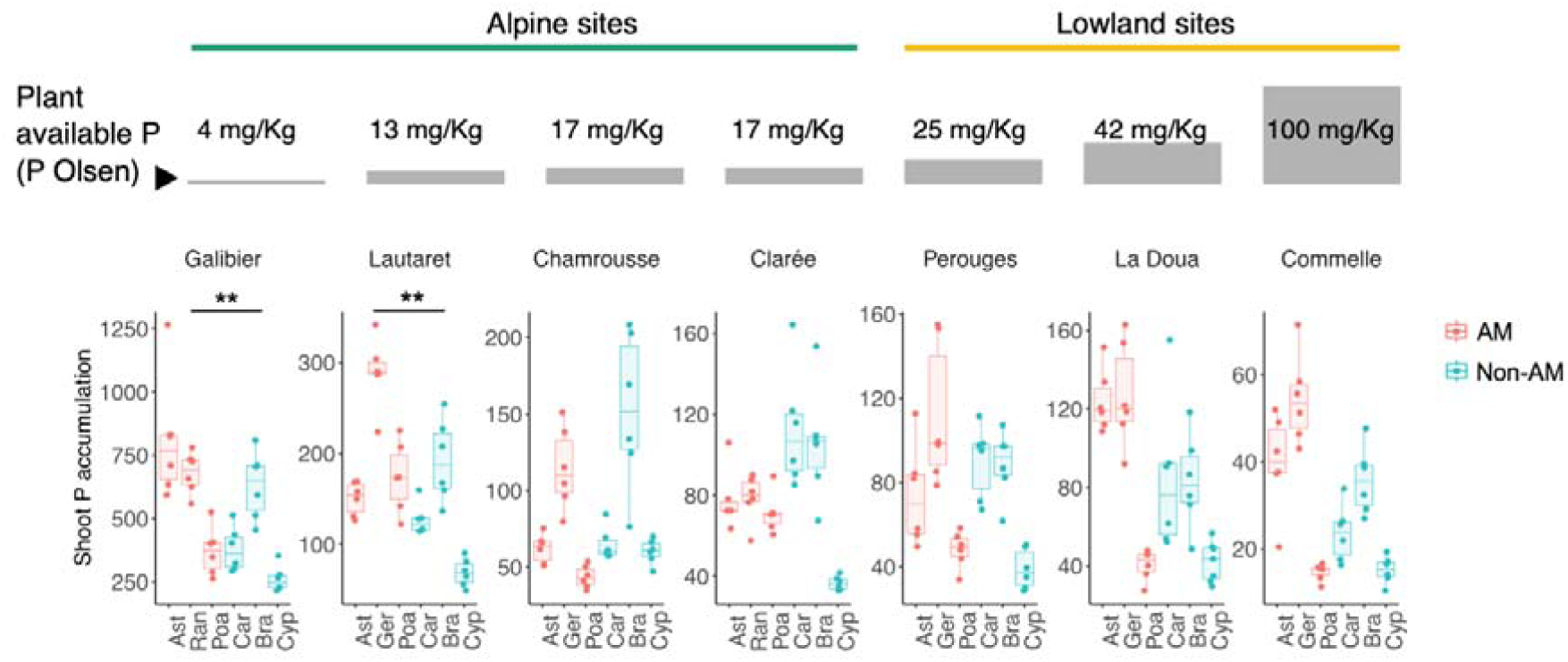
Shoot P-accumulation levels across sampling sites, plant families, and plant mycorrhizal status. P-accumulation in plant shoots was calculated as the ratio between Plant shoot P concentration (ICP optics) to plant-available soil P (Olsen; ICP optics). Sites are ordered according to levels of plant-available P in soil (P_Oslen_). Abbreviations: Ast., Asteraceae; Ger., Geraniaceae; Poa., Poaceae; Car., Caryophyllaceae; Bra., Brassicaceae; Cyp., Cyperaceae; Ran., Ranunculaceae. Asterisks indicate significant differences between AM and non-AM plants within sites (ANOVA, *P* < 0.01) (n = 6 plants per species per site).

**Supplementary Figure 10.**
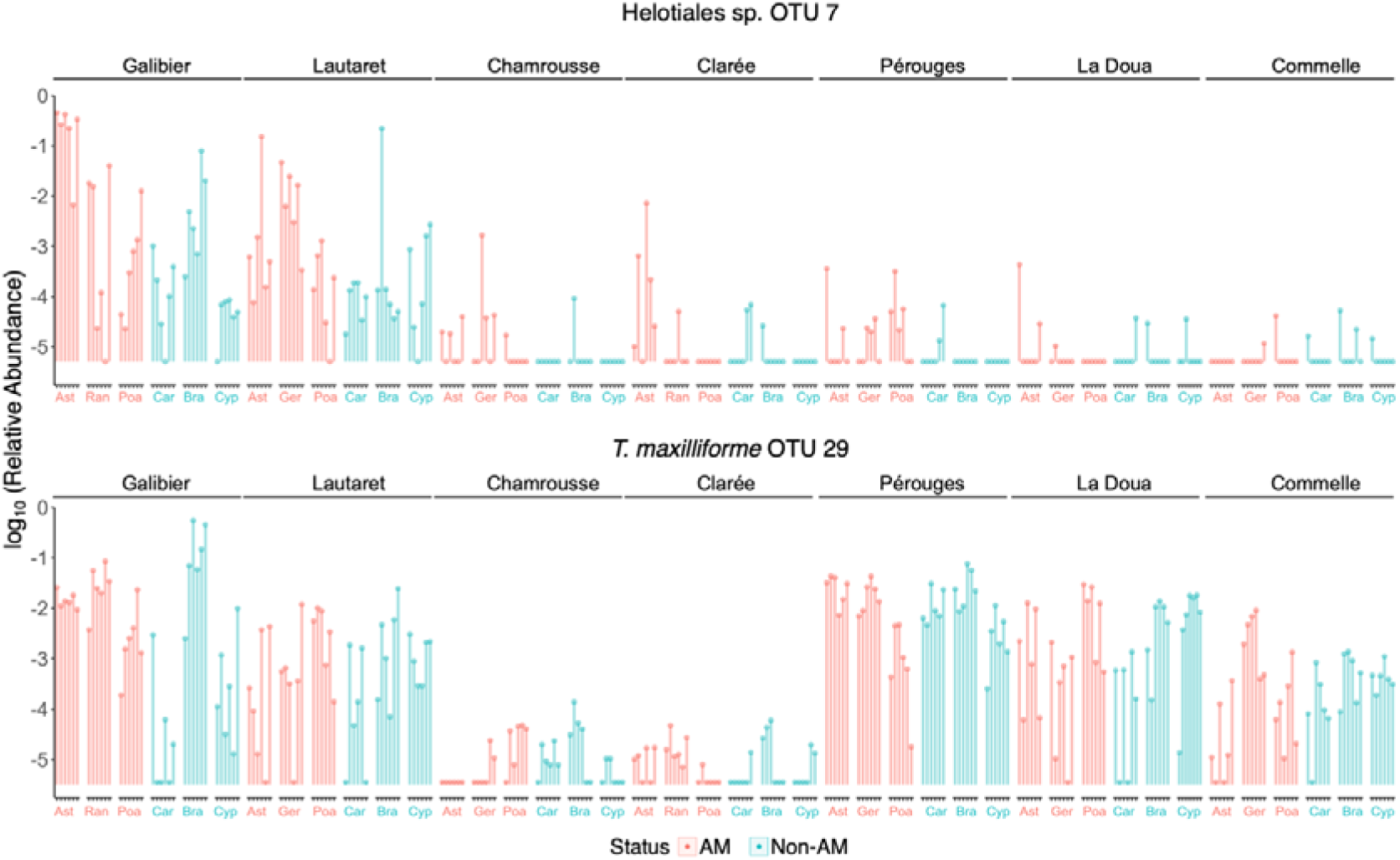
Relative abundance of OTU7 and OTU29 in the roots of AM (pink) and non-AM (blue) plants. Sites are ordered according to their soil available P (measured as P Olsen). Abbreviations: Ast, Asteraceae; Ger, Geraniaceae; Poa, Poaceae; Car, Caryophyllaceae; Bra, Brassicaceae; Cyp, Cyperaceae; Ran, Ranunculaceae.

**Supplementary Figure 11.**
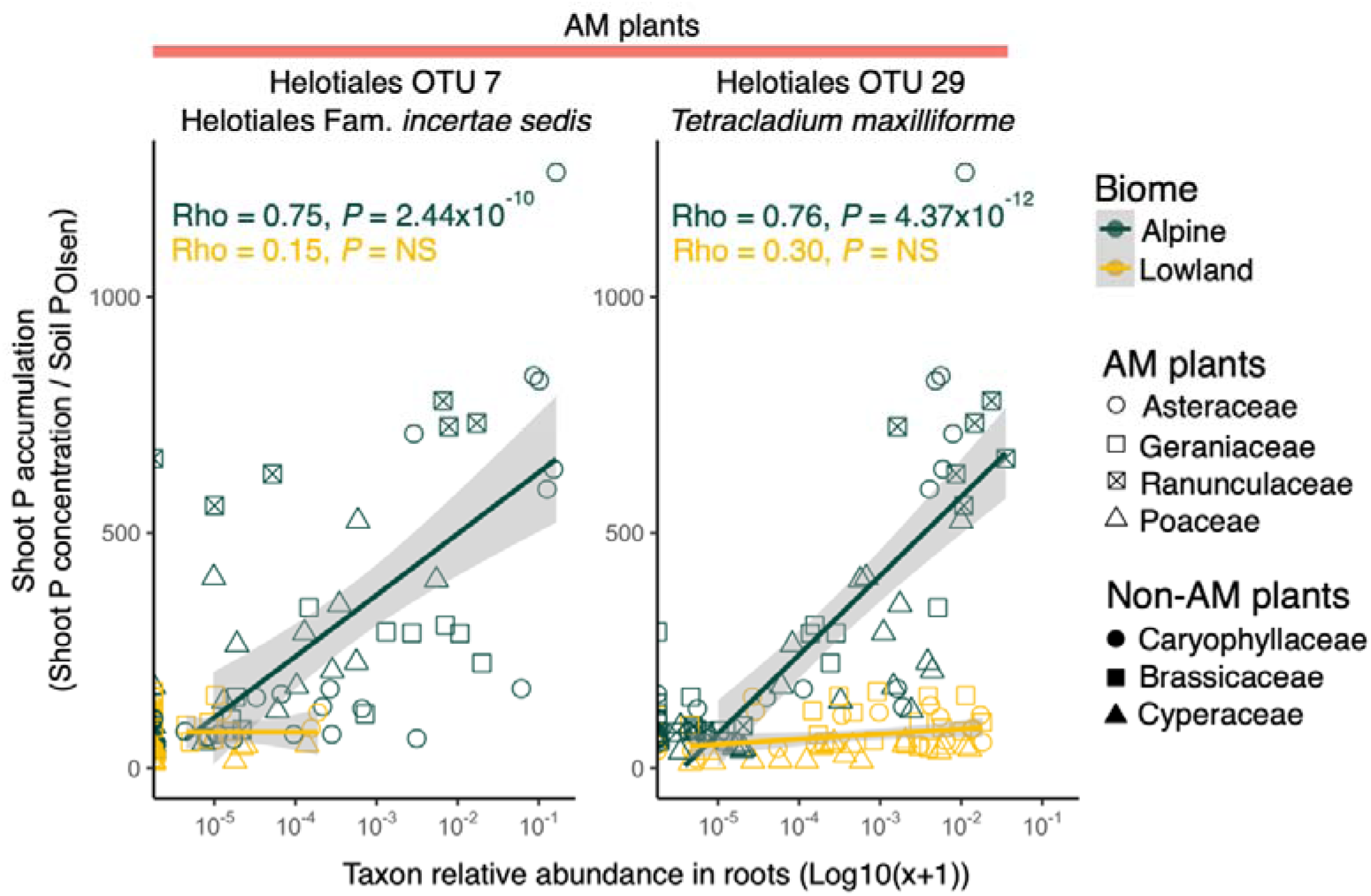
Correlations between shoot P-accumulation in AM plants and the relative abundance of Helotiales OTU7 and OTU29 in roots. Each point represents an individual plant. Correlations were calculated across all AM plants within each biome. Shoot P-accumulation was calculated as in Fig. 2. Spearman correlation coefficients (rho) and associated *P*-values are indicated for each condition.

**Supplementary Figure 12.**
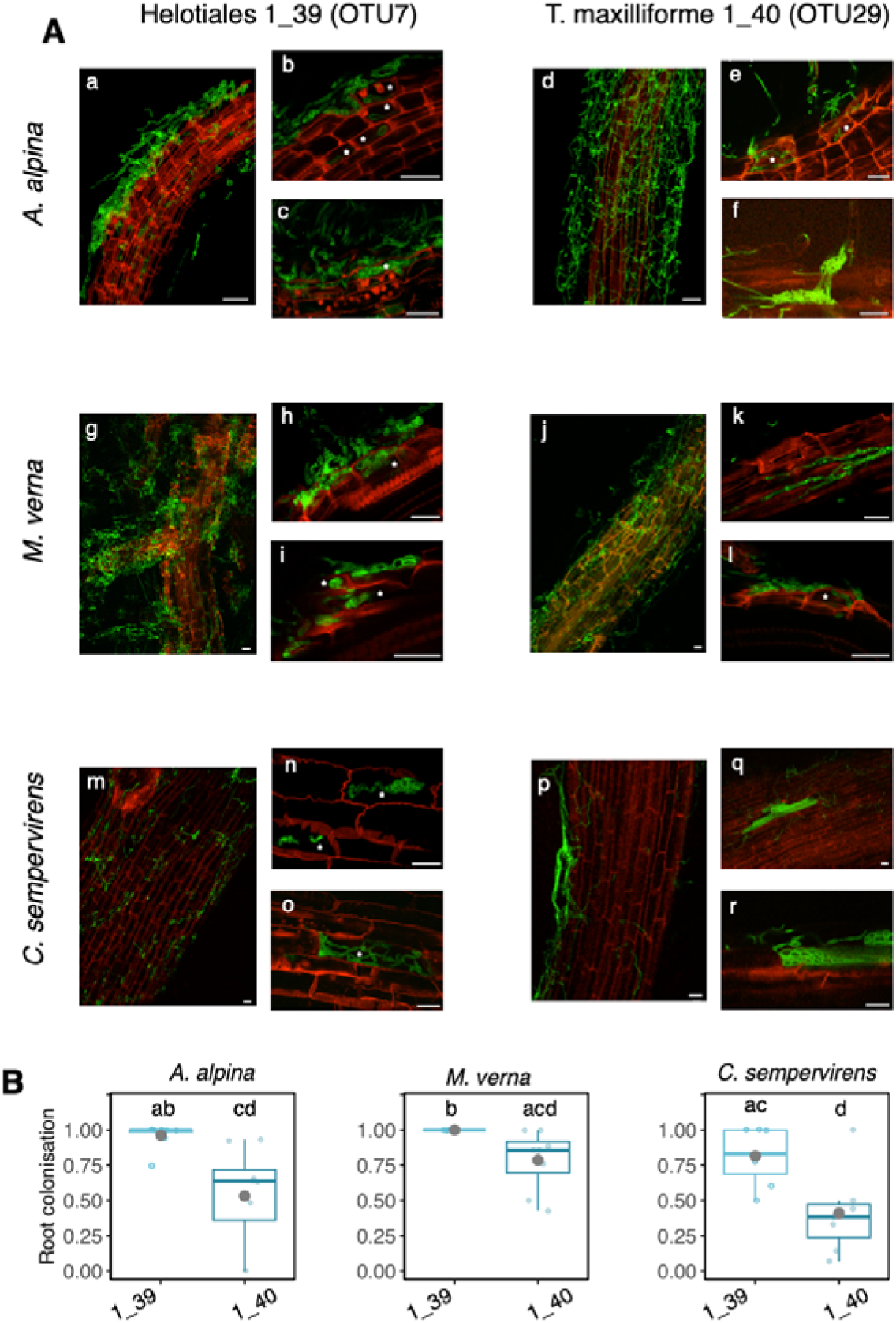
Root colonization by Helotiales 1_39 (OTU7) and *T. maxilliforme* 1_40 (OTU29) of diverse non-AM hosts in low-P soil microcosms. Plants were grown for 21 days in sterile sand-perlite-peat microcosms amended with 0.5X low-P Hoagland solution (100 µM P) and inoculated with isolate 1_39 or 1_40 (10^4^ propagules per plant) (see Fig. 8). (A) Examples of fungal root colonization visualized by confocal microscopy after staining the fungal cell wall (WGA-Alexa, green) and the plant cell wall (PI, red). Asterisks indicate intracellular colonization by the fungus, scale bars indicate 20 µm. (B) Fungal colonization rate per treatment estimated with the intersections method [68]. The experiment was done twice, each time 3-4 plants per treatment were randomly chosen to assess fungal colonization. Compiled results from both experiments are shown. All measures were done on individual plants (n = 7-8 plants per treatment). Different letters indicate significant differences between treatments (Wilcoxon test, *P* <0.05).

**Supplementary Figure 13.**
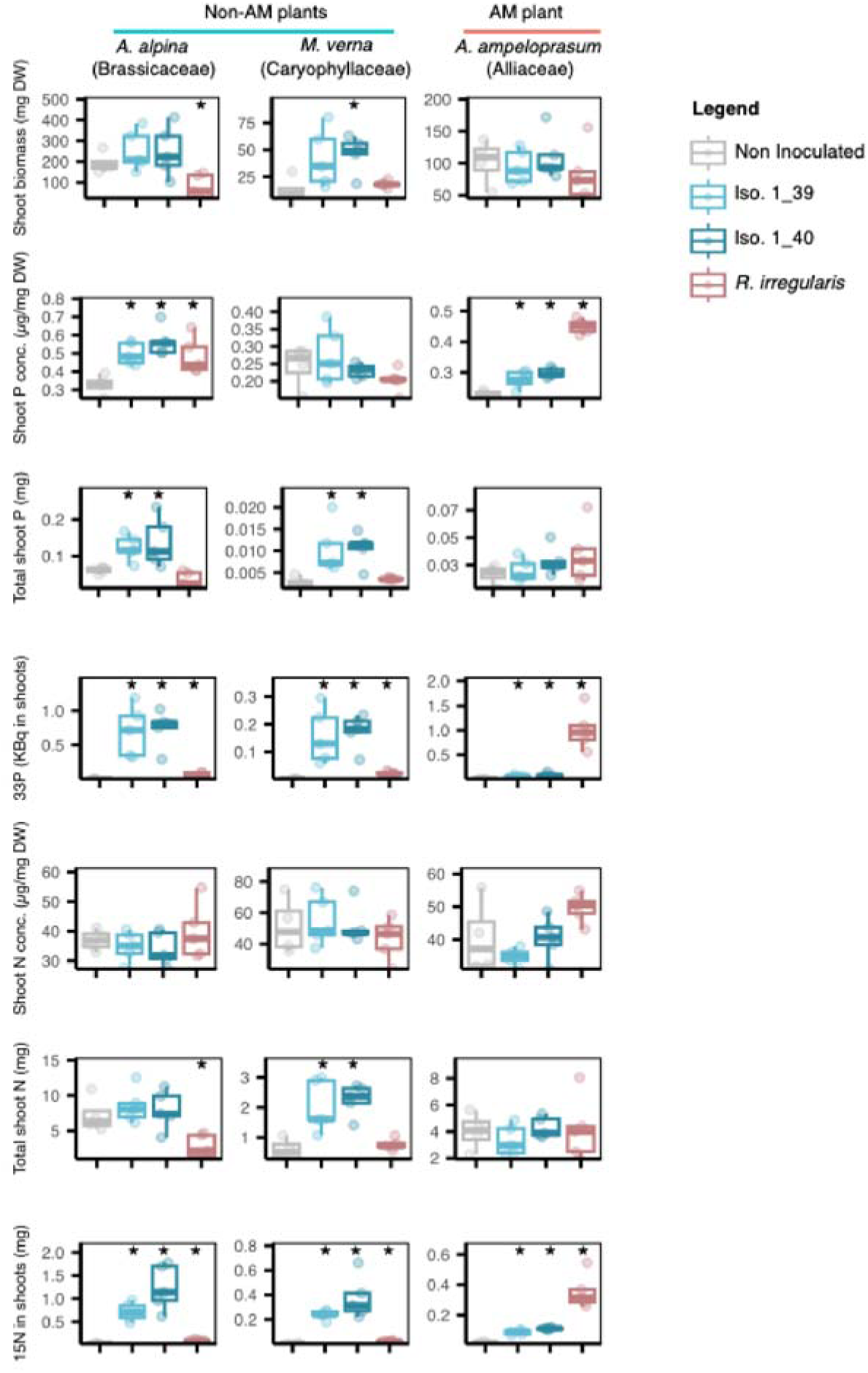
Effect of Helotiales 1_39 (OTU7) and *T. maxilliforme* 1_40 (OTU29) on plant growth and nutrition in the ^33^P/^15^N transfer experiment (Fig. 9). Non-AM *A. alpina* and *M. verna* as well as the model AM plant *A. ampeloprasum* were inoculated with Helotiales 1_39, *T. maxilliforme* 1_40, or the AM fungus *R. irregularis* (Ri) in low-P compartmentalized microcosms used to detect ^33^P and ^15^N transfer from fungi to plants (see Fig. 9). At the end of the experiment plant shoots were dried and used for measuring biomass and levels of P (Molybdate blue method), N (CN analyzer), ^33^P (scintillation counting), and ^15^N (IRMS). Each treatment included 4-5 microcosms, with 3-4 plants pooled per microcosm. Water was used in the non-inoculated (NI) treatment (n = 4-5). Asterisks indicate significant differences to the NI control (Wilcoxon test, *P* <0.05).

**Supplementary Figure 14.**
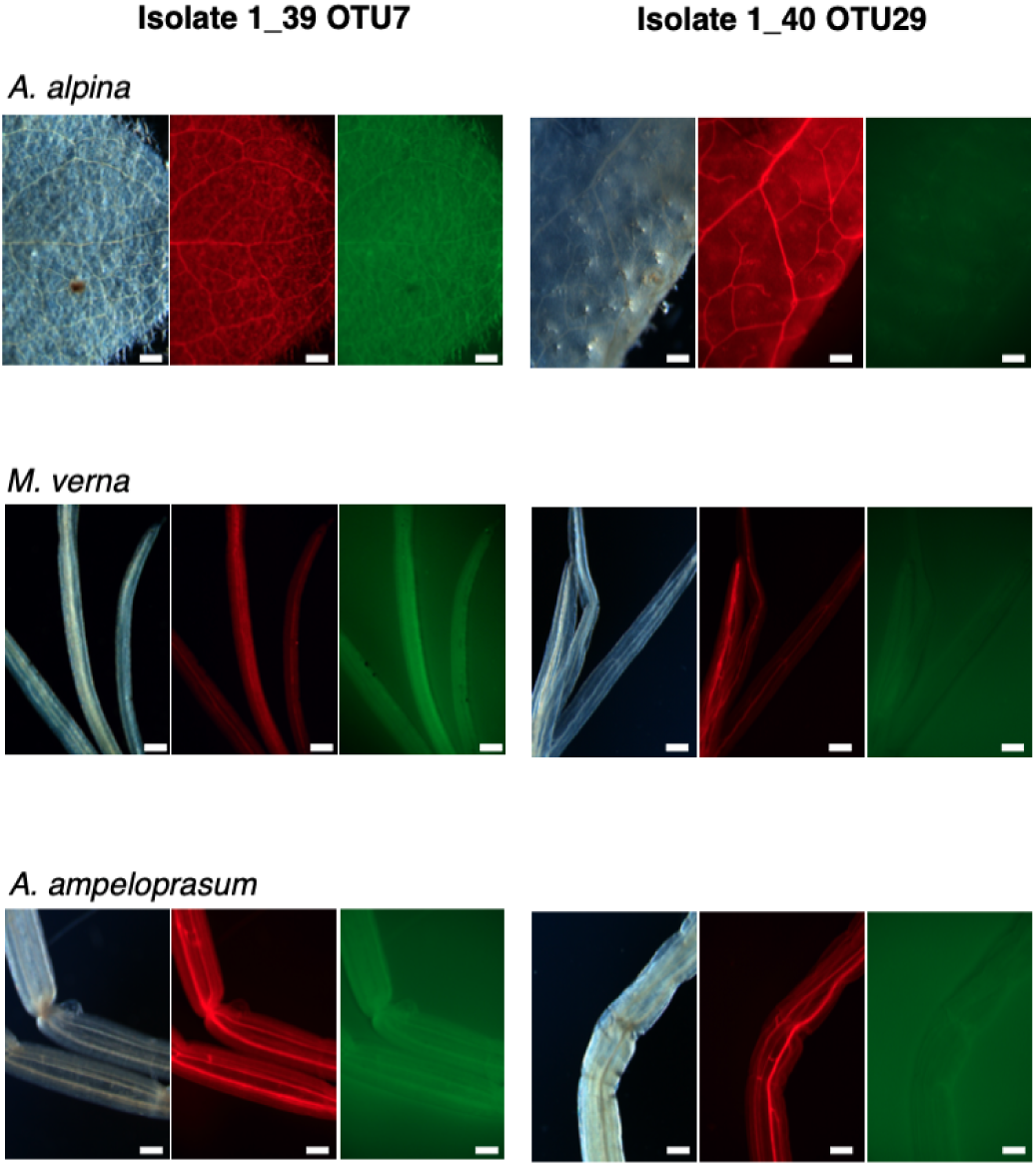
Control for fungal colonization of the plant shoot in the ^33^P/^15^N transfer experiment (Fig. 9). For each treatment, a fungus-inoculated plant was sampled, and the leaves were examined for signs of fungal colonization using a fluorescence binocular loupe. Fungal cell walls were stained with WGA-Alexa (green), while plant cell walls were labelled with propidium iodide (red). Observations revealed no fungal presence within the leaf tissues. Example pictures were taken at X40 magnification. The scale bar represents 1 mm.

## Supplementary tables

Table S1. Lists of genes and genomes used for the GEMOMA analysis of plant AM genes and the analysis results (Fig. 1).

Table S2. Soil physico-chemical characteristics of the seven sampling sites.

Table S3. Site environmental properties and plant species sampled for each family.

Table S4. Plant samples characteristics.

Table S5. Taxonomy and log10 (relative abundance +1) of each OTU associated either to AM or non-AM plants in each sample for each biome. An OTU can be present twice if it was enriched in AM or non-AM plants in each biome.

Table S6. Taxonomy, Correlation indication, and Relative abundance (log10(x+1)) of OTUs associated with higher plant P-accumulation for each sample.

Table S7. Mean out of bag error in RF classification models.

Table S8. Mean of squared residuals for each iteration in RF regression models.

## Notes

### Competing Interest Statement

The authors have declared no competing interest.

## References

1. Wang S, Ciais P, Reich PB, Cescatti A, Ellsworth DS, Janssens IA, et al. Phosphorus constrains global photosynthesis more than nitrogen does. Nat Ecol Evol. Nature Publishing Group; 2025;9:2025–35. 10.1038/s41559-025-02842-0

2. Holford ICR. Soil phosphorus: its measurement, and its uptake by plants. Soil Res. CSIRO PUBLISHING; 1997;35:227–40. 10.1071/s96047

3. Lambers H. Phosphorus Acquisition and Utilization in Plants. Annual Review of Plant Biology. Annual Reviews; 2022;73:17–42. 10.1146/annurev-arplant-102720-125738

4. Richardson AE, Barea J-M, McNeill AM, Prigent-Combaret C. Acquisition of phosphorus and nitrogen in the rhizosphere and plant growth promotion by microorganisms. Plant Soil. 2009;321:305–39. 10.1007/s11104-009-9895-2

5. Smith SE, Smith FA, Jakobsen I. Mycorrhizal fungi can dominate phosphate supply to plants irrespective of growth responses. Plant Physiol. 2003;133:16–20. 10.1104/pp.103.024380

6. Smith SE, Smith FA, Jakobsen I. Functional diversity in arbuscular mycorrhizal (AM) symbioses: the contribution of the mycorrhizal P uptake pathway is not correlated with mycorrhizal responses in growth or total P uptake. New Phytol. 2004;162:511–24. 10.1111/j.1469-8137.2004.01039.x

7. Whiteside MD, Werner GDA, Caldas VEA, Padje A van’t, Dupin SE, Elbers B, et al. Mycorrhizal Fungi Respond to Resource Inequality by Moving Phosphorus from Rich to Poor Patches across Networks. Current Biology. Elsevier; 2019;29:2043–2050.e8. 10.1016/j.cub.2019.04.061

8. Wang W, Shi J, Xie Q, Jiang Y, Yu N, Wang E. Nutrient Exchange and Regulation in Arbuscular Mycorrhizal Symbiosis. Molecular Plant. Elsevier; 2017;10:1147–58. 10.1016/j.molp.2017.07.012

9. Delaux P-M, Radhakrishnan GV, Jayaraman D, Cheema J, Malbreil M, Volkening JD, et al. Algal ancestor of land plants was preadapted for symbiosis. PNAS. National Academy of Sciences; 2015;112:13390–5. 10.1073/pnas.1515426112

10. Brundrett MC, Tedersoo L. Evolutionary history of mycorrhizal symbioses and global host plant diversity. New Phytol. 2018;220:1108–15. 10.1111/nph.14976

11. Werner GDA, Cornelissen JHC, Cornwell WK, Soudzilovskaia NA, Kattge J, West SA, et al. Symbiont switching and alternative resource acquisition strategies drive mutualism breakdown. Proc Natl Acad Sci USA. 2018;115:5229–34. 10.1073/pnas.1721629115

12. Radhakrishnan GV, Keller J, Rich MK, Vernié T, Mbadinga Mbadinga DL, Vigneron N, et al. An ancestral signalling pathway is conserved in intracellular symbioses-forming plant lineages. Nat Plants. 2020;6:280–9. 10.1038/s41477-020-0613-7

13. Cunning R, Silverstein RN, Baker AC. Investigating the causes and consequences of symbiont shuffling in a multi-partner reef coral symbiosis under environmental change. Proc R Soc Lond B Biol. Royal Society; 2015;282:20141725. 10.1098/rspb.2014.1725

14. Delaux P-M, Varala K, Edger PP, Coruzzi GM, Pires JC, Ané J-M. Comparative phylogenomics uncovers the impact of symbiotic associations on host genome evolution. PLOS Genet. Public Library of Science; 2014;10:e1004487. 10.1371/journal.pgen.1004487

15. Suzuki TA, Ley RE. The role of the microbiota in human genetic adaptation. Science. American Association for the Advancement of Science; 2020;370:eaaz6827. 10.1126/science.aaz6827

16. Petipas RH, Geber MA, Lau JA. Microbe-mediated adaptation in plants. Ecol Lett. 2021;24:1302–10.1111/ele.13755

17. Vandenkoornhuyse P, Quaiser A, Duhamel M, Le Van A, Dufresne A. The importance of the microbiome of the plant holobiont. New Phytologist. 2015;206:1196–206. 10.1111/nph.13312

18. Hiruma K, Gerlach N, Sacristán S, Nakano RT, Hacquard S, Kracher B, et al. Root endophyte Colletotrichum tofieldiae confers plant fitness benefits that are phosphate status dependent. Cell. 2016;165:464–74. 10.1016/j.cell.2016.02.028

19. Almario J, Jeena G, Wunder J, Langen G, Zuccaro A, Coupland G, et al. Root-associated fungal microbiota of nonmycorrhizal Arabis alpina and its contribution to plant phosphorus nutrition. Proc Natl Acad Sci USA. 2017;114:E9403–12. 10.1073/pnas.1710455114

20. Almario J, Fabiańska I, Saridis G, Bucher M. Unearthing the plant–microbe quid pro quo in root associations with beneficial fungi. New Phytol. 2022;234:1967–76. 10.1111/nph.18061

21. Yuan J, Wen T, Zhang H, Zhao M, Penton CR, Thomashow LS, et al. Predicting disease occurrence with high accuracy based on soil macroecological patterns of Fusarium wilt. ISME J. 2020;14:2936–50. 10.1038/s41396-020-0720-5

22. Chang H-X, Haudenshield JS, Bowen CR, Hartman GL. Metagenome-Wide Association Study and Machine Learning Prediction of Bulk Soil Microbiome and Crop Productivity. Front Microbiol [Internet]. Frontiers; 2017 [cited 2025 Nov 23];8. 10.3389/fmicb.2017.00519

23. Potato crop performance is predicted by tuber microbiome. Nat Microbiol. Nature Publishing Group; 2025;10:8–9. 10.1038/s41564-024-01893-6

24. Kemen E, Mahmoudi M, Hu Y, Almario J, Stincone P, Tenzer L-M, et al. Machine Learning Reveals Biocontrol Agents Shaping Disease Protection in Natural Arabidopsis Populations and Synthetic Communities [Internet]. Research Square; 2025 [cited 2025 Nov 23]. 10.21203/rs.3.rs-6130404/v1

25. Keilwagen J, Wenk M, Erickson JL, Schattat MH, Grau J, Hartung F. Using intron position conservation for homology-based gene prediction. Nucleic Acids Res. 2016;44:e89. 10.1093/nar/gkw092

26. Jin Y, Qian H. U.PhyloMaker: An R package that can generate large phylogenetic trees for plants and animals. Plant Divers. 2023;45:347–52. 10.1016/j.pld.2022.12.007

27. Sandaña P. Phosphorus uptake and utilization efficiency in response to potato genotype and phosphorus availability. European Journal of Agronomy. 2016;76:95–106. 10.1016/j.eja.2016.02.003

28. Bizabani C, Dames J. Effects of inoculating Lachnum and Cadophora isolates on the growth of Vaccinium corymbosum. Microbiol Res. 2015;181:68–74. 10.1016/j.micres.2015.08.005

29. Nguyen NH, Song Z, Bates ST, Branco S, Tedersoo L, Menke J, et al. FUNGuild: An open annotation tool for parsing fungal community datasets by ecological guild. Fungal Ecology. 2016;20:241–8. 10.1016/j.funeco.2015.06.006

30. Bock BM, Hoeksema JD, Johnson NC, Gehring CA. Evidence for common fungal networks among plants formed by a Dark Septate Endophyte in Sorghum bicolor. Commun Biol. Nature Publishing Group; 2025;8:996. 10.1038/s42003-025-08432-x

31. Bruyant P, Doré J, Vallon L, Moënne-Loccoz Y, Almario J. Needle in a Haystack: Culturing Plant-Beneficial Helotiales Lineages From Plant Roots. Environmental Microbiology. 2025;27:e70082. 10.1111/1462-2920.70082

32. Trautwig AN, Jackson MR, Kivlin SN, Stinson KA. Reviewing ecological implications of mycorrhizal fungal interactions in the Brassicaceae. Front Plant Sci [Internet]. Frontiers; 2023 [cited 2024 Nov 27];14. 10.3389/fpls.2023.1269815

33. Veiga RSL, Faccio A, Genre A, Pieterse CMJ, Bonfante P, van der Heijden MGA. Arbuscular mycorrhizal fungi reduce growth and infect roots of the non-host plant Arabidopsis thaliana. Plant Cell Environ. 2013;36:1926–37. 10.1111/pce.12102

34. Cardinal-McTeague WM, Sytsma KJ, Hall JC. Biogeography and diversification of Brassicales: A 103 million year tale. Mol Phylogenet Evol. 2016;99:204–24. 10.1016/j.ympev.2016.02.021

35. Yao G, Jin J-J, Li H-T, Yang J-B, Mandala VS, Croley M, et al. Plastid phylogenomic insights into the evolution of Caryophyllales. Mol Phylogenet Evol. 2019;134:74–86. 10.1016/j.ympev.2018.12.023

36. Elliott TL, Spalink D, Larridon I, Zuntini AR, Escudero M, Hackel J, et al. Global analysis of Poales diversification – parallel evolution in space and time into open and closed habitats. New Phytol. 2024;242:727–43. 10.1111/nph.19421

37. Hammond J, Broadley M, White P, King G, Bowen H, R H, et al. Shoot yield drives phosphorus use efficiency in Brassica oleracea and correlates with root architecture traits. J Exp Bot. J Exp Bot; 2009;60:1953–68. 10.1093/jxb/erp083

38. Hartman K, Schmid MW, Bodenhausen N, Bender SF, Valzano-Held AY, Schlaeppi K, et al. A symbiotic footprint in the plant root microbiome. Environ microbiome. 2023;18:65. 10.1186/s40793-023-00521-w

39. Song Y, Atza E, Sánchez-Gil JJ, Akkermans D, de Jonge R, de Rooij PGH, et al. Seed tuber microbiome can predict growth potential of potato varieties. Nat Microbiol. Nature Publishing Group; 2025;10:28–40. 10.1038/s41564-024-01872-x

40. Heklau H, Schindler N, Buscot F, Eisenhauer N, Ferlian O, Prada Salcedo LD, et al. Mixing tree species associated with arbuscular or ectotrophic mycorrhizae reveals dual mycorrhization and interactive effects on the fungal partners. Ecology and Evolution. 2021;11:5424–40. 10.1002/ece3.7437

41. Corona Ramírez A, Symanczik S, Gallusser T, Bodenhausen N. Quantification of arbuscular mycorrhizal fungi root colonization in wheat, tomato, and leek using absolute qPCR. Mycorrhiza. 2023;33:387–97. 10.1007/s00572-023-01122-8

42. Frew A. What does colonisation tell us? Revisiting the functional outcomes of root colonisation by arbuscular mycorrhizal fungi. New Phytologist [Internet]. 2025 [cited 2025 Dec 4];247:1572–8. 10.1111/nph.70284

43. Ihrmark K, Bödeker ITM, Cruz-Martinez K, Friberg H, Kubartova A, Schenck J, et al. New primers to amplify the fungal ITS2 region – evaluation by 454-sequencing of artificial and natural communities. FEMS Microbiol Ecol. 2012;82:666–77. 10.1111/j.1574-6941.2012.01437.x

44. Lavrinienko A, Jernfors T, Koskimäki JJ, Pirttilä AM, Watts PC. Does Intraspecific Variation in rDNA Copy Number Affect Analysis of Microbial Communities? Trends in Microbiology. Elsevier; 2021;29:19–27. 10.1016/j.tim.2020.05.019

45. Bruyant P, Moënne-Loccoz Y, Almario J. Root-associated Helotiales fungi: Overlooked players in plant nutrition. Soil Biol Biochem. 2024;191:109363. 10.1016/j.soilbio.2024.109363

46. Behie SW, Zelisko PM, Bidochka MJ. Endophytic Insect-Parasitic Fungi Translocate Nitrogen Directly from Insects to Plants. Science [Internet]. American Association for the Advancement of Science; 2012 [cited 2020 Jun 4];336:1576–7. 10.1126/science.1222289

47. Field KJ, Bidartondo MI, Rimington WR, Hoysted GA, Beerling David J, Cameron DD, et al. Functional complementarity of ancient plant–fungal mutualisms: contrasting nitrogen, phosphorus and carbon exchanges between Mucoromycotina and Glomeromycotina fungal symbionts of liverworts. New Phytologist [Internet]. 2019 [cited 2024 Oct 14];223:908–21. 10.1111/nph.15819

48. Peng L, Yang Y, Martin FM, Yuan Z. Endophytes with mycorrhizal potentials: biological and ecological implications. New Phytologist [Internet]. 2025 [cited 2025 Dec 4];nph.70772. 10.1111/nph.70772

49. Edgar RC. Muscle5: High-accuracy alignment ensembles enable unbiased assessments of sequence homology and phylogeny. Nat Commun. Nature Publishing Group; 2022;13:6968. 10.1038/s41467-022-34630-w

50. Nguyen L-T, Schmidt HA, von Haeseler A, Minh BQ. IQ-TREE: A Fast and Effective Stochastic Algorithm for Estimating Maximum-Likelihood Phylogenies. Molecular Biology and Evolution. 2015;32:268–74. 10.1093/molbev/msu300

51. Minh BQ, Schmidt HA, Chernomor O, Schrempf D, Woodhams MD, von Haeseler A, et al. IQ-TREE 2: New Models and Efficient Methods for Phylogenetic Inference in the Genomic Era. Mol Biol Evol. 2020;37:1530–4. 10.1093/molbev/msaa015

52. Murphy J, Riley JP. A modified single solution method for the determination of phosphate in natural waters. Analytica Chimica Acta. 1962;27:31–6. 10.1016/S0003-2670(00)88444-5

53. Gardes M, Bruns TD. ITS primers with enhanced specificity for basidiomycetes--application to the identification of mycorrhizae and rusts. Mol Ecol. 1993;2:113–8. 10.1111/j.1365-294x.1993.tb00005.x

54. Rohland N, Reich D. Cost-effective, high-throughput DNA sequencing libraries for multiplexed target capture. Genome Res. 2012;22:939–46. 10.1101/gr.128124.111

55. Schloss PD, Westcott SL, Ryabin T, Hall JR, Hartmann M, Hollister EB, et al. Introducing mothur: open-source, platform-independent, community-supported software for describing and comparing microbial communities. Appl Environ Microbiol. American Society for Microbiology; 2009;75:7537–41. 10.1128/AEM.01541-09

56. Martin M. Cutadapt removes adapter sequences from high-throughput sequencing reads. EMBnet J. 2011;17:10–2. 10.14806/ej.17.1.200

57. Bengtsson-Palme J, Ryberg M, Hartmann M, Branco S, Wang Z, Godhe A, et al. Improved software detection and extraction of ITS1 and ITS2 from ribosomal ITS sequences of fungi and other eukaryotes for analysis of environmental sequencing data. Methods Ecol Evol. 2013;4:914–9. 10.1111/2041-210X.12073

58. Nilsson RH, Larsson K-H, Taylor AFS, Bengtsson-Palme J, Jeppesen TS, Schigel D, et al. The UNITE database for molecular identification of fungi: handling dark taxa and parallel taxonomic classifications. Nucl Acids Res. 2019;47:D259–64. 10.1093/nar/gky1022

59. Abarenkov K, Zirk A, Piirmann T, Pöhönen R, Ivanov F, Nilsson RH, et al. UNITE mothur release for eukaryotes [Internet]. 2022 [cited 2023 Aug 30]. 10.15156/BIO/2483921

60. Species Fungorum - Species synonymy [Internet]. [cited 2024 Oct 10]. https://www.speciesfungorum.org/Names/SynSpecies.asp?RecordID=291360. Accessed 10 Oct 2024

61. Murashige T, Skoog F. A Revised Medium for Rapid Growth and Bio Assays with Tobacco Tissue Cultures. | EBSCOhost [Internet]. 1962 [cited 2024 Oct 10]. p. 473. 10.1111/j.1399-3054.1962.tb08052.x

62. Trouvelot A, Kough JL, Gianinazzi-Pearson V. Estimation of vesicular arbuscular mycorrhizal infection levels. Resarch for methods having a functionnal significance. Physiological and genetical aspects of mycorrhizae = Aspects physiologiques et genetiques des mycorhizes[]: proceedings of the 1st European Symposium on Mycorrhizae, Dijon, 1-5 July 1985 [Internet]. Paris[]: Institut national de le recherche agronomique, c1986.; 1986 [cited 2021 May 4]; https://agris.fao.org/agris-search/search.do?recordID=US201301430989. Accessed 4 May 2021

63. R Core Team. 2019. R: A language and environment for statistical computing. https://www.R-project.org/. [Internet]. R Foundation for Statistical Computing, Vienna, Austria.; https://www.R-project.org/.

64. Peterson RA, Cavanaugh JE. Ordered quantile normalization: a semiparametric transformation built for the cross-validation era. J Appl Stat. Taylor & Francis; 2020;47:2312–27. 10.1080/02664763.2019.1630372

65. McMurdie PJ, Holmes S. phyloseq: An R package for reproducible interactive analysis and graphics of microbiome census data. PLOS ONE. Public Library of Science; 2013;8:e61217. 10.1371/journal.pone.0061217

66. Oksanen J, Blanchet FG, Friendly M, Kindt R, Legendre P, McGlinn D, et al. Package “vegan”, community ecology package V.2.5.7. https://cran.r-project.org, https://github.com/vegandevs/vegan. 2020.

67. Breiman L, Cutler A, Taylor AFS, Wiener M. randomForest: Breiman and Cutlers Random Forests for Classification and Regression [Internet]. 2024 [cited 2024 Oct 10]. https://cran.r-project.org/web/packages/randomForest/index.html. Accessed 10 Oct 2024

68. McGONIGLE TP, Miller MH, Evans DG, Fairchild GL, Swan JA. A new method which gives an objective measure of colonization of roots by vesicular—arbuscular mycorrhizal fungi. New Phytologist. 1990;115:495–501. 10.1111/j.1469-8137.1990.tb00476.x

