## Supplementary Figures for "Helotiales fungi as potential nutritional partners for non-mycorrhizal plants: a machine learning and experimental approach"

**Fig. S1**

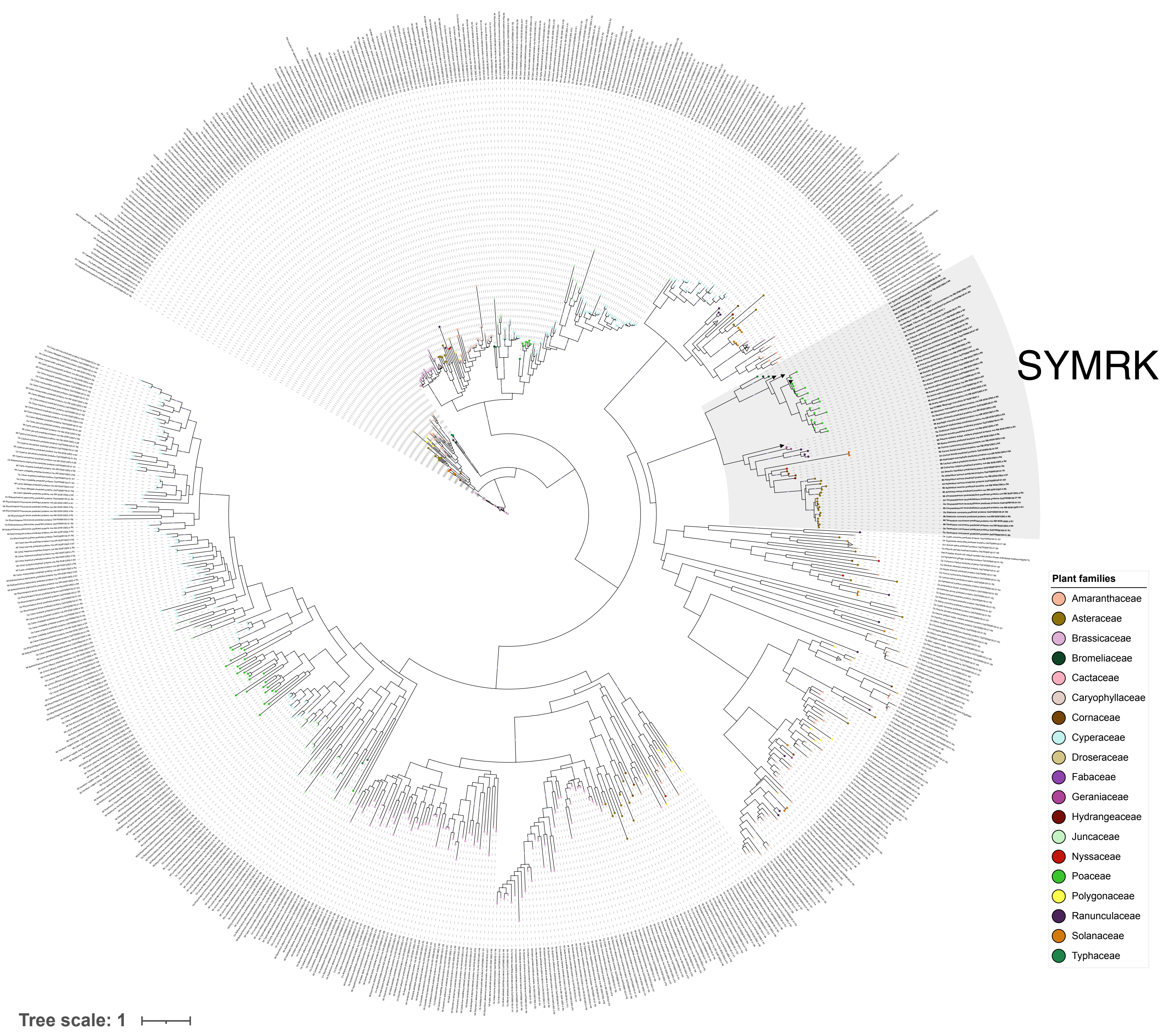
SYMRK
Plant families
Amaranthaceae
Asteraceae
Brassicaceae
Bromeliaceae
Cactaceae
Caryophyllaceae
Cornaceae
Cyperaceae
Droseraceae
Fabaceae
Geraniaceae
Hydrangeaceae
Juncaceae
Nyssaceae
Poaceae
Polygonaceae
Ranunculaceae
Solanaceae
Typhaceae
Tree scale: 1

**Fig. S2**

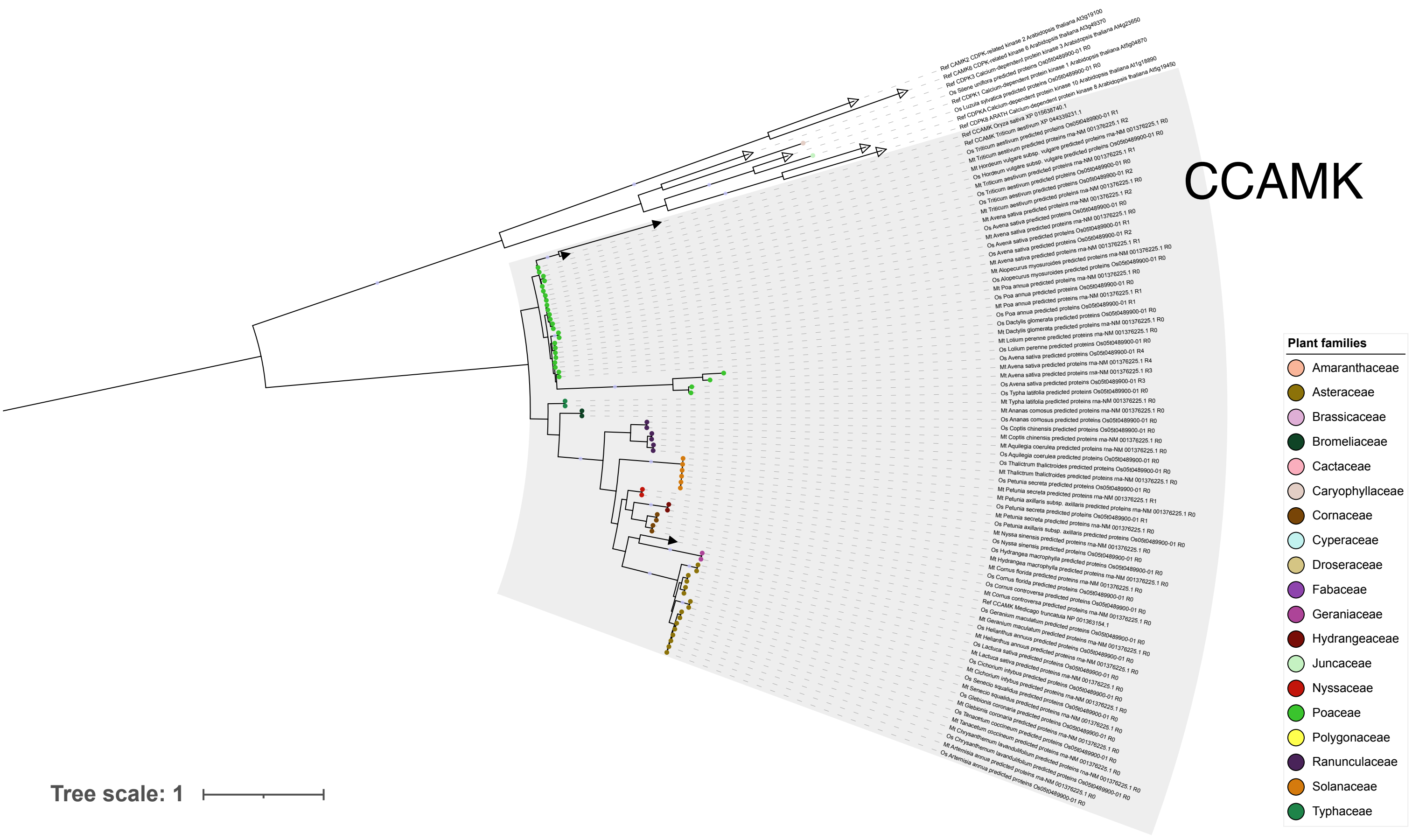
CCAMK
Plant families
Amaranthaceae
Asteraceae
Brassicaceae
Bromeliaceae
Cactaceae
Caryophyllaceae
Cornaceae
Cyperaceae
Droseraceae
Fabaceae
Geraniaceae
Hydrangeaceae
Juncaceae
Nyssaceae
Poaceae
Polygonaceae
Ranunculaceae
Solanaceae
Typhaceae
Tree scale: 1

**Fig. S3**

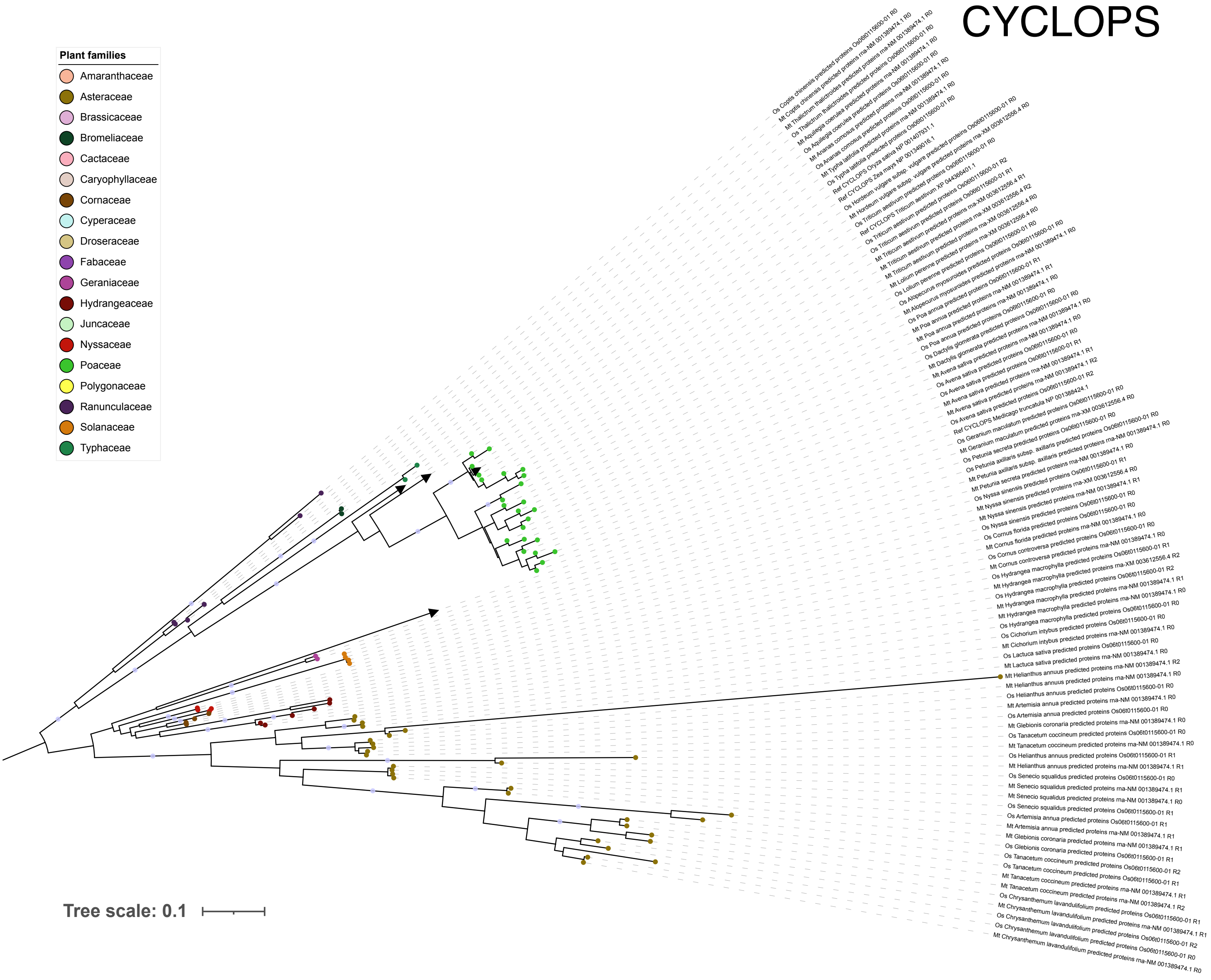
CYCLOPS
Plant families
Amaranthaceae
Asteraceae
Brassicaceae
Bromeliaceae
Cactaceae
Caryophyllaceae
Cornaceae
Cyperaceae
Droseraceae
Fabaceae
Geraniaceae
Hydrangeaceae
Juncaceae
Nyssaceae
Poaceae
Polygonaceae
Ranunculaceae
Solanaceae
Typhaceae
Tree scale: 0.1

#### Fig. S4

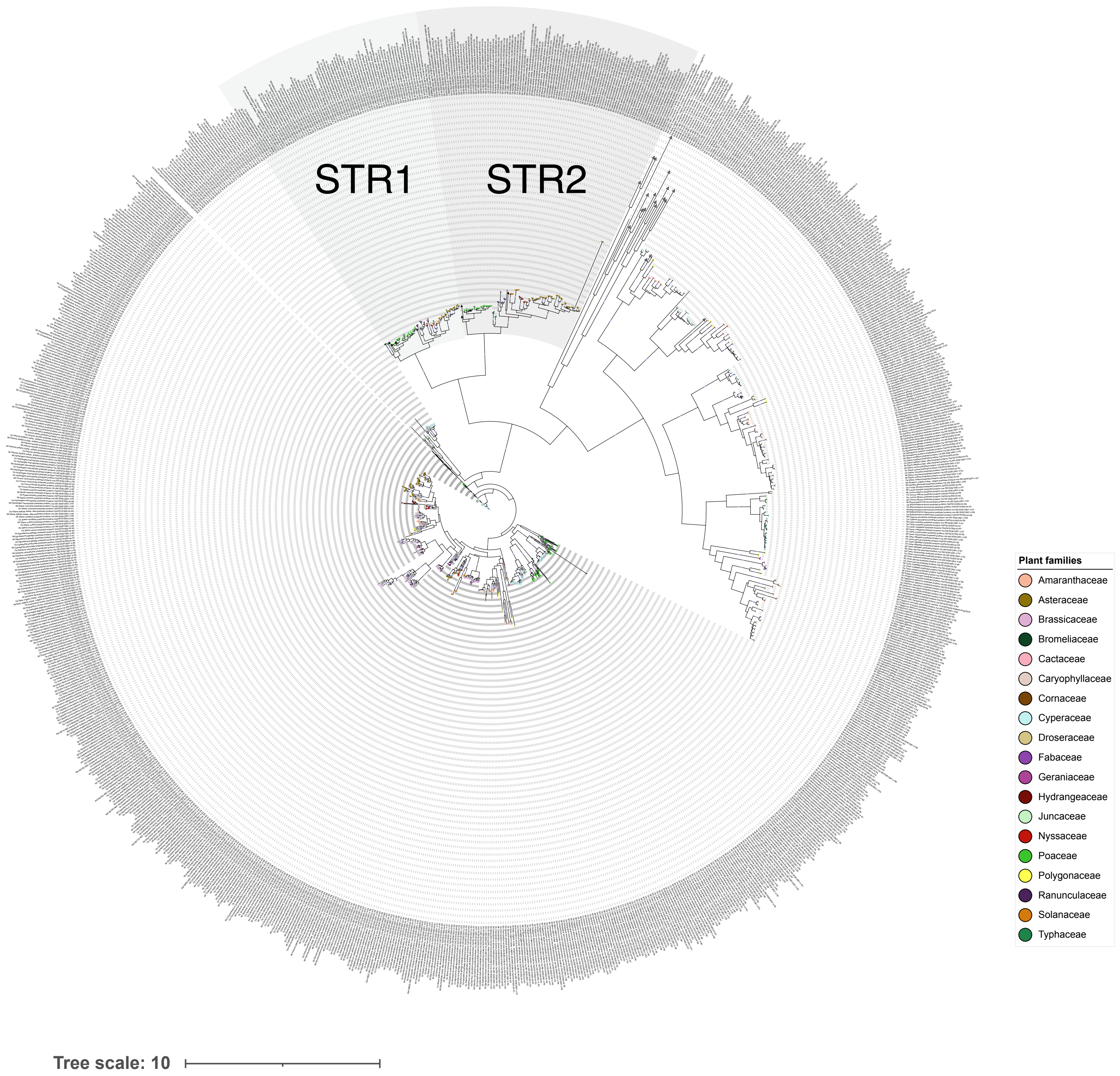
STR1
STR2
Plant families
Amaranthaceae
Asteraceae
Brassicaceae
Bromeliaceae
Cactaceae
Caryophyllaceae
Cornaceae
Cyperaceae
Droseraceae
Fabaceae
Geraniaceae
Hydrangeaceae
Juncaceae
Nyssaceae
Poaceae
Polygonaceae
Ranunculaceae
Solanaceae
Typhaceae
Tree scale: 10

**Fig. S5**

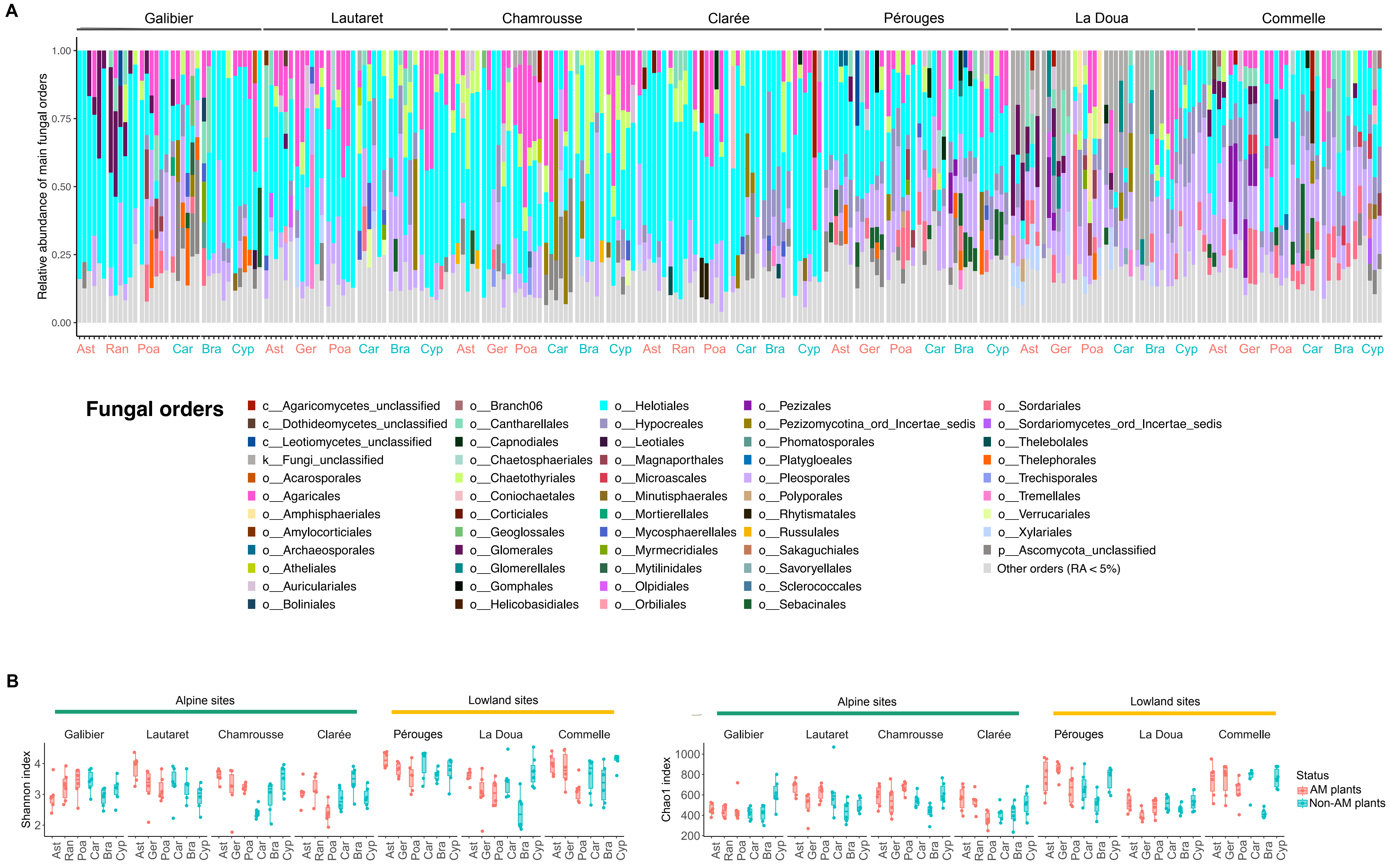
A
Galibier
Lautaret
Chamrousse
Clarée
Pérouges
La Doua
Commelle
Relative abundance of main fungal orders
1.00
0.75
0.50
0.25
0.00
Ast Ran Poa Car Bra Cyp Ast Ger Poa Car Bra Cyp Ast Ger Poa Car Bra Cyp Ast Ran Poa Car Bra Cyp Ast Ger Poa Car Bra Cyp Ast Ger Poa Car Bra Cyp Ast Ger Poa Car Bra Cyp
Fungal orders
c__Agaricomycetes_unclassified
c__Dothideomycetes_unclassified
c__Leotiomycetes_unclassified
k__Fungi_unclassified
o__Acarosporales
o__Agaricales
o__Amphisphaeriales
o__Amylocorticiales
o__Archaeosporales
o__Atheliales
o__Auriculariales
o__Boliniales
o__Branch06
o__Cantharellales
o__Capnodiales
o__Chaetosphaeriales
o__Chaetothyriales
o__Coniochaetales
o__Corticiales
o__Geoglossales
o__Glomerales
o__Glomerellales
o__Gomphales
o__Helicobasidiales
o__Helotiales
o__Hypocreales
o__Leotiales
o__Magnaporthales
o__Microascales
o__Minutisphaerales
o__Mortierellales
o__Mycosphaerellales
o__Myrmecridiales
o__Mytilinidales
o__Olpidiales
o__Orbiliales
o__Pezizales
o__Pezizomycotina_ord_Incertae_sedis
o__Phomatosporales
o__Platygloeales
o__Pleosporales
o__Polyporales
o__Rhytismatales
o__Russulales
o__Sakaguchiales
o__Savoryellales
o__Sclerococcales
o__Sebacinales
o__Sordariales
o__Sordariomycetes_ord_Incertae_sedis
o__Thelebolales
o__Thelephorales
o__Trechisporales
o__Tremellales
o__Verrucariales
o__Xylariales
p__Ascomycota_unclassified
Other orders (RA < 5%)
B
Alpine sites
Lowland sites
Galibier
Lautaret
Chamrousse
Clarée
Pérouges
La Doua
Commelle
Shannon index
4
3
2
Ast Ran Poa Car Bra Cyp Ast Ger Poa Car Bra Cyp Ast Ger Poa Car Bra Cyp Ast Ran Poa Car Bra Cyp Ast Ger Poa Car Bra Cyp Ast Ger Poa Car Bra Cyp Ast Ger Poa Car Bra Cyp
Chao1 index
1000
800
600
400
200
Status
AM plants
Non-AM plants

Fig.S6

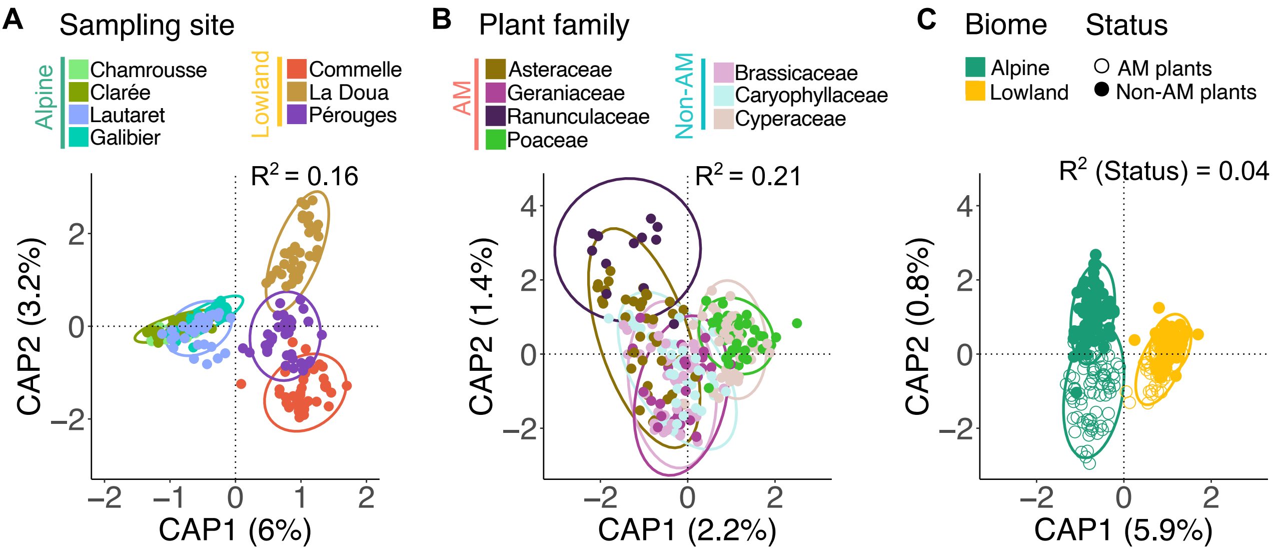

A
Sampling site
Alpine
Chamrousse
Clarée
Lautaret
Galibier
Lowland
Commelle
La Doua
Pérouges
R² = 0.16
CAP2 (3.2%)
CAP1 (6%)
B
Plant family
AM
Asteraceae
Geraniaceae
Ranunculaceae
Poaceae
Non-AM
Brassicaceae
Caryophyllaceae
Cyperaceae
R² = 0.21
CAP2 (1.4%)
CAP1 (2.2%)
C
Biome
Alpine
Lowland
Status
AM plants
Non-AM plants
R² (Status) = 0.04
CAP2 (0.8%)
CAP1 (5.9%)

Fig. S7

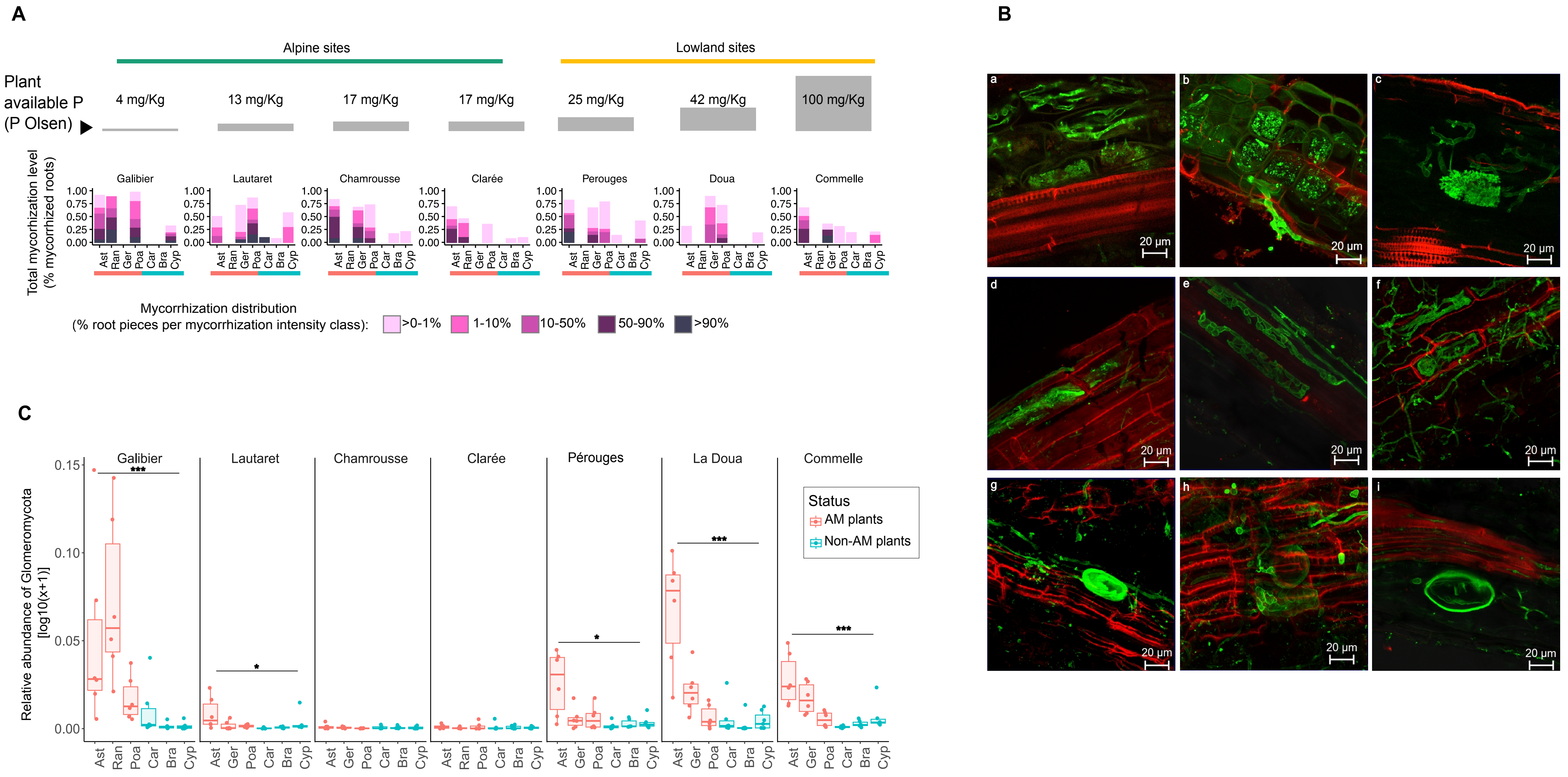

A
Alpine sites
Lowland sites
Plant available P (P Olsen)
4 mg/Kg
13 mg/Kg
17 mg/Kg
17 mg/Kg
25 mg/Kg
42 mg/Kg
100 mg/Kg
Total mycorhization level (% mycorrhized roots)
Galibier
Lautaret
Chamrousse
Clarée
Perouges
Doua
Commelle
Ast Ran Ger Poa Car Bra Cyp
Mycorrhization distribution (% root pieces per mycorrhization intensity class):
>0-1%
1-10%
10-50%
50-90%
>90%
B
20 µm
C
Relative abundance of Glomeromycota [log10(x+1)]
Galibier
Lautaret
Chamrousse
Clarée
Pérouges
La Doua
Commelle
Status
AM plants
Non-AM plants

### Fig. S8

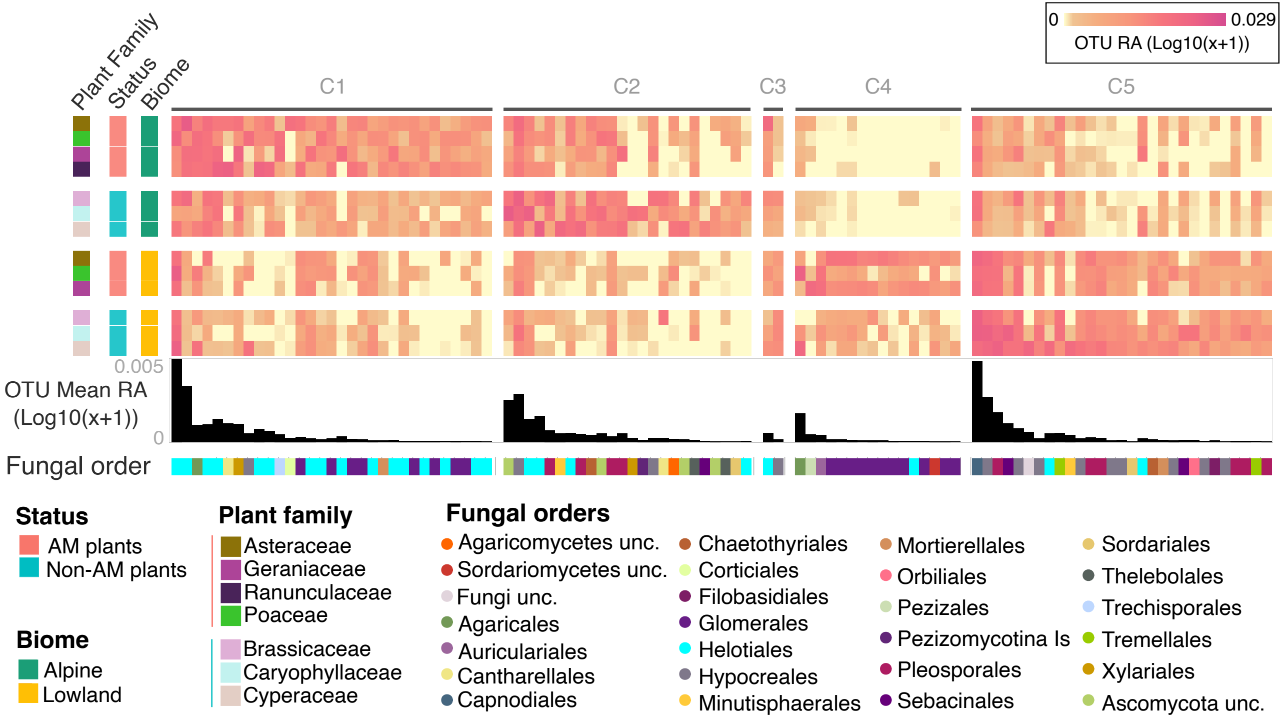

0
0.029
OTU RA (Log10(x+1))
Plant Family
Status
Biome
C1
C2
C3
C4
C5
0.005
OTU Mean RA
(Log10(x+1))
0
Fungal order
Status
AM plants
Non-AM plants
Biome
Alpine
Lowland
Plant family
Asteraceae
Geraniaceae
Ranunculaceae
Poaceae
Brassicaceae
Caryophyllaceae
Cyperaceae
Fungal orders
Agaricomycetes unc.
Sordariomycetes unc.
Fungi unc.
Agaricales
Auriculariales
Cantharellales
Capnodiales
Chaetothyriales
Corticiales
Filobasidiales
Glomerales
Helotiales
Hypocreales
Minutisphaerales
Mortierellales
Orbiliales
Pezizales
Pezizomycotina Is
Pleosporales
Sebacinales
Sordariales
Thelebolales
Trechisporales
Tremellales
Xylariales
Ascomycota unc.

### Fig. S9

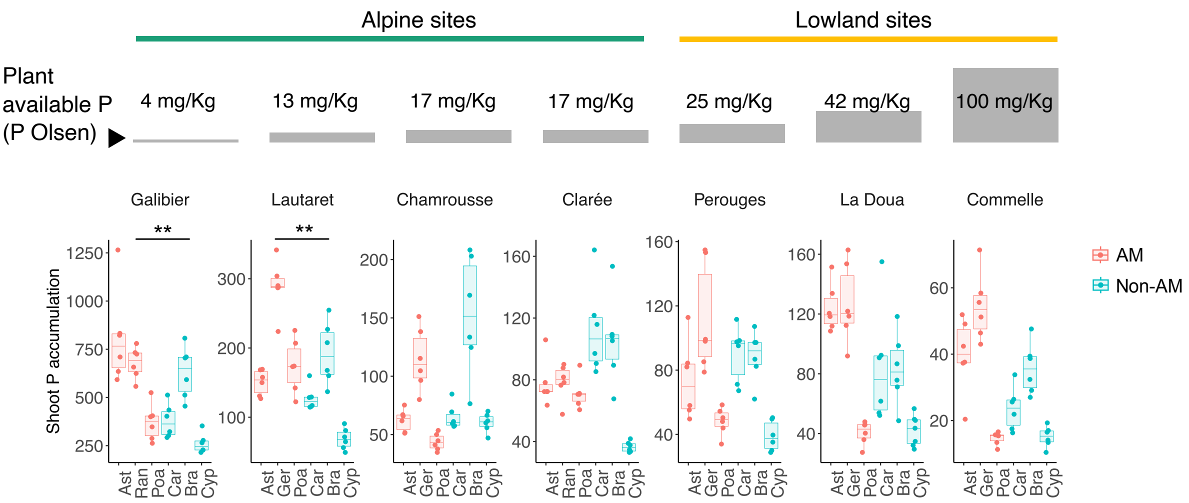

Alpine sites
Lowland sites
Plant available P (P Olsen)
4 mg/Kg
13 mg/Kg
17 mg/Kg
17 mg/Kg
25 mg/Kg
42 mg/Kg
100 mg/Kg
Galibier
Lautaret
Chamrousse
Clarée
Perouges
La Doua
Commelle
**
**
Shoot P accumulation
Ast
Ran
Ger
Poa
Car
Bra
Cyp
AM
Non-AM

**Fig. S10**

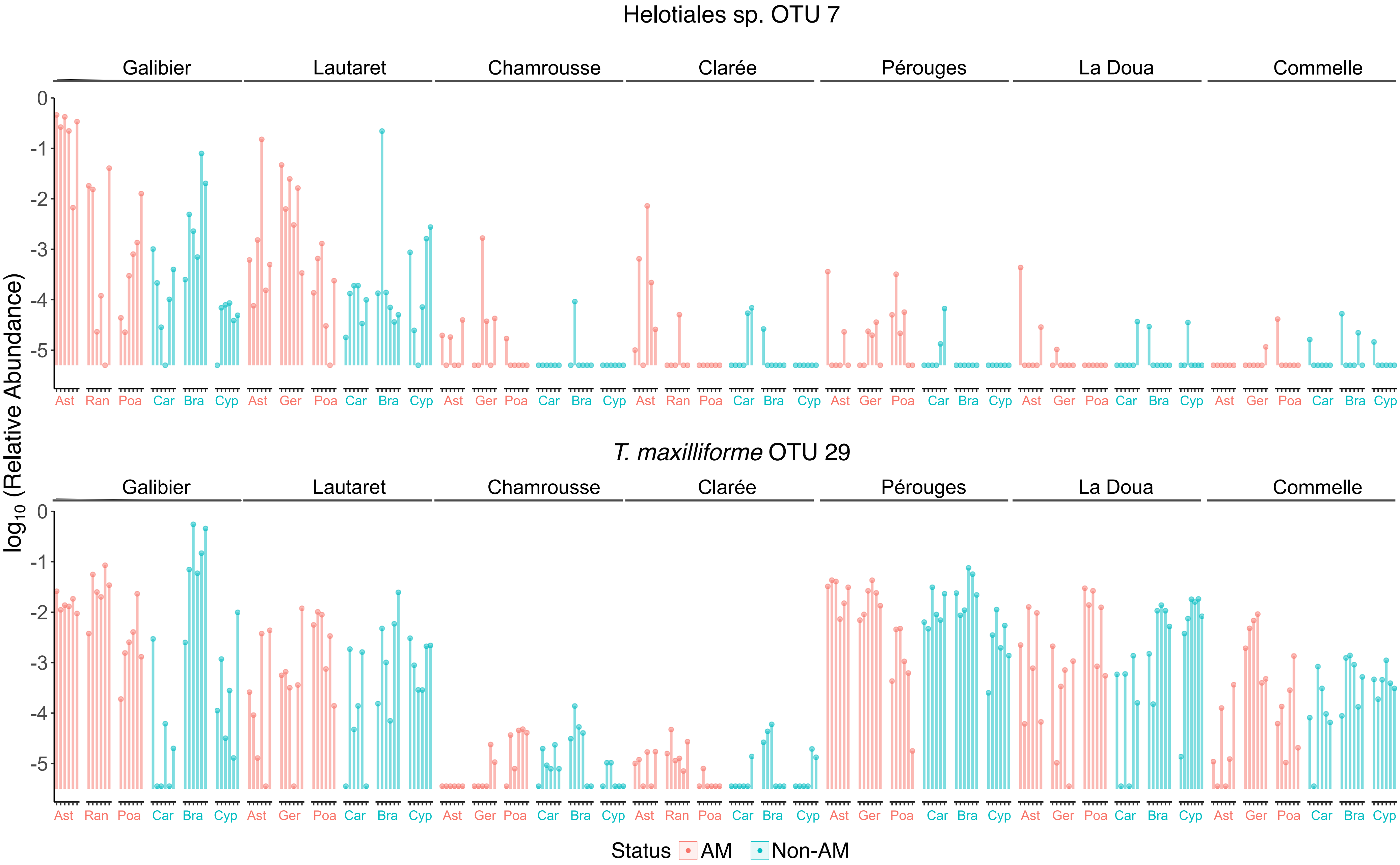
Helotiales sp. OTU 7
Galibier
Lautaret
Chamrousse
Clarée
Pérouges
La Doua
Commelle
0
-1
-2
-3
-4
-5
Ast Ran Poa Car Bra Cyp Ast Ger Poa Car Bra Cyp Ast Ger Poa Car Bra Cyp Ast Ran Poa Car Bra Cyp Ast Ger Poa Car Bra Cyp Ast Ger Poa Car Bra Cyp Ast Ger Poa Car Bra Cyp
T. maxilliforme OTU 29
Galibier
Lautaret
Chamrousse
Clarée
Pérouges
La Doua
Commelle
0
-1
-2
-3
-4
-5
Ast Ran Poa Car Bra Cyp Ast Ger Poa Car Bra Cyp Ast Ger Poa Car Bra Cyp Ast Ran Poa Car Bra Cyp Ast Ger Poa Car Bra Cyp Ast Ger Poa Car Bra Cyp Ast Ger Poa Car Bra Cyp
log10 (Relative Abundance)
Status AM Non-AM

**Fig. S11**

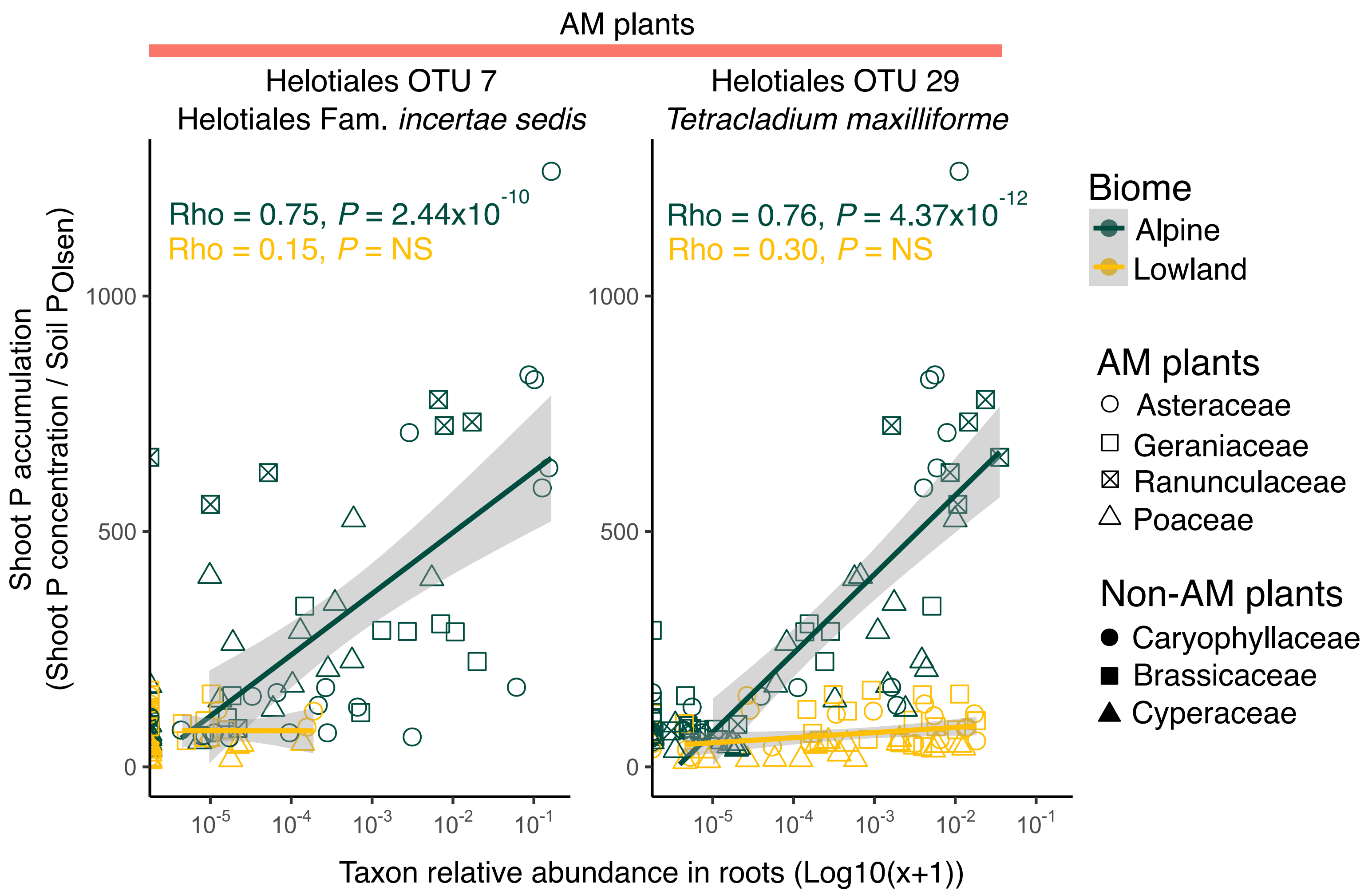

AM plants
Helotiales OTU 7
Helotiales Fam. incertae sedis
Helotiales OTU 29
Tetracladium maxilliforme
Rho = 0.75, P = 2.44x10^-10
Rho = 0.15, P = NS
Rho = 0.76, P = 4.37x10^-12
Rho = 0.30, P = NS
Shoot P accumulation
(Shoot P concentration / Soil P Olsen)
1000
500
0
10^-5
10^-4
10^-3
10^-2
10^-1
Taxon relative abundance in roots (Log10(x+1))
Biome
Alpine
Lowland
AM plants
Asteraceae
Geraniaceae
Ranunculaceae
Poaceae
Non-AM plants
Caryophyllaceae
Brassicaceae
Cyperaceae

Fig. S12

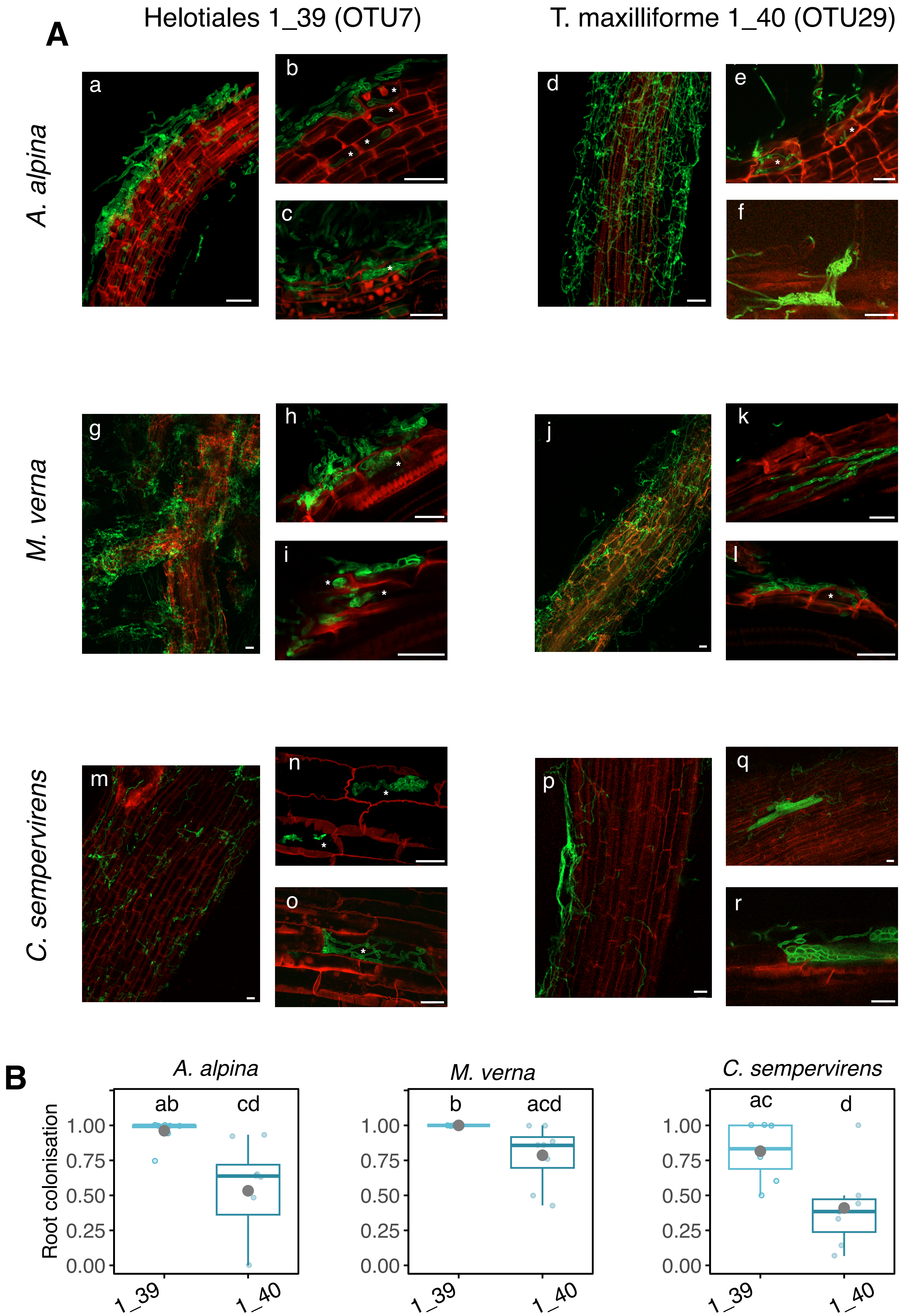
A
Helotiales 1_39 (OTU7)
T. maxilliforme 1_40 (OTU29)
A. alpina
M. verna
C. sempervirens
a
b
c
d
e
f
g
h
i
j
k
l
m
n
o
p
q
r
B
A. alpina
M. verna
C. sempervirens
Root colonisation
1.00
0.75
0.50
0.25
0.00
ab
cd
b
acd
ac
d
1_39
1_40

**Fig. S13**

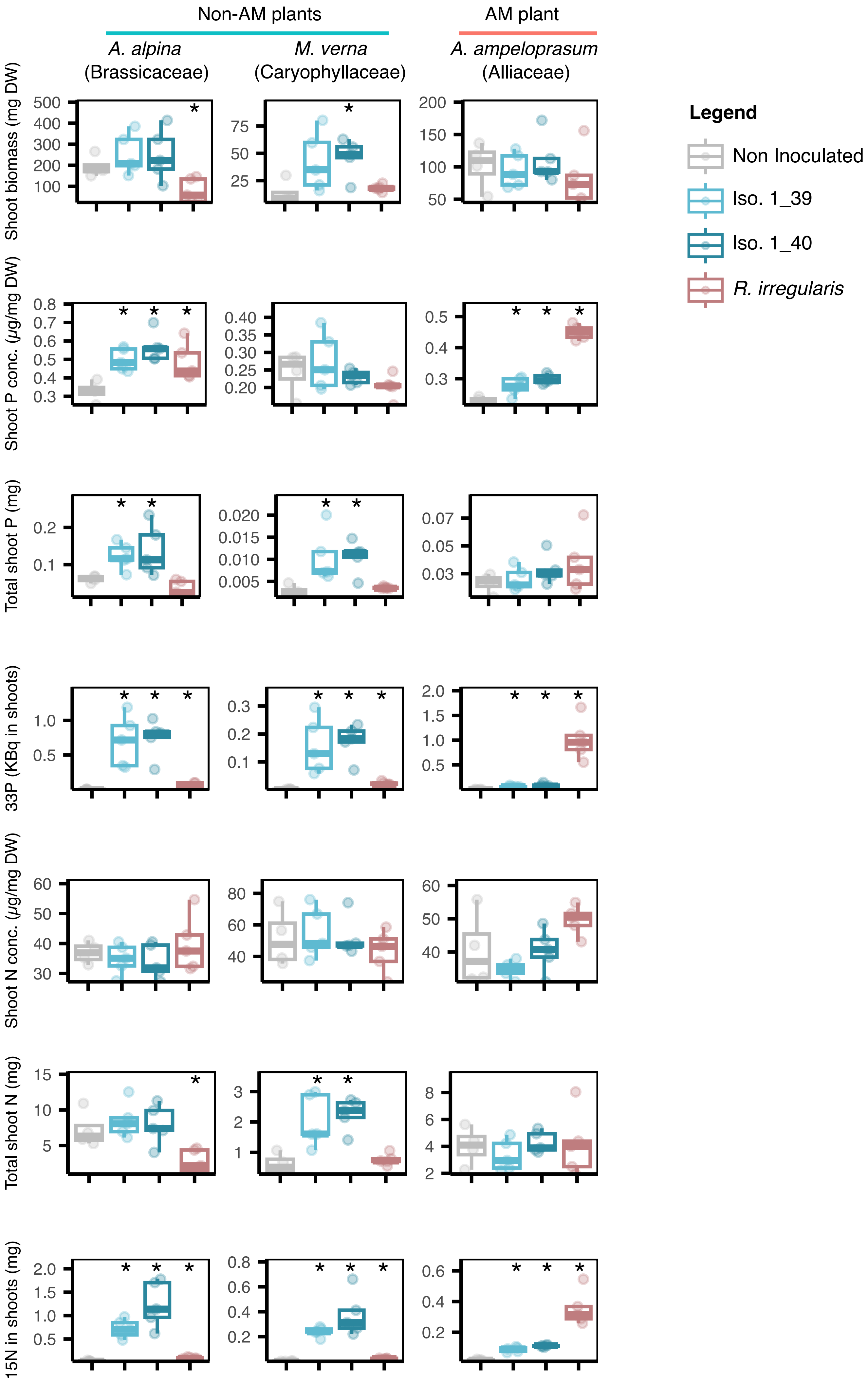
Non-AM plants
AM plant
*A. alpina*
(Brassicaceae)
*M. verna*
(Caryophyllaceae)
*A. ampeloprasum*
(Alliaceae)
Shoot biomass (mg DW)
Shoot P conc. (µg/mg DW)
Total shoot P (mg)
33P (KBq in shoots)
Shoot N conc. (µg/mg DW)
Total shoot N (mg)
15N in shoots (mg)
Legend
Non Inoculated
Iso. 1_39
Iso. 1_40
*R. irregularis*

**Fig. S14**

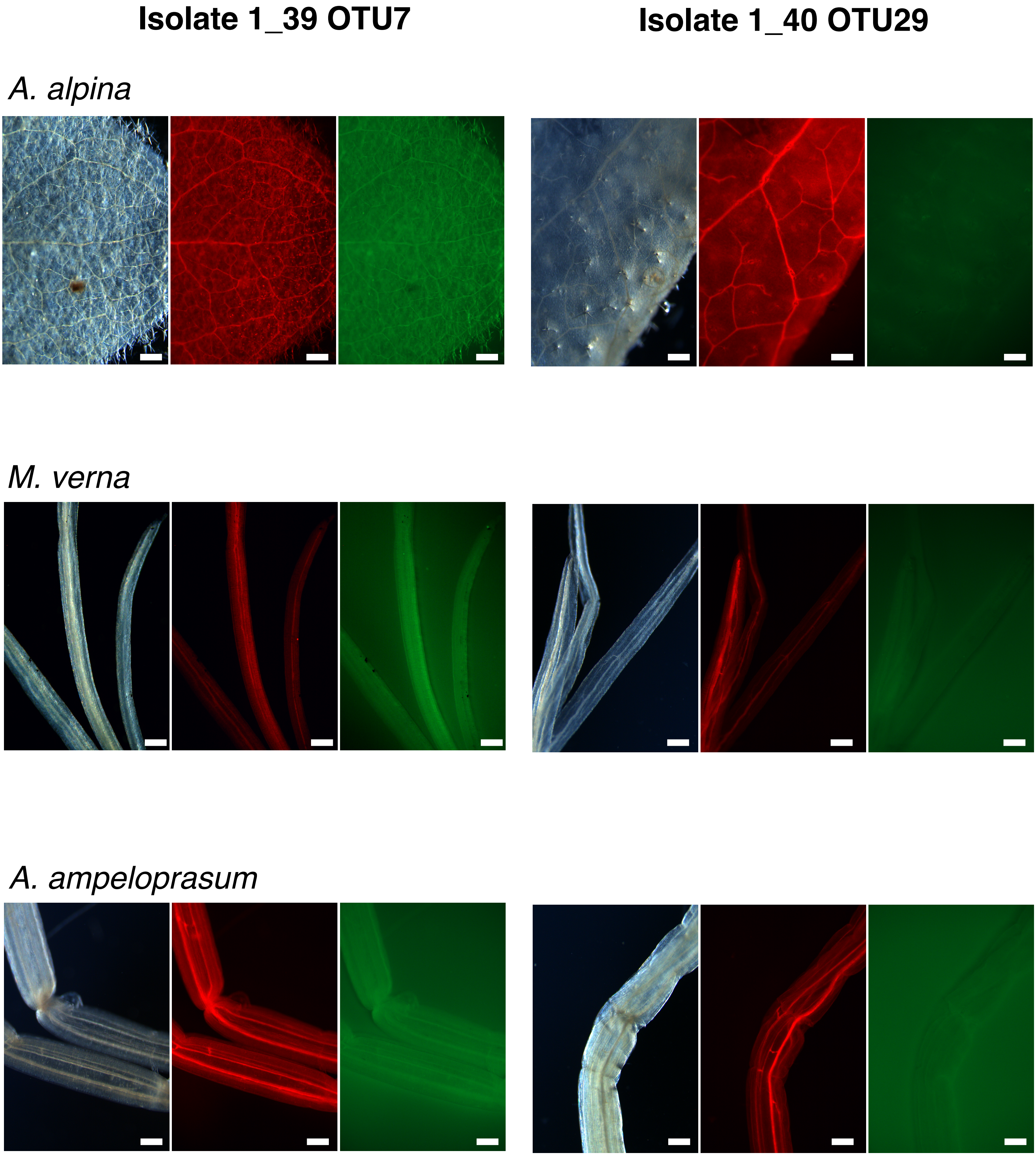
Isolate 1_39 OTU7
Isolate 1_40 OTU29
A. alpina
M. verna
A. ampeloprasum
